# Molecular basis of Siglec-10 ligand recognition and antibody blockade

**DOI:** 10.1101/2025.06.10.658867

**Authors:** Klaudia Sobczak, Asier Antoñana-Vildosola, Pablo Valverde, Eunate Valdaliso-Díez, Amaia Mentxaka, Maria Alejandra Travecedo, Zeinab Jame-Chernaboo, Edward N. Schmidt, Larissa Heinze, Silvia D’Andrea, Iker Oyenarte, Maria Elena Laugieri, Maju Joe, Fahima Mozaneh, Sung-Yao Lin, Mikel Azkargorta, Alexandre Bosch, Maria Jesús Moure, Antonio Franconetti, So Young Lee, Jone Etxaniz-Díaz de Durana, Lorena Pérez-Gutierrez, Asís Palazón, Filipa Marcelo, Elisa Fadda, Francisco Corzana, Ana Gimeno, Felix Elortza, Luciano A. Masullo, Ralf Jungmann, Matthew S. Macauley, Jesús Jiménez-Barbero, June Ereño-Orbea

## Abstract

Siglec-10 is a sialic acid-binding immunoglobulin-like lectin implicated in immune regulation, yet the molecular basis for ligand recognition and how this is functionally linked to immune modulation remains poorly defined. Herein, we present a multidisciplinary study encompassing structural, biochemical, and cellular approaches to elucidate Siglec-10-carbohydrate interactions and their functional consequences. The crystal structure of the extracellular domain of Siglec-10 in complex with α2-6 sialyllactose revealed the presence of two key arginine residues within the Siglec-10 binding site that interact with the carboxyl group of sialic acid, the canonical R119 and R127, suggesting potential dual contributions to ligand engagement. Saturation Transfer Difference (STD)-Nuclear Magnetic Resonance (NMR) confirmed that R119 is essential for sialoglycan binding in solution, whereas R127 appears dispensable for interactions with glycans under these conditions. In contrast, cell-based binding assays using primary human T cells and engineered monocytic lines demonstrated that both arginine residues (R119 and R127) are critical for cellular recognition, revealing a context-dependent interaction. By obtaining direct images at a molecular resolution of 6-7 nm, super-resolution microscopy further revealed glycan-independent dimerization of the Siglec-10 receptor on the surface of human monocytes. Ligand blockade mediated by anti-Siglec-10 mAb (clone S10A) restores CAR-T cell cytotoxicity *in vitro*, supporting its role as an immune checkpoint receptor. Finally, although CD24 was not identified as a Siglec-10 ligand on T cells, proximity labeling and mass spectrometry uncovered other sialylated glycoproteins that may mediate this interaction. Together, these results identify Siglec-10 as a modulatory receptor with structural and functional features distinct from other Siglec family members and highlight its potential for therapeutic targeting in cancer immunotherapy.

## INTRODUCTION

Aberrant glycosylation is a hallmark of cancer. A prominent example is hypersialylation characterized by an increase of sialic acid in cancer, which profoundly alters tumor cell interactions with their microenvironment1,2,3,4. In fact, the interactions of sialic acids with the sialic acid-binding immunoglobulin (Ig)-like lectins (Siglecs) family of immune receptors play a pivotal role in immune surveillance within the tumor microenvironment (TME)5,6,7. The Siglec family of receptors gathers 14 members, broadly divided into two sub-families based on their conservation across species8,9. The therapeutic potential of the Siglec-sialic acid axis within the TME has been highlighted as a novel immune checkpoint that drives innate and adaptive anti-tumor immune responses10,11. Indeed, the inhibitory Siglec receptors8,9 display structural and signaling motifs similar to those of the well-known inhibitory programmed cell death protein-1 (PD-1)12,13. These receptors contain immunoreceptor tyrosine-based inhibitory motif (ITIM) and ITIM-like sequences that can become phosphorylated and recruit SHP1 and SHP2 phosphatases, thereby dampening immune cell signaling14.

Siglec-10 has recently emerged as an immune checkpoint receptor expressed on tumor-associated macrophages (TAMs)15. In the tumor microenvironment, Siglec-10 has been reported to recognize heavily *O*-glycosylated, GPI-anchored cell-surface proteins such as CD24 and CD52, which function as “don’t-eat-me” signals to protect cancer cells from immune clearance15,16,17,18. Despite these emerging links to tumor immune evasion, the molecular mechanisms underlying Siglec-10 engagement with these ligands remain poorly defined. Different monoclonal antibodies (mAbs) that block the interactions of Siglec-10 and its binding partner(s) promote anti-tumor responses and are currently under evaluation for the treatments of several types of cancer15,19. As leading example, ONC-841 (from OncoC4) is currently in Phase 1 Clinical Trial for the treatment of advanced metastatic solid tumors20.

The extracellular domain of Siglec-10 comprises one N-terminal variable (V)-immunoglobulin (Ig) domain and four constant (C)-Ig domains11. Siglecs contain a conserved canonical arginine residue in the V-set Ig domain (R119 in Siglec-10) on β-strand F, which typically forms a salt bridge between its guanidinium group and the carboxyl group of *N*-acetylneuraminic acid (Neu5Ac) 10. The V-Ig domain of Siglec-10 recognizes sialylated ɑ2-3 and ɑ2-6 linked glycans 10,21. Similar to Siglecs-2, -3, -5, -7, -9, and -1522, binding of Siglec-10 to sulfated structures has also been reported using glycan microarrays23. The binding constants of Siglecs, including Siglec-10, for Neu5Ac attached to Gal/GalNAc moieties through ɑ2-3- or ɑ2-6-linkages are rather weak (*K*_D_ ranging from 0.1-3 mM) at the monovalent level as shown by nuclear magnetic resonance (NMR) experiments24,25. Due to the intrinsic nature of these interactions, Siglec oligomerization and clustering are crucial in order to enable multivalent engagement, thus increased avidity providing strong and stable signaling26,27. For Siglec-2 (CD22), monomers have been shown to cluster with their glycan-binding partners on the B-cell surface (*cis*-binding), thereby establishing a signaling threshold that must be surpassed by antigen engagement before B-cell activation 28,29,30.

Herein, we determined the first X-ray crystal structure of the extracellular domain of Siglec-10 in complex with the canonical ligand, α2-6-sialyllactose. Interestingly, we identified a second arginine residue (R127), located proximal to the canonical R119, which also engages the carboxyl group of Neu5Ac. By introducing the corresponding point mutations (R119A and R127A) and assessing their binding properties, we disentangled the respective roles of these two arginine residues in Siglec-10 interactions with a variety of sialoglycans, as measured by NMR, as well as in binding to human T cells, as assessed by flow cytometry.

To determine how mutations in the canonical and non-canonical Arg residues affect Siglec-10 organization on human macrophages, we used super-resolution fluorescence microscopy to quantify receptor distribution and nanoscale clustering. Recent structural work has suggested that Siglec-10 extracellular domains can form homodimers mediated by a hydrophobic interface in domain 2, highlighting a potential role for receptor multimerization in ligand recognition and signaling31. Specifically, we employed DNA-Points Accumulation for Imaging in Nanoscale Topography (DNA-PAINT), a form of single-molecule localization microscopy (SMLM)32,33 in which transient binding of fluorescent “imager” strands to complementary DNA “docking” strands enables precise localization of individual molecules34. Because this blinking mechanism arises from reversible DNA hybridization rather than photophysical switching, DNA-PAINT permits extended imaging with minimal photobleaching and achieves near–single-nanometer or even Ångstrom resolution localization precision35,36,37. This approach allowed us to directly visualize individual Siglec-10 molecules and assess their nanoscale dimerization behavior.

In addition, we have delineated the binding epitope on the Siglec-10 V-Ig domain that is recognized by the fragment antigen binding (Fab) of the anti-Siglec-10 blocking Ab (S10A clone)38. Binding experiments with recombinant Siglec-10 have confirmed the blocking capacity of S10A Ab on human T cells, while cytotoxicity coculture assays have revealed that engagement of cell surface Siglec-10 by S10A Ab has the capability to restore CAR-T cytotoxicity. Overall, these results provide detailed structural and functional insights into the molecular recognition features of Siglec-10 across multiple levels of biological complexity.

## RESULTS

### Three-dimensional structure of Siglec-10 determined by X-ray crystallography

The extracellular domain of Siglec-10 containing the first two N-terminal Ig domains (Siglec-10_d1d2_), was crystallized in complex with the Fab portion of an anti-Siglec-10 mAb (S10A) 38. Consistent with the previous characterization of this IgG antibody 38, the affinity of the S10A Fab was measured to be in the low-nanomolar range, with a *K*_D_ of 1.1 ± 0.1 nM (**Figure S1**). To facilitate the crystallization of the Siglec-10_d1d2_-Fab complex, we employed a crystallization chaperone that binds the human kappa light chain (LC) of the Fab 39. The structure of the ternary complex was solved from a crystal that diffracted to 2.5 Å resolution and processed in C2221 space group (**Table S1**), with two molecules in the asymmetric unit (**Figure S2**). The V-Ig (d1) domain is composed of two antiparallel β-sheets formed by β-strands AA’BED and C’CFG, which are connected by a C41-C101 disulfide linkage, while the second Ig domain (d2) adopts a C1-type fold with strands ABED and CFG (**Figure 1a and Figure S3a**). After endoglycosidase H treatment, the remaining GlcNAc moiety at the N100-linked glycan is located within a hydrophobic patch of the V domain (**Figure S3b**), burying 126.9 Å² of surface area. This N-linked glycan is conserved among Siglecs-2, -4, -5, and -7 29,40,41,42 (**Figure S3c**). To assess the functional relevance of the N100 glycan in Siglec-10, we analyzed its contribution to protein folding and surface presentation in human monocytes by flow cytometry (**Figure S3d**). Notably, whereas the homologous N-glycan is critical for proper folding and surface presentation of other Siglecs, such as Siglec-2 (CD22), the absence of the N100 glycan in Siglec-10 does not impair receptor expression at the cell surface, underscoring functional divergence among Siglec family members.

**Figure 1.**
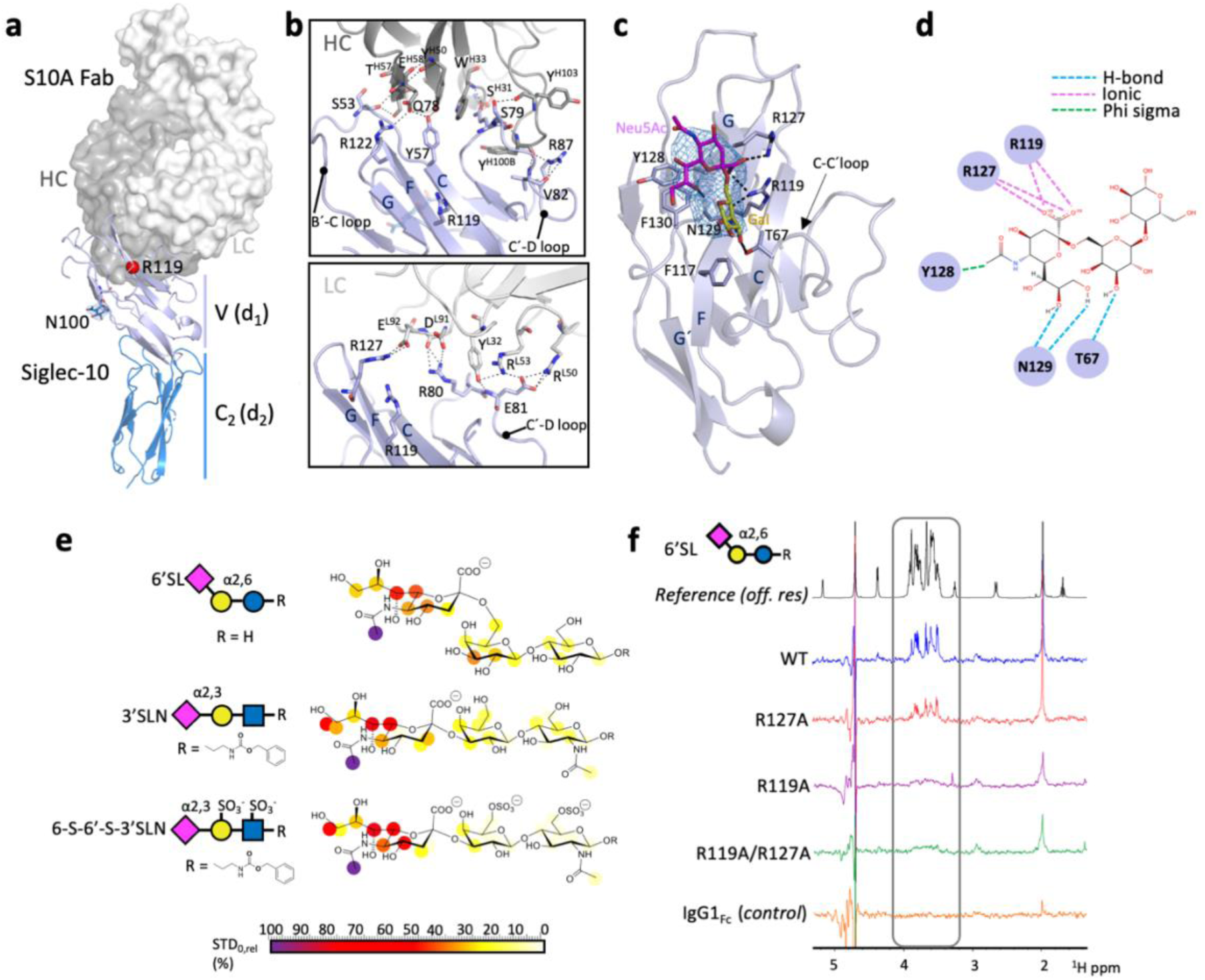
Overall structure of human Siglec-10 in complex with S10A Fab and 6’SL, and its’ molecular recognition patterns with 6′SL, 3′SL and 6-*S*-6’-*S*-3’SLN. a) Surface and cartoon representation of Siglec-10 in complex with anti-Siglec-10 S10A Fab. S10A is composed of a heavy chain (HC) (in dark grey) and a light chain (LC) (in light grey). S10A Fab binds to the V-Ig domain of Siglec-10 (in purple). N100-linked NAG glycan is depicted as sticks with the blue mesh representing the composite electron density map (1.0 σ contour level). b) Amino acid residues involved in interactions between S10A HC (top) and S10A LC (bottom) with Siglec-10. c) Cartoon representation of Siglec-10 V-Ig domain (in purple) with 6’SL represented as sticks with Neu5Ac in pink and Gal in yellow, the Glc moiety is missing due to lack of direct contact with Siglec-10. A blue mesh represents the composite omit electron density map (1.0 σ contour level) associated with Neu5Ac and Gal sugar moieties. Highlighted are the residues involved in polar interactions between Siglec-10 and 6’SL. d) 2D interaction diagram highlighting the key intermolecular contacts responsible for the interactions between Siglec-10 and the 6’SL. e) STD-NMR based epitope mapping for 6′SL, 3′SL and 6-*S*-6’-*S*-3’SLN in the presence of Siglec-10. The relative STD response is coded according to the legend. e) STD-NMR spectra of WT and mutant forms of Siglec-10Fc binding to 6’SL with a molar ratio of 1:100 (Siglec-10: ligand). f) Reference off-resonance spectrum in black, Siglec-10Fc WT in blue, Siglec-10Fc R127A in red, Siglec-10Fc R119A in pink, Siglec-10Fc R119/127A in green and the IgG1Fc negative control in orange. The sialic acid region gated in grey.

S10A Fab binds primarily at the interface of the two β-sheets of the V-Ig domain (**Figure 1a**) (covering a 886.5 Å^2^ buried surface area (BSA)), with the light chain (LC) mainly contributing (BSA= 577.1 Å^2^), as compared to the heavy chain (HC) (BSA = 309.4 Å^2^) (**Figure 1b**). All three of the S10A HC complementarity-determining regions (HCDRs) are involved in the interaction with Siglec-10, providing polar contacts with strand C and B’-C and C’-D loops. Alternatively, the light-chain CDRs 1 (LCDR1) and 3 (LCDR3) interact mainly with strand G and the Ć-D loop of Siglec-10 (**Figure 1b**). Based on this structure, we can infer that S10A might sterically hinder ligand binding.

### Identification of two arginines that mediate Siglec-10–sialic acid binding

To elucidate the binding mode of Siglec-10 to sialoglycans, we attempted to determine complex crystal structures of Siglec-10 with the natural ligands α2-3 and α2-6 sialyllactose. Diffracting crystals were obtained only when α2-6 sialyllactose ([Neu5Acα(2-6)Galβ(1-4)Glc], 6′SL) was soaked into pre-formed crystals of Siglec-10_d1-d2_–S10A Fab. Siglec-10 structure in complex with 6’SL ligand was solved at 3.3 Å resolution (**Table S1**). Fittingly, one of the Siglec-10 molecules of the asymmetric unit was found to be bound to 6’SL (**Figure S2b**). The ligand binding site is formed by strands F and G and loop C–C′ (**Figure 1c**). Overall, the complex and unliganded structures of Siglec-10 V-Ig domain are highly similar (Cα r.m.s.d. of 0.36 Å for d1), indicating that carbohydrate recognition by Siglec-10 is largely mediated by a preformed binding site (**Figure S4a**). No electron density was observed for the glucose (Glc) moiety in our crystal structure (**Figure 1c**). This is likely due to a lack of stabilizing interactions of this moiety with Siglec-10. Most of the interactions occur through the Neu5Ac portion of the ligand (96.5 Å^2^ of BSA for Neu5Ac out of a total of 243.8 Å^2^ for Neu5Acα(2-6)Gal) (**Table S2**). As for other Siglecs, the electrostatic bond established by the highly conserved arginine residue (R119) and the anionic carboxylate group of the Neu5Ac moiety is clearly observed (**Figure 1d**) 43,26. However, an additional arginine residue (R127) was also observed at the β-strand G, which is not found in other Siglecs (**Figure S4b**). Remarkably, the negatively charged C1 carboxylate of the Neu5Ac moiety also interacts with the guanidinium group of this R127 (**Figure 1d**), as well as with that of R119. Additional interactions include N129, which generates hydrogen bonds with the oxygen (O8 and O9) atoms of the Neu5Ac and T67, which establishes another hydrogen bond with the oxygen (O3) of the Gal moiety (**Figure 1d**).

These observations are consistent with very recently reported structural analyses of Siglec-10–sialyllactose complexes, which revealed distinct ligand conformations in α2-3 and α2-6 binding modes and suggested differences in glycan engagement44,31. Motivated by these findings, we further investigated the functional contribution of the non-canonical arginine residue R127 to Siglec-10 glycan recognition.

### Additional insights into the binding mode of Siglec-10 to sialoglycans determined by NMR spectroscopy

To characterize the binding of sialoglycans to Siglec-10, we performed STD-NMR experiments with recombinant Siglec-10_d1-d2_ WT fused to human fragment crystallizable (Fc) from IgG1 (Siglec-10Fc), and 6’-Sialyllactose (6’SL), 3’-Sialyl-*N*-acetyllactosamine (3’SLN), and the sulfated molecule 6-*SO_3_*-6’-*SO_3_*-3’SLN (6-*S*-6’-*S-*3’SLN). The analysis of the STD-NMR profiles revealed that all three ligands exhibit nearly identical binding epitopes, and similar to that described for other related Siglecs24,45 (**Figure 1e**). The *N*-acetyl group of the Neu5Ac residue exhibits the strongest saturation transfer intensity, closely followed by H6 and H7 of the glycerol side chain, strongly suggesting that this region of the Neu5Ac ring is in closer contact with the protein surface, fully consistent with the co-crystal structure. Overall, the sialic acid moiety displays high to moderate relative STD effects, while those found for the adjacent lactose/*N*-acetyllactosamine units are markedly weaker. Only a few weak-moderate relative STDs (20-40%) are detected for the Gal unit, suggesting limited additional contacts with the protein, while the Glc/GlcNAc unit is expected to be mostly solvent exposed. Collectively, these data demonstrate that Siglec-10 specifically recognize both ɑ2-3- and ɑ2-6-sialoglycans and that the binding event in solution is not significantly altered by carbohydrate sulfation.

To further investigate the role of R119 and R127 in binding to sialylated molecules, we introduced R→A or R→K point mutations in recombinant Siglec-10_d1-d2_. These recombinant proteins were still recognized by the S10A Fab, as revealed by BLI (**Figure S4c**), suggesting that they are correctly folded and functional. We compared the STD-NMR profiles of the natural ligand 6’SL in the presence of the Siglec-10 wild type (WT), the two single mutants R119A, R127A, and the double mutant R119A/127A (**Figure 1f**). No detectable STD signals were observed for either the R119A or R119A/R127A variants (**Figure 1f**), whereas clear STD responses were obtained for the WT and R127A proteins. In contrast, the ligand epitope map obtained with the R127A mutant closely resembled that of the WT protein, although the absolute STD₀ values were reduced by approximately 30% (**Figure 1f**). The same approach was undertaken to evaluate the effect of R127A and R119A/R127A point mutations on the binding of 3’SLN and 6-*S*-6’-*S*-3’SLN to Siglec-10Fc (**Figures S5-S8**). A consistent pattern of STD signals was detected for the WT and R127A proteins for both ligands, whereas no STD responses were observed for the R119A/R127A double mutant. (**Figures S5-S8**). For the R127A mutant, the relative STD profiles were preserved but exhibited globally reduced intensities compared to the WT. Together, these observations indicate that R119 is required for detectable sialoglycan binding by Siglec-10 in solution, whereas substitution of R127 reduces overall binding strength, although without markedly altering the ligand interaction pattern.

To complement the experimental data, we performed 1 µs molecular dynamics (MD) simulations46 for various Siglec-10/glycan complexes (**Figure S9**). The simulations for the complex formed between Siglec-10 WT and 6’SL, the ligand remained associated with the binding site throughout the trajectory, with residue R119 playing a central role in ligand binding. Notably, R127 also interacts with both the carboxylate group of the sialic acid and the O8 hydroxyl group of the Neu5Ac moiety (**Figures S9a and b**). In contrast, MD simulations of the R127A mutant in complex with 6’SL showed that, while the complex remained stable for approximately the initial 350 ns, the ligand eventually dissociated (**Figure S9c**). These results are in full agreement with the experimental observations.

### Siglec-10 binding to T cells and the role of two arginine residues in the binding pocket

The role of R119 and R127 residues in Siglec-10 binding to human CD4⁺ and CD8⁺ T cells was further analyzed. Enzymatic removal of surface N- and *O*-linked glycans reduced Siglec-10 WT binding (**Figure 2a and S10a**), confirming the glycan-dependence of this interaction. Similarly, treatment with neuraminidase A, which removes both α2-3- and α2-6-linked sialic acids, significantly reduced Siglec-10 binding (**Figure 2a and S10a**). Specific removal of α2-3-linked sialic acids using neuraminidase S yielded a significant reduction in binding to the human T cells (**Figure 2a and S10a**). Consistent results were observed when testing Siglec-10Fc WT binding to murine CD4⁺ and CD8⁺ T cells deficient in individual sialyltransferases. Binding was preserved in ST3Gal1⁻/⁻ and ST3Gal4⁻/⁻ T cells, suggesting that either another ST3Gal enzyme creates Siglec-10 ligands or that redundancy exists among the ST3Gal enzymes in producing Siglec-10 ligands on murine T cells. In contrast, Siglec-10Fc binding was markedly reduced in ST6Gal1⁻/⁻ T cells, which lack α2-6-linked sialic acids (**Figure 2b**). These findings indicate a key role for α2-6-sialylated glycoconjugates in mediating Siglec-10 recognition on murine T cells. We also assessed the role of glycan sulfation, a known modulator of Siglec binding22,47, using a panel of U937 monocytic cells engineered to overexpress sulfotransferases carbohydrate sulfotransferase 1 (CHST1) and 2 (CHST2)48. We found that Siglec-10Fc binding was reduced in *N*-acylneuraminate cytidylyltransferase (CMAS)49 knockout U937 cells (**Figure 2c and Figure S10b**). However, unlike other Siglecs22,47, Siglec-10Fc showed no enhanced binding to sulfated glycans (**Figure 2d and Figure S10b**). These data are consistent with the STD-NMR results, indicating a lack of dependence on sulfation for ligand recognition.

**Figure 2.**
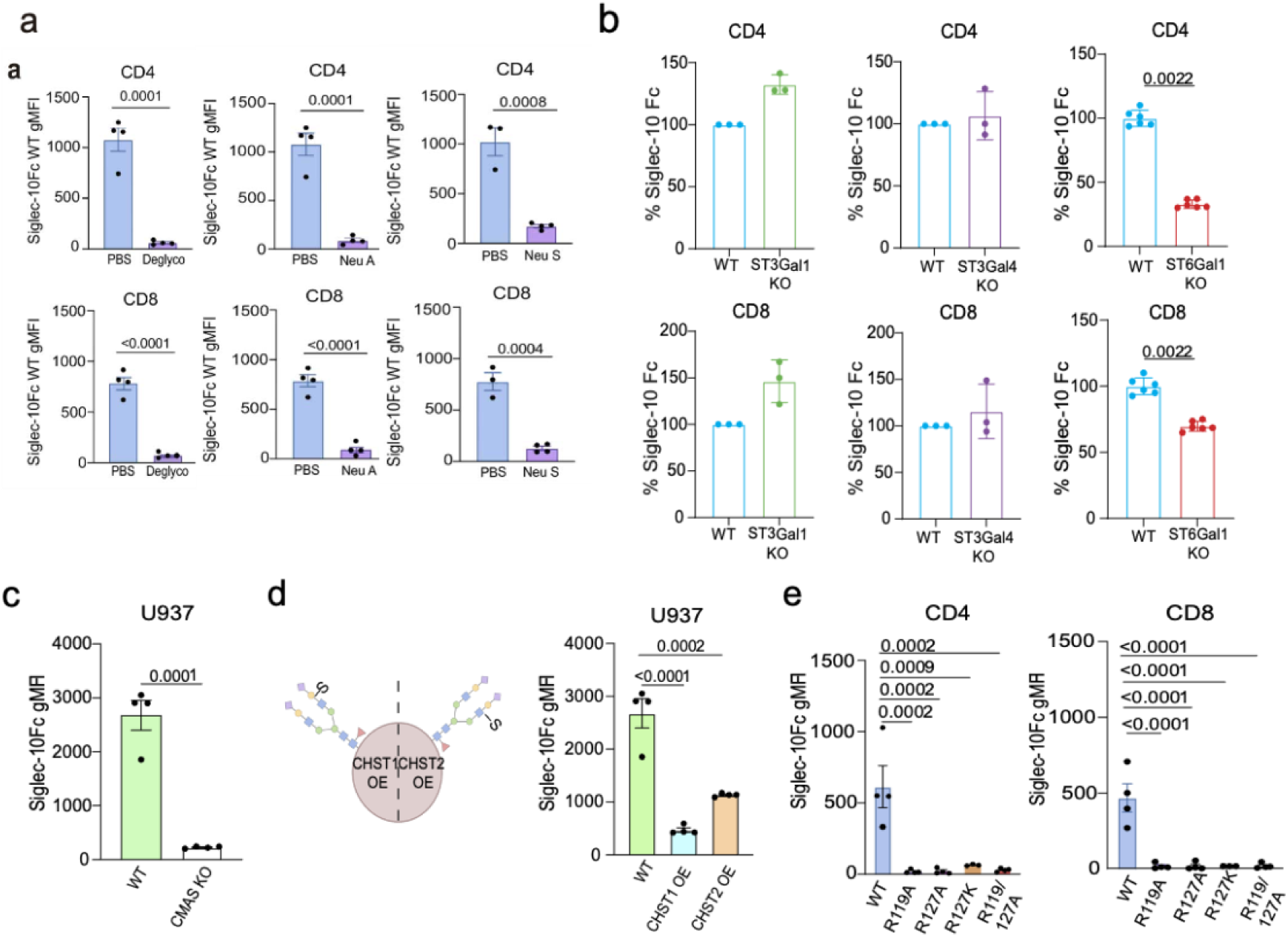
Siglec-10 binding to human cell lines measured by flow cytometry. a) Siglec-10Fc WT binding to deglycosylated, neuraminidase A and neuraminidase S treated CD4+ and CD8+ T cells. b) Pooled data of Siglec-10Fc WT binding to ST3Gal1⁻/⁻, ST3Gal4⁻/⁻ and ST6Gal1⁻/⁻ murine T cells (n=3 donors) c) Pooled data representing the effect of CMAS KO on Siglec-10Fc WT binding (n = 4 donors). d) Right – Cartoon representation of the CHST1 and CHST2 overexpression to increase cell surface sulfation. Left – Siglec-10Fc WT binding (n = 4 donors). e) Siglec-10Fc WT and mutant proteins binding to CD4+ and CD8+ human T cells (n = 4 donors). Error bars denote SEM as determined by two-tailed, unpaired Student’s t test.

To evaluate the individual contributions of the arginine residues to cellular binding, we next examined the binding of Siglec-10 mutants (R119A, R127A, R127K, and the double mutant R119A/R127A) to primary human T cells. Remarkably, all mutations, whether targeting the canonical R119 or the non-canonical R127, completely abolished binding to both CD4⁺ and CD8⁺ T cells (**Figure 2e** and **Figure S10c**), underscoring that both arginine residues are essential for cellular recognition in the cellular context.

Building on these findings, we sought to gain mechanistic basis for engagement of Siglec-10 with multi-sialylated epitopes, such as the disialylated ganglioside GT1b50, at the molecular level. Thus, we performed two independent 500-ns molecular dynamics (MD) simulations for Siglec-10 in complex with GT1b embedded in a lipid bilayer. A full-length model of Siglec-10 was generated using AlphaFold246, which produced a highly accurate model with strong agreement to the experimentally determined V-set domain structure (RMSD = 0.429 Å; **Figure S11**).

Interestingly, in both MD replicas, GT1b remained associated with the protein and formed stable contacts with both arginines. In the first replica, R119 primarily engaged the α2-8-linked Neu5Ac, maintaining >95% occupancy, while R127 predominantly interacted with the terminal α2-3-linked Neu5Ac (77% occupancy). In the second replica, R119 shifted its interaction to the terminal α2-3-linked Neu5Ac, indicating variability in its engagement with differently linked sialic acids. In contrast, R127 consistently maintained interaction with the α2-3 Neu5Ac across both simulations (**Figure S11)**. Together, these simulations support that Siglec-10 utilizes both canonical and non-canonical arginine residues to achieve high affinity and selective recognition of multi-sialylated epitopes, such as the GT1b ganglioside.

### Siglec-10 oligomerization on human macrophages

Taking into account that Siglecs, such as CD2219, can engage in *cis* with sialoglycans and assemble into higher-order oligomers on the same cell surface, we next examined the organizational state of Siglec-10 on human macrophages. Because both arginine residues (R119 and R127), participate directly in sialic acid recognition, we assessed whether these residues are necessary for Siglec-10 oligomerization. To this end, human monocyte cell lines (THP-1) expressing intracellularly GFP-tagged Siglec-10 variants (WT, R119A and R127A) were generated. Cells were seeded onto glass slides, labelled with anti-GFP nanobodies, and imaged using DNA-PAINT at ∼3 nm localization precision (**Figure 3a, c and e**). Single-protein coordinates were then reconstructed from the resulting localization datasets. Then, simulations (n= 50 repeats per set of parameters) with different proportions of monomers and dimers were generated and the best fitting model was obtained (**solid lines in Figure 3b, d and f**).

**Figure 3.**
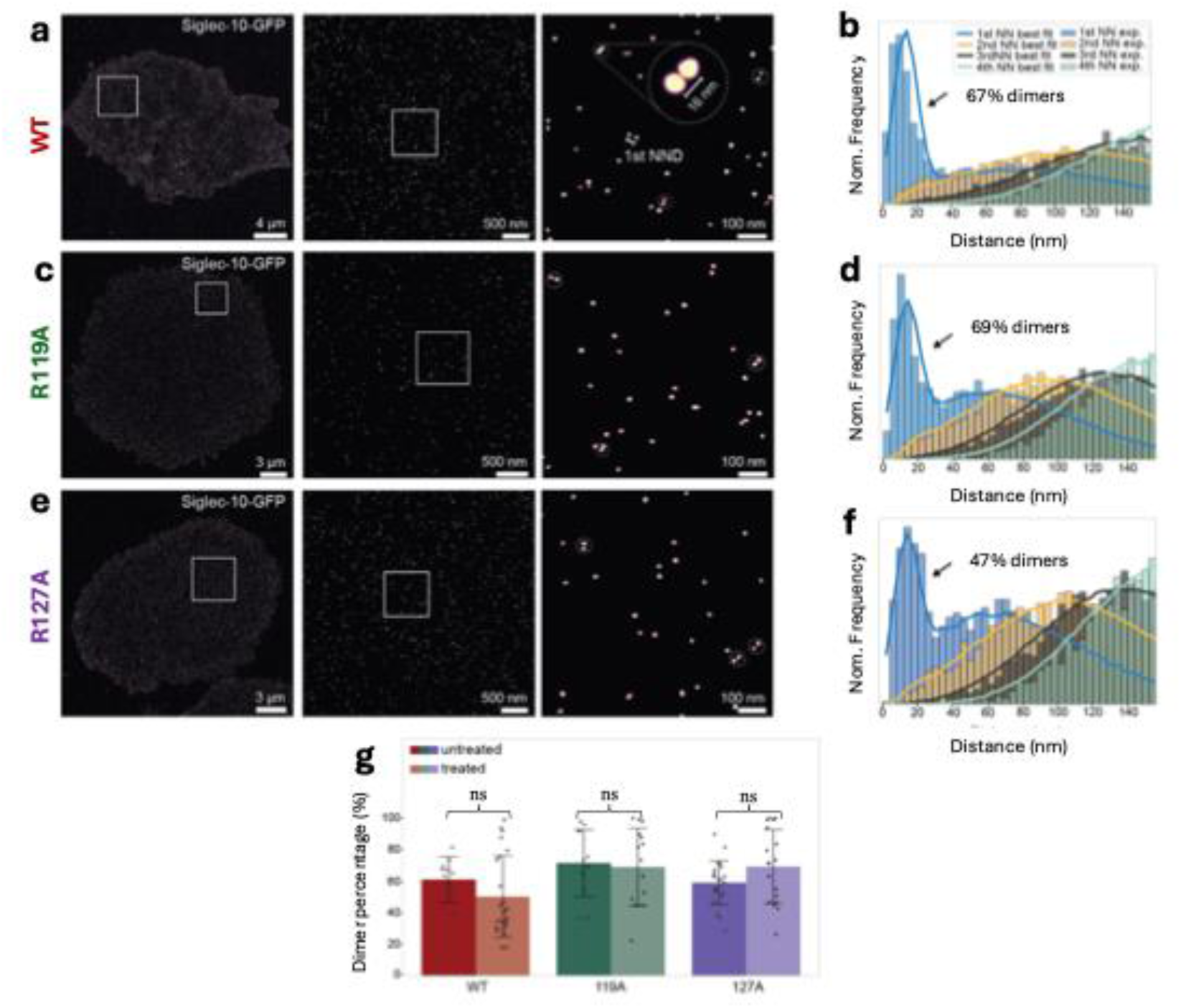
The oligomerization state of Siglec-10. a,c,e) DNA-PAINT images with zoom-ins of intracellularly GFP-tagged Siglec-10 in THP1 wild-type (WT) (a), carrying the R119A mutation (c) or the R127A mutation (e) cells. b,d,f) Corresponding nearest-neighbor distance (NND) histograms with SPINNA-based model fits (solid lines), indicating 67% (b), 69% (d), and 47% (f) dimeric structures, respectively. g) Bar chart showing the mean ± SD of measured dimer percentages before and after tunicamycin treatment. Pre-treatment values: WT, 61 ± 14% (n = 11); R119A, 72 ± 21% (n = 12); R127A, 59 ± 14% (n = 23). Post-treatment values: WT, 50 ± 26% (n = 23); R119A, 69 ± 25% (n = 15); R127A, 69 ± 23% (n = 18). Statistical significance was determined using pairwise Welch’s t-tests, with ns indicating p > 0.05 and * indicating 0.01 < p < 0.05

Despite the point mutations on Siglec-10, oligomerization was detected for all Siglec-10 variants examined. Quantitative analysis revealed 67% of dimer formation for the WT protein (**Figure 3b**), compared to 69% dimers for the R119A mutant (**Figure 3d**) and 47% of dimers for the R127A mutant (**Figure 3f**). These data suggest that homo-oligomerization of Siglec-10 is not affected by its capacity to bind sialoglycans. To further assess the contribution of glycosylation to Siglec-10 organization, we performed the same set of experiments in the presence of tunicamycin, a potent inhibitor of N-linked glycosylation51 (**Figure S12**). No significant differences in oligomerization were observed between untreated *versus* tunicamycin treated cells, further supporting the fact that Siglec-10 dimerization is not sialic acid-mediated (**Figure 3g**).

### Siglec-10 binding on human T cells occurs independently of CD24

CD24 expression on T cells is required for optimal homeostatic proliferation of both CD4 and CD8 T cells52. Thus, we next investigated whether CD24 contributes to Siglec-10 recognition on human T cells, beginning by assessing CD24 expression on peripheral blood–derived T cells. CD24 expression was maintained on non-activated T cells and persisted at 24, 48, and 72 hours post-activation (**Figure 4a and Figure S13a**), in agreement with previous reports53,54,52. We next examined whether CD24 functionally mediates Siglec-10 interactions by performing binding assays in the presence of a CD24-blocking monoclonal antibody (clone SN3)38. Despite robust CD24 expression, blockade with SN3 did not alter Siglec-10 binding to activated T cells (**Figure 4b**). These results were further validated by measuring CD24-overexpressing HEK293T cells treated with phospholipase C, which cleaves GPI-anchored surface proteins. Although surface CD24 levels showed a reduction under these conditions, Siglec-10 binding remained unaffected (**Figure 4c**). Together, these results indicate that CD24 is not a major ligand for Siglec-10 in this context.

**Figure 4.**
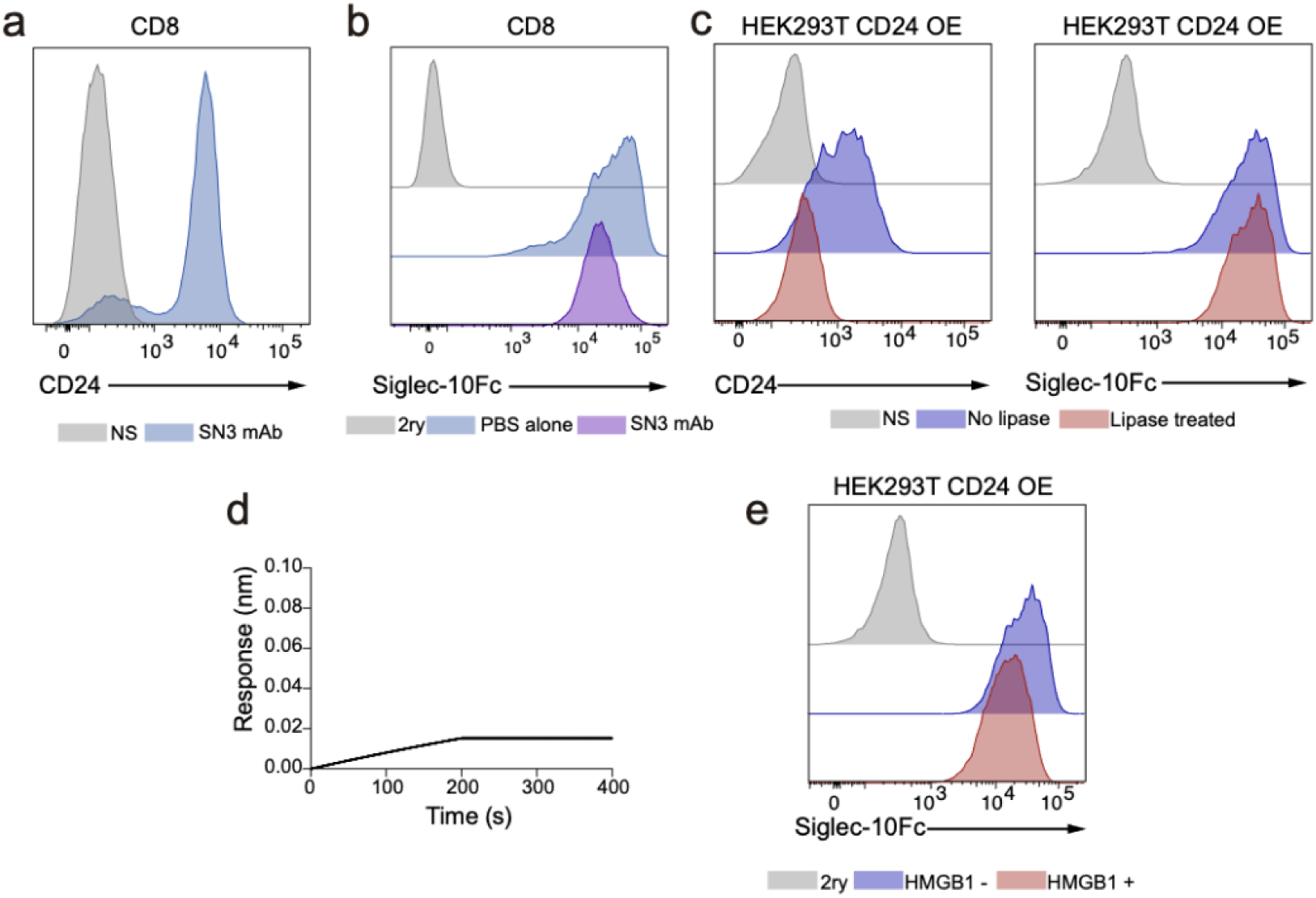
Flow cytometry analysis of CD24 expression and Siglec-10 binding. a) Representative histogram of CD24 expression on activated human T cells. b) Histogram showing Siglec-10 binding to activated T cells with (purple) or without (blue) SN3 anti-CD24 mAb blockade. c) Left – Histogram representing the cleavage of CD24 post phospholipase C (lipase) treatment where non-treated cells are represented in dark blue and treated cells are in red. Right – Representation of Siglec-10 binding to phospholipase C (lipase) treated cells (red) in comparison to non-treated (dark blue). d) BLI sensorgrams for the interaction between CD24 and HMGB1 recombinant proteins. e) Histogram representing Siglec-10 binding to HEK293T CD24 overexpressing (OE) cells in the presence (red) and absence (dark blue) of HMGB1 recombinant protein.

We next assessed whether Siglec-10 and CD24 form a ternary complex with high-mobility group box 1 (HMGB1), a protein reported to negatively regulate their stimulatory activity and inhibit immune responses55. HMGB1–Siglec-10 interactions were examined using CD24-overexpressing HEK293T cell lines (**Figure S13b**) and by biolayer interferometry (BLI) with recombinant proteins. However, no differences in Siglec-10 binding were observed under these conditions (**Figure 4d,e**). Together, these findings suggest that Siglec-10 recognition of human T cells is mediated by ligands distinct from CD24 or HMGB1, prompting us to pursue an unbiased strategy to identify alternative interaction partners.

Thus, proximity labeling experiments were carried out to identify candidate Siglec-10 ligands on activated human T cells from two independent donors. CD24 was not detected as a Siglec-10 interactor in our analysis (**Table S4),** although several other glycoproteins and receptors were identified whose functional relevance remains to be validated. Collectively with the binding and blockade data, these results suggest that Siglec-10 recognition on human T cells occurs independently of CD24, in contrast to the observations reported in murine systems56. These findings point to potential species-specific differences in ligand engagement and underscore the need to define alternative Siglec-10 ligands in human immune regulation.

### Antibody-mediated blockade of Siglec-10 restores CAR-T cell cytotoxicity

To evaluate the impact of blocking Siglec-10 on T cell cytotoxicity, we performed an antigen-specific *in vitro* cytotoxicity assay using THP-1 myeloid cells engineered to express a non-signaling truncated CD19 (CD19t) together with either Siglec-10 WT or ligand-binding-deficient mutants (R119A, R127A, R119/127A) (**Figure 5a** and **Figure S13a**). Primary human T cells were transduced with a validated anti-CD19 CAR construct57 (**Figure S13b**). In this assay, overexpression (OE) of Siglec-10 WT significantly impaired CAR-T cell-mediated killing, whereas THP-1 cells expressing the R119A, R127A, or R119/127A variants exhibited similar inhibitory levels of CAR-T inhibition comparable to those of Siglec-10 WT OE. These results strongly suggest that Siglec-10 suppresses CAR-T activity independently of its ligand-binding capacity (**Figure 5b and c**).

**Figure 5.**
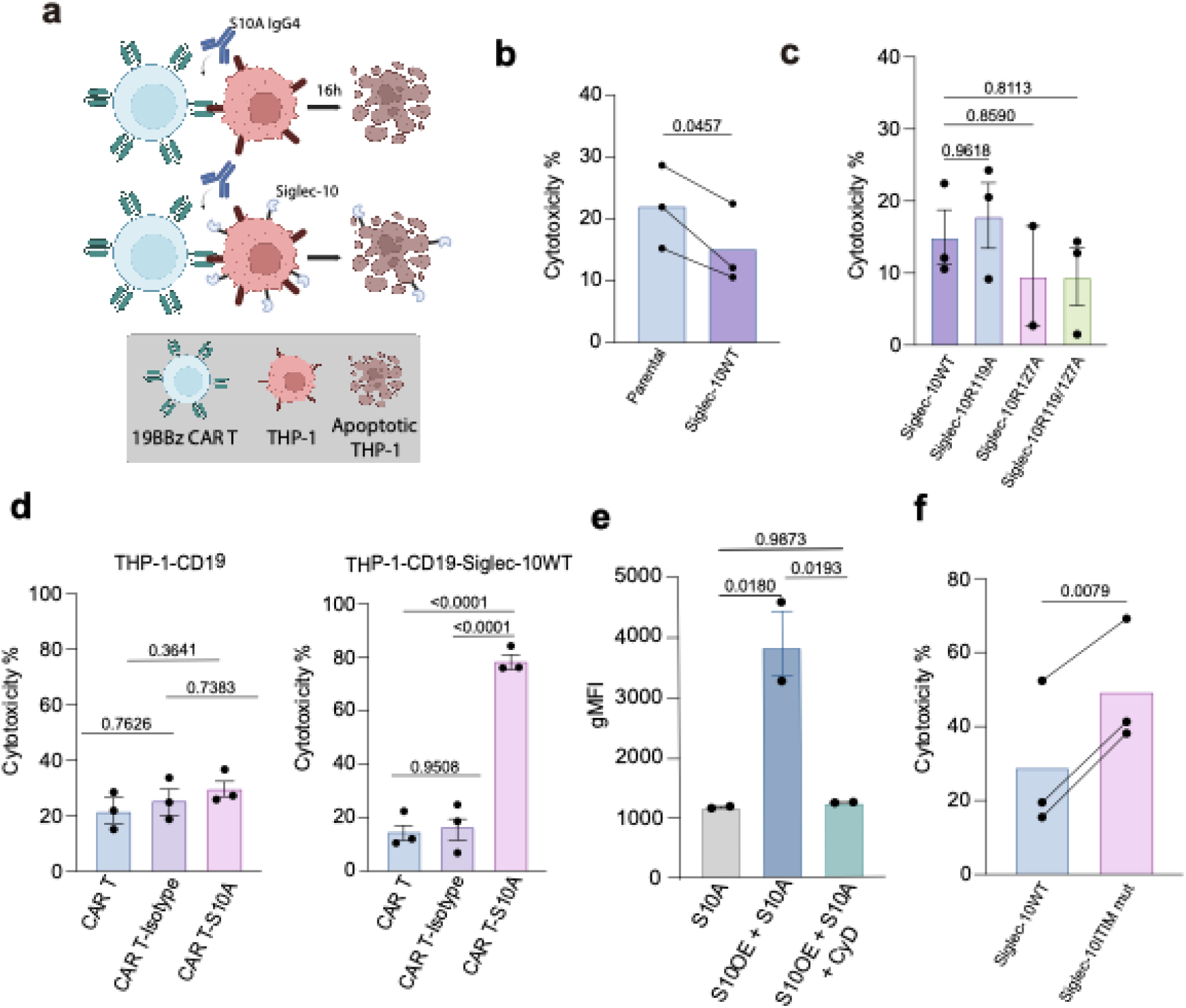
Siglec-10 modulates CAR-T cell cytotoxicity. a) Schematic representation of the *in vitro* cytotoxic assay. b) Pooled data showing the impact of Siglec-10WT expression on CAR-T cell cytotoxicity compared to parental THP-1 cell line (n= 3 donors, statistical significance was determined by paired Student’s t test). c) Pooled data representing the impact of Siglec-10 R119A, R127A and R119/127A on CAR-T cell cytotoxicity in comparison to WT (n=3 donors). d) Pooled data showing the effect of S10A IgG4 mAb treatment on CAR-T cytotoxicity in the presence (left) or absence (right) of Siglec-10 expression (n= 3 donors). e) Pooled quantification of S10A IgG4 internalization in THP-1 cells with and without endocytosis inhibition by CyD, at different time points by flow cytometry (n= 2 donors). f) Pooled data representing the impact of ITIM signaling motif mutation on CAR-T cell function in comparison to Siglec-10WT (n=3 donors, data represented as paired Student’s t test). For all of the above data sets, data represent mean ± SEM; statistical significance was determined by one-way ANOVA, unless stated otherwise.

To determine whether Siglec-10 blockade could restore CAR-T cytotoxicity, cocultures were treated with the S10A IgG4 blocking monoclonal antibody (**Figure S16c**). S10A treatment led to a 4-fold increase in CAR-T-mediated killing compared to control conditions, demonstrating effective reversal of Siglec-10–mediated inhibition (**Figure 5d and S16e**). Analysis with pH-sensitive pHrodo red fluorophore in combination with confocal microscopy showed internalization of Siglec-10 on target cells following S10A engagement, suggesting that antibody-mediated receptor removal contributes to enhanced CAR-T effector function (**Figure 5e and S16f**). Notably, overnight coculture with anti-CD19 CAR-T cells induced Siglec-10 expression on the CAR-T cell surface (**Figure S14d**). Thus, although S10A-mediated enhancement of cytotoxicity is consistent with antibody-induced removal of Siglec-10 from target cells, a contribution from direct modulation of Siglec-10 on CAR-T cells cannot be excluded.

To further explore the mechanism of Siglec-10–mediated suppression, we analysed THP-1 cells expressing signaling-deficient Siglec-10 mutants. CAR-T-mediated cytotoxicity increased from approximately 30% against cells expressing WT Siglec-10 to ∼50% against cells expressing ITIM mutant Siglec-10, suggesting that the intracellular signaling of Siglec-10 partially mediates the inhibitory effect (**Figure 5f**). Together, these findings demonstrate that Siglec-10 limits CAR-T cell cytotoxicity through a combination of both its expression and intracellular signaling, and that antibody-mediated blockage with S10A can effectively restore CAR-T effector function *in vitro*.

## DISCUSSION

Siglec-10 is a member of the Siglec family that modulates immune responses through recognition of sialoglycans. Despite its functional relevance in immune tolerance and tumor immune evasion, the structural and molecular basis of Siglec-10 ligand recognition has remained elusive. Here, we have combined structural biology, NMR spectroscopy, DNA-PAINT, and cellular assays to provide a comprehensive picture of the glycan-binding interface of Siglec-10 and to assess its functional implications and consequences in immune cell recognition.

The analysis of our crystal structure of Siglec-10 in complex with 6’SL has revealed the presence of two Arg residues, R119 and R127, within the carbohydrate recognition domain. To our knowledge the presence of this dual-arginine display has not been previously identified for Siglecs and suggests a distinct mode of sialic acid engagement, with a potential cooperative role of the two arginines. Consistent with this observation, STD-NMR experiments demonstrated that mutation of the canonical R119 residue alone is sufficient to abolish binding to trisaccharide glycan ligands in solution. In contrast, cell-based assays highlighted the importance of the biological context: mutation of either R119 or R127, completely eliminated Siglec-10 binding to human CD4⁺ and CD8⁺ T cells. Together, these findings indicate that although R127 is dispensable for recognition of minimal glycan motifs *in vitro*, it becomes essential for productive glycan engagement in the cellular environment. Additionally, our super-resolution microscopy data have further revealed that Siglec-10 homo-dimerization occurs independently of canonical sialic acid recognition mediated by R119 and R127, suggesting that *cis* engagement with sialoglycans is not the primary driver of its nanoscale organization. Instead, our data support a model in which Siglec-10 clustering is governed by intrinsic protein–protein interactions or membrane-associated factors, rather than glycan-dependent crosslinking. One possible explanation for the role of R127 is that it stabilizes or properly orients the carbohydrate recognition domain for productive interactions with complex or multivalent glycans present at the cell surface, such as disialylated gangliosides. This interpretation is further supported by MD simulations of Siglec-10 bound to the GT1b ganglioside, which underscore the key contribution of R127 for maintaining persistent Siglec-10/glycan interactions under membrane-associated conditions mimicking those physiologically relevant. Notably, reliance on non-canonical residues for recognition of complex sialylated structures has been described for other Siglecs; for instance, Siglec-7 has been shown to rely on both canonical and non-canonical residues to engage di- and tri-sialylated structures58.

The glycan dependence of Siglec-10 binding was further investigated using a combination of enzymatic treatments, genetically modified cell models, and sialyltransferase-deficient mice. Enzymatic removal of N- and *O*-linked glycans from human T cells completely abolished Siglec-10 binding, confirming that the interaction is glycan-dependent. Additionally, treatment with an α2-3-specific neuraminidase also resulted in a marked reduction of binding. In murine T cells, Siglec-10 binding was markedly reduced in ST6Gal1⁻^/^⁻ cells, which lack α2-6-linked sialic acids, while binding was largely preserved in ST3Gal1⁻^/^⁻ and ST3Gal4⁻^/^⁻ cells. Together, these results point out that both α2-3- and α2-6-sialylation are required for Siglec-10 recognition in T cells, although species-specific differences may exist between human and murine systems. These conclusions should nevertheless be interpreted with caution, as the experimental approach used in the two systems differ and display distinct limitations. For instance, α2-6-linked sialic acids can adopt conformations, such as hairpin-like structures, which render them less accessible and may limit the efficiency of enzymatic cleavage. In contrast, terminal α2-3-linked sialic acids are typically solvent-exposed, which may account for the more pronounced effects observed with α2-3-specific neuraminidase treatment in human cells. Moreover, genetic knockout models must be interpreted with caution, as compensatory pathways or broader alterations in the glycome architecture may indirectly influence Siglec-10 binding outcomes in ways that are not immediately apparent.

Further supporting the requirement for sialylated ligands, Siglec-10 binding was substantially reduced in CMAS-deficient U937 cells, which lack *de novo* sialic acid biosynthesis. This observation confirms that sialic acids are essential for Siglec-10 interactions across multiple cellular contexts. Moreover, in contrast to several other Siglec family members, Siglec-10 binding was not enhanced by overexpression of the sulfotransferases CHST1 or CHST2, indicating that sulfation does not play a major role in modulating ligand affinity. This observation is consistent with the STD-NMR data, which revealed minimal engagement with the sulfated glycan epitopes.

In light of previous studies implicating CD24 as a key Siglec-10 ligand in cancer, we investigated whether CD24 mediates Siglec-10 interactions on T cells. Although we confirmed high CD24 expression on T cells, neither antibody-mediated CD24 blockade nor CD24 overexpression altered Siglec-10 binding. These results strongly suggest that CD24 does not play a major role in Siglec-10 recognition in these cellular contexts, in contrast to earlier reports linking CD24 to Siglec-10 signaling within tumor-associated macrophages. Previous studies have also proposed that the high-mobility group box 1 (HMGB1) protein facilitates the formation of a ternary complex involving Siglec-10 and either CD2459 or CD5260. However, in our experimental system, we did not detect any Siglec-10–CD24 interaction even in the presence of HMGB1. Consistent with these findings, a recent CRISPR activation (CRISPRa) screen in K-562 cells identified N-linked glycans terminating in α2-3- or α2-6-linked sialic acids as preferred Siglec-10 ligands61,62. Notably, CD24 was not among the candidate ligands identified in this screen. In our study, proximity labelling coupled with mass spectrometry likewise failed to detect CD24 among the candidate counter-receptors recognized by Siglec-10 on T cells. These findings collectively support the notion that Siglec-10 interacts with a broader repertoire of sialylated glycoproteins in a context-dependent manner. It remains conceivable that CD24 contributes to Siglec-10 engagement only in specific immunological environments or under distinct post-translational modifications, such as particular fucosylation patterns or clustered *O*-glycosylation motifs, which may not be recapitulated in the systems examined here.

Taken together, our results highlight the complex interplay between structural determinants and cellular context for Siglec-10/ligand recognition. The dual-arginine display in the CRD may enable Siglec-10 to engage structurally diverse sialylated glycan ligands across different cellular environments. The lack of sulfation dependence and the lack of evidence for CD24 as a universal ligand further suggest that Siglec-10 likely recognizes distinct, yet-to-be-identified glycoproteins and/or glycolipids on T cells. Identifying and functionally validating these ligands on T cells will be essential for fully identifying the endogenous Siglec-10 interactome and their relevance in immune regulation in both health and disease conditions.

In this context, our previous work on Siglec-1524, together with a recent study by Saini et al.63, has demonstrated that integrins can function as Siglec ligands. Notably, interactions between Siglec-10 and α3β1 integrin have been shown to enhance macrophage-mediated phagocytosis in pancreatic cancer. Building on these findings, an important next step will be to determine whether additional putative Siglec-10 ligands, such as the integrins, also contribute to the regulation of T-cell effector function.

Overall, our findings define how Siglec-10 integrates glycan recognition with receptor-intrinsic inhibitory signaling to restrain T cell effector function in a context-dependent manner. The data support a role for Siglec-10 as a glyco-immune checkpoint receptor capable of modulating T cell effector function and demonstrate that this suppression can be alleviated through Siglec-10 blockade. By linking ligand engagement, receptor nanoscale organization, and ITIM-dependent signaling, our findings provide mechanistic insight into Siglec-10 function, not only reinforcing Siglec-10 as a *bona fide* immune checkpoint, but also providing a strong rationale for its therapeutic targeting to enhance T cell–based immunotherapies, including CAR-T cell strategies.

## EXPERIMENTAL SECTION

### Primary human T cells

T cells were obtained from buffy coats of healthy donors (Biobanco Vasco, BIOEF) after ethical approval (PI + CES-BIOEF 2019-08). Briefly, PBMCs were separated by gradient differentiation using FicollHistopaque (17-1440-03, Fisher scientific), and CD3+ T cells were purified by negative selection using EasySep™ Human T Cell Enrichment Kit (Stemcell) following manufacturer’s instructions. Purity was confirmed by flow cytometry to be >95%. T cells were then activated with anti-CD3/CD28 Dynabeads (11131D, Thermo Fisher) in CST OpTimizer medium (A1048501, Gibco) supplemented with IL-2 at 100 IU/mL (130-097-743, Miltenyi Biotec).

### CAR constructs

Second generation CARs targeting CD19 were developed in our laboratory. The anti-CD19 scFv was derived from the FMC63 monoclonal antibody (REF Nicholson Mol. Immunol. 34, 1157–1165 (1997)). The CAR construct included CD8a signaling peptide, the anti-CD19 scFv, the CD8alpha hinge/transmembrane domain, 4-1BB intracellular domain, and CD3z intracellular domain. To facilitate CAR-T cell identification and selection, a T2A self-cleaving peptide followed by truncated NGFR was incorporated into the CAR coding sequence. The construct was cloned into a third-generation lentiviral vector under the control of the EF1a constitutive promoter and synthesized by Genscript.

### Lentivirus production and titration

To generate lentiviral particles, 5 × 10^6^ 293T cells were seeded in a 100 mm dish. Twenty-four hours after cell seeding, cells were transfected using a mix of lentiviral plasmids (three plasmids were used for 2^nd^ generation lentivirus used for THP-1 cell line transduction: transfer plasmid; 5 µg, psPAX2 (Addgene #12260; 4 µg) and VSV-G (Addgene #8454; 1.5 µg. In the case of 3^rd^ generation lentivirus for transduction of primary T cells, four plasmids were used: transfer plasmid; 6.9 µg, RRE 3.4 µg, REV 1.7 µg and VSV 2.0 µg)) and jetPEI transfection reagent (Polyplus) following manufacturer’s instructions. After 48 h, lentiviral particles were harvested from the supernatant, filtered through a 0.45 µm filter (VWR) and concentrated using LentiX concentrator (Takara) at a 3:1 ratio at 4 °C overnight. Lentiviral particles were then concentrated by centrifugation at 1500g for 45 min, aliquoted, and stored at −80 °C until use.

The titration of the lentivirus was performed using Jurkat cells. Briefly, 5×10^4^ Jurkat cells were plated in 96-well plates in complete RPMI medium containing serial dilutions of thawed lentivirus (e.g. 3-fold serial dilutions from 3:100 to 3:100,000) and 8 mg/ml polybrene. After 48 hours, cells were analyzed for transgene expression by flow cytometry. Virus titer was calculated according to the following formula: Titer (pfu/ml) = [(5×10^4^) * (%positive cells) / 100] / [Volume of virus added (ml)].

### Human T cell transduction

For CAR lentiviral transduction, CD3^+^ T cells were initially activated with CD3/CD28 Dynabeads in OpTmizer medium supplemented with human IL-2. Two days post-activation, these cells were transduced with the desired lentivirus at a multiplicity of infection (MOI) of 6 in the presence of 8 µg/mL Polybrene (Merck). Spinoculation was performed at 800g for 1 hour at 32°C and cells were maintained in complete OpTmizer media containing IL-2 at a density of 0.5×10^6^ cells/mL, with fresh media replenished every 2-3 days. Four days after transduction, beads were magnetically removed, and CAR-T cells were isolated using the EasySep Human CD271 Positive Selection Kit II (StemCell Technologies). These selected cells were expanded until days 10–12 post-activation, then cryopreserved in FBS with 10% DMSO. The purity of the CAR-T cells was routinely verified to be >85% by flow cytometry.

### Cell lines

HEK293F (R79007, Thermo Fisher) and HEK293S (CRL-3022) cells were grown at 37°C, 70% humidity 8% CO2 at 130 rpm in Freestyle media (12338018, Thermo Fisher). Jurkat, THP-1 (WT and THP-1-CD19t-GFP transduced) (TIB-202, ATCC) and U937 (the U937-CHST1, U937-CHST2 and U937-CMAS KO cell lines were generated according to a previously published protocols47,64) cells were grown in RPMI (15140122, Gibco) medium. The HEK293T and HEK293T Lenti-X (632180, Takara Bio Inc.) cell lines were maintained in DMEM (41966-029, Gibco). Both, RPMI and DMEM media were supplemented with 10% heat-inactivated FBS (A3840002, ThermoFisher) and 1% penicillin/streptomycin mix (pen/strep) (15140122, ThermoFisher). THP-1 cell line was grown in T25 flasks (156340, ThermoFisher), whereas HEK293T LentiX were maintained in T75 flasks (156472, ThermoFisher), and passed twice a week. To keep optimal conditions for cell culture, the THP-1 and HEK293T LentiX cells were kept at 37°C, 5% CO2, and 70% humidity.

For cytotoxicity experiments, THP-1 cell line was transduced with a lentivirus expressing luciferase-EGFP and CD19 target antigen followed by T2A-puromycin resistance gene and was then sorted for CD19^+^GFP^+^ cells. Following, the THP-1-GFP-CD19 cell line was transduced with various Siglec-10 constructs, specifically Siglec-10 WT, R119A, R127A, R119/127A and ITIM mut followed by a T2A-blasticidin resistance gene, generating five different Siglec-10 expressing THP-1-GFP-CD19 cell lines.

For super-resolution microscopy, THP-1 cell line was transduced with a lentivirus expressing various constructs of Siglec-10, specifically Siglec-10 WT, R119A, R127A mut and luciferase-EGFP, followed by a T2A-blasticidin resistance gene, generating three different THP-1-Siglec-10-GFP cell lines.

For CD24 experiments, HEK293T cells were transduced with GPI anchored CD24. The construct was designed according to a previously published study to ensure GPI anchoring, correct protein folding and functionality65. This was followed by T2A-blasticidin resistance gene for selection purposes.

### Mouse Strains

All mice used in this study were on a C57BL/6J genetic background. The following strains were obtained from The Jackson Laboratory: C57BL/6J (Strain #: 000664), CD4^Cre^ (Strain #: 022071), ST6Gal1^fl/fl^ (Strain #: 006901). To generate the ST6Gal1^fl/fl^ × CD4^Cre^ mice, ST6Gal1^fl/fl^ animals were crossed with CD4^Cre^ mice. Genotyping was performed by PCR using DNA extracted from ear notch tissue. For the ST6Gal1 locus, the forward primer 5’-TCAGCAGCCTGGGGTTAAGT-3’ and reverse primer 5’-CCTGTACACTGAATGGTGGACTGTGG-3’ were used. The wild-type allele generated a 1115 bp product, while the floxed allele produced a 900 bp product. ST3Gal1 and ST3Gal4 knockout mice were generously provided by Dr. Jamey Marth (Sanford Burnham Prebys Medical Discovery Institute, USA).

### Construct design of Siglec-10 extracellular domain and S10A proteins

The DNA encoding the first Ig domains of the extracellular domain (d1-d2) of human Siglec-10 (UniprotKB Q96LC7, residues 17-238) fused to 6xHis tag, codon optimized for expression in human cells and cloned into pHLsec vector between and KpnI restriction sites. The DNA encoding human Siglec-10_d1–d2_ WT and R119A, R127A, R127K and R119A/R127A mutants fused to the human IgG1 Fc region (UniprotKB P01857, residues 99–330) and with a C-terminal 6x His tag, were subcloned between XbaI and AfeI restriction sites into pcDNA 3.4 (Invitrogen) and codon optimized for expression in human cells. For the S10A Fab, the heavy (residues 1-214) and light (residues 1–224) chains were synthesized and cloned into pHLsec vector between and KpnI restriction sites. For the S10A IgG4 mAb, the light chain used was the same as for S10A Fab, as described above, whereas the heavy chain was fused with the IgG4 backbone (residues 1-327) and was synthesized and cloned into pcDNA3.1 vector between Xbal and AgeI restriction sites. All plasmids were synthesized by GenScript.

### Protein expression and purification

Siglec-10_d1-d2_ and Siglec-10Fc (WT and mutants) constructs were transiently transfected using HEK293F/S suspension cells. Cells were split in 500 mL cultures at 1.0 × 10^6^ cells/mL. The DNA: FectoPRO transfection reagent solution (101000007, Polyplus) was then added directly to the cells, and cells were incubated at 37 °C, 130 rpm, 8% CO_2_ and 70% humidity for 6–7 days. Supernatants were passed through a His-Trap Ni-NTA (17528601, GE Healthcare) and then separated on a Superdex 200 Increase size exclusion column (GE28-9909-44, GE Healthcare) in 20 mM Tris pH 9.0 (PHG0002, Sigma-Aldrich), 300 mM NaCl (S9888, Sigma-Aldrich) buffer to achieve size homogeneity. The heavy and light chains of S10A Fab and S10A IgG4 were co-expressed at 2(heavy):1(light) ratio using HEK293F cells as described elsewhere59. S10A Fab/IgG4 were purified using KappaSelect affinity (17545812, GE Healthcare). Fractions containing protein were pooled and run on a Superdex 200 Increase gel filtration column to obtain purified samples.

Siglec-10_d1-d2_–S10A Fab complex was obtained by transiently co-transfecting Siglec-10_d1-d2_ with the LC and HC of S10A Fab using HEK293S suspension cells at 2:2:1 ratio, respectively. Expression was achieved following the same procedure described for Siglec-10 alone. The protein complex was purified using HiTrap Ni-NTA column (GE Healthcare) followed by Superdex 200 Increase size exclusion column (GE Healthcare). In order to facilitate crystallization, the complex was treated with endoglycosidase H (EndoH), which cleaves high-mannose N-linked glycans present on N100 residue of Siglec-10_d1d2_. The complex was then incubated for 2hrs at RT with a 100x molar excess of V_H_H nanobody (REF) and then the ternary complex was purified using Superdex 200 Increase size exclusion column (GE Healthcare) in 20 mM Tris pH 9.0, 300 mM NaCl buffer to achieve size homogeneity.

Residues 1–215 of HMGB1 (UniProt accession Q5T7C4) were cloned into the pHLsec vector and codon-optimized for expression in human cells. The construct was transiently transfected into HEK293F suspension cells as described for Siglec-10. Culture supernatants were purified using a HisTrap Ni-NTA column (17528601, GE Healthcare) and further fractionated by size-exclusion chromatography on a Superdex 200 Increase column (GE28-9909-44, GE Healthcare) equilibrated in 20 mM Tris-HCl, pH 9.0 (PHG0002, Sigma-Aldrich), and 300 mM NaCl (S9888, Sigma-Aldrich) to achieve size homogeneity.

### Ligands

3’SLN and 6’SL were kindly provided by Prof Rob Field from the Manchester Institute of Biotechnology in the UK. The detailed synthesis of 6-*SO_3_*-6’-*SO_3_*-3’SLN can be found in *supplementary methods* and **Figures S15-19**.

### Crystallization, X-ray diffraction data collection, and 3D structure determination

Purified Siglec-10_d1d2_-S10A-V_H_H complex was concentrated to 13 mg/mL in a buffer containing 20 mM Tris pH 9.0 and 300 mM NaCl. Crystals were obtained by hanging drop vapor diffusion at 291 K in 0.16M ammonium sulfate, 0.08M sodium acetate and 20 % (v/v) PEG 4000 in 24-well plates after mixing 1 μL of the protein complex and 1 μL of the precipitant solution. Crystals were cryo-preserved by soaking them in mother liquor solution containing 25 % glycerol and flash frozen in liquid nitrogen. X-ray diffraction data was collected at the DLS IO4 synchrotron beamline at DIAMOND (UK). Data was processed using autoPROC in C2221 space group at 2.4 Å resolution. The structure was determined by molecular replacement using the Siglec-15 (PDB ID: 7ZOZ) and the light and heavy chains of 5G12 Fab (PDB ID: 7ZOR) as search model in Phaser. Crystals of Siglec-10–S10A-V_H_H-ɑ2-6SL complex were obtained by soaking of the already existing crystals of Siglec-10–S10A-V_H_H complex with 100 μM of the ligand (ɑ2-6 sialylactose). The ligand solution was added to the crystal containing drop at a 1:3 protein: ligand ratio.The soaked crystals were then collected and flash frozen in crystallization buffer containing 25% glycerol at different time-points (1 min, 5 mins, and 1 hour). X-ray diffraction data was collected at the DLS IO4 synchrotron beamline at DIAMOND (UK). Data was processed using autoPROC in C2221 space group at 3.27 Å resolution.

All structures were refined by manual building in Coot and using phenix.refine. PyMOL was utilized for structure analysis and figure rendering. All buried surface area values reported were calculated using EMBL-EBI PDBePISA. The crystal structures of Siglec-10_d1d2_/S10A and Siglec-10/S10A/ɑ2-6SL complex reported in this manuscript have been deposited in the Protein Data Bank, www.rcsb.org with PDB ID 29TG and 9R8E, respectively.

### Super-resolution fluorescence microscopy (DNA-PAINT) experiments

THP-1 cells overexpressing Siglec-10-GFP WT, 119A and 127A mutant cell lines were grown in RPMI 1640 + 10% FBS + 1% Penicillin/Streptomycin at a concentration of around 2 x 10^5^ cells/mL and passaged every 2-3 days according to their density. The cells were kept at 37 °C and 5% CO_2_ for cultivation. For the treatment 2.5 µg/mL of tunicamycin (Fisher Scientific) were added to the culture flasks and kept in the incubator for 48hr before seeding.

Cells were seeded at 50,000 cells cm⁻² into poly-L-lysine (PLL)–coated wells or chambers of microscope slides (µ-Slides with 8 wells with a glass bottom, 80801 and µ-Slide VI 0.5 Glass Bottom, 80607 from Ibidi) and incubated overnight at 37 °C and 5% CO₂ to allow for adherence. The next day the cells were fixed with 4% pre-warmed methanol-free PFA in 1xPBS for 15 min. After washing 3 times with 1xPBS, the cells were permeabilized with 0.125% TritonX-100 in 1xPBS for 5 min. After washing 3 times with 1xPBS, the cells were blocked with blocking buffer at 4 °C for 4 hr before adding 25 nM of the DNA-conjugated R1 (TCCTCCTCCTCCTCCTCCT) anti-GFP nanobody (clone 1H1, N0305 from Nanotag Biotechnologies) in blocking buffer (1x PBS, 1 mM EDTA, 0.02% Tween-20, 0.05% NaN_3_, 2% BSA, 0.05 mg mL^-1^ salmon sperm DNA) for overnight incubation at 4 °C. For the preparation of the anti-GFP nanobody, the nanobody was previously conjugated to a DBCO-PEG4-Maleimide linker (CLK-A108P, Jean Bioscience). After removing the unreacted linker with Amicon centrifugal filters (10,000 MWCO), the DBCO-nanobody was conjugated via DBCO-azide click chemistry to a DNA docking strand. DNA oligonucleotides modified with C3-azide and Cy3B were ordered from Metabion and MWG Eurofins. On the next day the sample was washed 3 times with 1x PBS, post-fixed with 4% PFA and 0.2% glutaraldehyde in 1x PBS for 10 min. Then, the cells were quenched with 0.2 M NH_4_Cl in 1xPBS for 5 min and washed 3 times with 1xPBS. 90 nm gold nanoparticles in 1:1 dilution in 1xPBS were incubated for 5 min at RT. After washing the cells 3 times with 1x PBS they were ready for imaging. The samples were imaged in the imaging buffer (1× PBS, 1 mM EDTA, 500 mM NaCl (pH 7.4), 0.02% Tween; supplemented with 1× Trolox, 1× PCA and 1× PCD) with an R1* (AGGAGGA-Cy3B) imager strand concentration of 400-500 pM for 40,000 frames with 100 ms exposure time per frame at 30 mW laser power after the objective, corresponding to a power density of 150 W cm^-2^.

Fluorescence imaging was carried out on an inverted microscope (Nikon Instruments, Eclipse Ti2) with the Perfect Focus System, applying an objective-type TIRF configuration equipped with an oil-immersion objective (Nikon Instruments, Apo SR TIRF×100, NA 1.49, Oil). A 560-nm laser (MPB Communications, 1 W) was used for excitation and coupled into the microscope via a Nikon manual TIRF module. The laser beam was passed through a cleanup filter (Chroma Technology, ZET561/10) and coupled into the microscope objective using a beam splitter (Chroma Technology, ZT561rdc). Fluorescence was spectrally filtered with an emission filter (Chroma Technology, ET600/50 m, and ET575lp) and imaged on an sCMOS camera (Hamamatsu Fusion BT) without further magnification, resulting in an effective pixel size of 130 nm (after 2×2 binning). TIR illumination was used for all measurements. The central 1152×1152 pixels (576×576 pixel after binning) of the camera were used as the region of interest. The camera’s scan mode was set to “ultra quiet scan” (readout noise = 0.7 e^-^ r.m.s., 80 μs readout time per line). Raw microscopy data were acquired using µManager (Version 2.0.1)66

Raw fluorescence data were reconstructed using the Picasso software package (the latest version is available at https://github.com/jungmannlab/picasso). For this, single molecules were localized in Picasso using the Gaussian least squares option. Drift correction was performed using AIM67 with gold particles (G-90-100 from Cytodiagnostics) serving as fiducials. For further analysis, single cells were picked and protein positions were determined with a clustering algorithm. For this, circular clusters of localizations centered around local maxima were identified and grouped. The centers of the localization groups were calculated as a weighted mean, using the squared inverse localization precisions as the weights. The oligomerization of Siglec-10 was analyzed using SPINNA analysis method66. Briefly, a model containing the coordinates of the proteins was created. The parameters used were: dimer distance = 16 nm, labelling uncertainty = 6 nm, labelling efficiency = 37 % according to previous reports67, granularity = 50. Then, simulations (n= 50 repeats per set of parameters) with different proportions of monomers and dimers were generated and the best fitting model was obtained.

### Biolayer interferometry

The binding affinities of S10A Fab to Siglec-10Fc was measured by BLI using the Octet R8 BLI system (Sartorius). AHC2 biosensors (18-5142, Sartorius) were hydrated in 1× kinetics buffer (PBS, pH 7.4, 0.002 % Tween, 0.01 % bovine serum albumin (BSA)) and loaded with 20 ng/µL of Siglec-10Fc for 200 s at 1000 rpm. Biosensors were then transferred into wells containing 1× kinetics buffer to baseline for 200 s before being transferred into wells containing a serial dilution of Fab starting at 120 nM and decreasing to 3.75 nM. The 200 s association phase was subsequently followed by a 200 s dissociation step in 1× kinetics. Analysis was performed using the Octet software (Sartorius), with a 1:1 fit model. All experiments were repeated in triplicate, values were averaged, and standard errors were calculated.

### NMR experiments

Ligand assignment and STD-NMR experiments with 6’SL were carried out on an 800 MHz Bruker spectrometer equipped with a triple-channel cryoprobe (Bruker, Billerica, MA, USA). All STD experiments were acquired at 298 K in all cases using an in-house pulse sequence.

STD-NMR experiments with 3’SLN and the sulfated 6-S-6’-S-3’SLN molecules were acquired on a Bruker Avance III 600 MHz spectrometer equipped with a 5-mm inverse detection triple-resonance cryogenic probe head with z-gradients and ligand assignment NMR experiments were acquired on a Bruker Avance 500 MHz equipped with a triple channel TXI probe. All spectra were acquired at 288 K. Detailed experimental procedures are provided in the *supporting materials and methods*.

### MD Simulations of the Siglec-10 and 6’SL complexes

MD simulations were carried out with AMBER 2268, implemented with ff14SB, which is an evolution of the Stony Brook modification of the Amber 99 force field force field (ff99SB)69 and GLYCAM0670 force fields. The structure of Siglec-10 was used for the calculations and the glycopeptides were build-up with Carbohydrate Builder (https://glycam.org/cb/). The complexes were generated using Protenix Server (https://protenix-server.com/login). Each complex was immersed in a 10 Å water box with TIP3P71 water molecules and charge neutralized by adding explicit counter ions (Na+). A two-stage geometry optimization approach was performed. The first stage minimizes only the positions of solvent molecules and ions, and the second stage is an unrestrained minimization of all the atoms in the simulation cell. The systems were then gently heated by incrementing the temperature from 0 to 300 K under a constant pressure of 1 atm and periodic boundary conditions. Harmonic restraints of 30 kcal·mol–1 were applied to the solute, and the Andersen temperature coupling scheme72. was used to control and equalize the temperature. The time step was kept at 1 fs during the heating stages, allowing potential inhomogeneities to self-adjust. Water molecules are treated with the SHAKE algorithm such that the angle between the hydrogen atoms is kept fixed. Long-range electrostatic effects are modelled using the particle-mesh-Ewald method73. An 8 Å cutoff was applied to Lennard-Jones and electrostatic interactions. Each system was equilibrated for 2 ns with a 2-fs time step at a constant volume and temperature of 300 K. Production trajectories were then run for additional 1.0 µs under the same simulation conditions.

### MD Simulations of the Siglec-10 in complex with GT1b

The 3D structure of full-length Siglec-10 ECD was generated with AlphaFold-2 (AF2)74 (AlphaFold Protein Structure Database number AF-Q96LC7)75. GT1b ganglioside from GlyTouCan (G08648UJ) was used. The equilibrium structure of the GT1b was sourced from the GlycoShape server76,77. The starting conformation of the complex was built to complement the 3D structure of GT1b, with an orientation selected to maximize the number of arginine residues engaging the trisialylated GT1b headgroup. Accordingly, the terminal α2-8-linked Neu5Ac was positioned to interact with the canonical R119, while R127, and R80 aligned to interact the internal and terminal α2-3-linked Neu5Ac residues, respectively. The complex was embedded in a symmetric 130 Å × 130 Å lipid bilayer composed of 60% 1,2-distearoyl-sn-glycero-3-phosphocholine (DSPC) and 40% cholesterol (CHL1) using the CHARMM-GUI Membrane Builder tool78. All molecular dynamics (MD) simulations were conducted with AMBER 22, using the CHARMM36m force field to parameterize the protein, lipids, and glycans. The system was first energy minimized for 5000 steps (2500 steps of steepest descent followed by 2500 steps of conjugate gradient minimization). Equilibration was performed following the CHARMM-GUI protocol79, consisting of six steps. Positional restraints of 10 kcal/mol·Å² were initially applied to the system and were gradually reduced during the NPT equilibration phase, progressively lowered to 5 kcal/mol·Å² for the protein and 2.5 kcal/mol·Å² for the membrane until they were completely removed. Temperature was maintained at 315.15 K using Langevin dynamics with a friction coefficient (gamma_ln) set to 1.0 ps − 1. and pressure was controlled semi-isotropically at 1 atm using the Berendsen barostat. Periodic boundary conditions were applied throughout. Long-range electrostatics were calculated using the Particle Mesh Ewald (PME) method with a cut-off of 11 Å. Bond lengths involving hydrogen atoms were constrained using the SHAKE algorithm, allowing for a 2 fs integration time step. Two independent 500-ns production runs were performed, resulting in a total sampling time of 1 μs. Trajectory analysis was conducted using Python3 scripts. Distances between key arginine side chains and the Neu5Ac residues of GT1b were calculated frame by frame using VMD. Occupancy was defined as the percentage of frames in which the atomic distance remained below 5 Å. All plots were generated using Matplotlib and Seaborn.

### Flow cytometry

Human T cells (1×10^6^) were collected on day 12 after activation with CD3/CD28 Dynabeads (11131D, Thermo Fisher). For evaluation of Siglec-10 binding to T cells, activated T cells were collected, washed with 1xPBS pH 7.4, and incubated with 8 μg/mL of Siglec-10Fc for 30 min at 4 °C. Cells where then washed with 1xPBS pH 7.4 at 450g for 5min, and incubated with anti-CD4 BUV395 (563550, BD Biosciences; 1:800) anti-CD8 APC/H7 (566855, BD Biosciences; 1:800), and anti-human IgG1Fc PE (12-4998-82, Thermo Fisher; 1:2500) for 30 min at 4 °C in the dark. In the case of CD24 detection, the anti-CD24 FITC (MHCD2401, ThermoFisher) was used at 1:250 for 30 mins on ice in the dark. After washing, cells were resuspended in 200 µL staining buffer containing DAPI (1:40,000) (D1306, Invitrogen) before acquisition in FACSymphony (BD Biosciences). Results were analyzed with FlowJo (BD Biosciences).

To investigate the sialic acid linkage dependency of Siglec10-Fc ligand binding on CD4+and CD8+T cells, spleens were harvested from C57BL/6J and sialyltransferase-deficient mice, ST6Gal1fl/flxCD4cre, ST3Gal1, and ST3Gal4, and processed into cell suspensions. Spleens were macerated in RPMI-1640 medium containing 2% FBS and filtered through a 40 μm mesh. Red blood cells were lysed using ammonium-chloride-potassium (ACK) buffer for 5 minutes at 4°C. Cells were washed and resuspended in HBSS containing either 2% FBS. Cells were stained on ice for 30 minutes using fluorophore-conjugated antibodies listed in the key material table. Siglec10-Fc, pre-complexed following a previously described protocol47, was incubated with the cell suspension to assess binding. After staining, cells were washed and resuspended in flow buffer supplemented with 1 µg/mL 7-AAD (cat# 420403, Biolegend) for viability assessment. Flow cytometric analysis was conducted on a BD Fortessa X-20 cytometer. CD4+T and CD8+T cell populations were gated B220-, CD3+, CD4+ and B220-, CD3+, CD8+ individually. The antibodies used for this experiment were rat anti-mouse B220 (clone:RA3-6B2) (cat# 563793 RRID: AB_2738427, BD Biosciences), rat anti-mouse CD3 (clone: 17A2) (cat# 100231; RRID: AB_11218805, BD Biolegend), rat anti-mouse CD4 (clone: GK1.5) (cat# 612952; RRID: AB_2813886, BD Biosciences), and mouse anti-mouse CD8a (clone: QA17A07) (cat# 155021; RRID: AB_2890707, BD Biolegend).

U937 cell lines were stained according to a previously described protocol80. Briefly, the cells (1 x 10^5^) were collected and washed with 1 x PBS pH 7.4. The cells were then stained with Siglec-10Fc complexed with a secondary anti-human IgG1 antibody (A-21445 ThermoFisher or 12-4998-82, Thermo Fisher). The cells were incubated for 45 mins at 4 °C in the dark, prior to being washed with 1 x PBS pH 7.4. The cells were then analyzed by either a Fortessa X-20 (BD Bioscience) or FACSymphony (BD Bioscience) flow cytometer. Data was analyzed using FlowJo software (BD Bioscience).

Siglec-10 blocking experiments were carried out either with S10A IgG4 produced in house, or anti-CD24 blocking SN3 antibody (MA5-11828, ThermoFisher). Either of the antibodies were used at 10 µg/mL and incubated with human T cells or breast cancer cell lines for 30mins, prior to washing the cells with 1x PBS pH7.4. Siglec-10Fc (8 µg/mL) was then precomplexed with anti-IgG1 Fc PE (12-4998-82, ThermoFisher; 1:2500) and incubated for 45 mins on ice in the dark. Cells were washed, stained with DAPI (1:20000) (D1306, Invitrogen) and analyzed on FACSymphony (BD Bioscience). Data was analysed using FlowJo software (BD Bioscience).

After 24h treatment of THP1 WT cells with tunicamycin (2.5 ug/mL), cells were stained using Streptavidin-APC (1:2500) which binds to ConcavalinA-biotin at a concentration of 5ug/mL.

### Siglec-10 binding to CD24 overexpressing cells

CD24 overexpressing cells (1 x 10^5^) were collected and washed with 1 x PBS pH 7.4. The corresponding wells were incubated with HMGB1 recombinant protein for 45mins on ice in the dark. Following the cells were washed with 1 x PBS pH 7.4. Next, the corresponding wells were treated with phospholipase C enzyme (ThermoFisher, P6466) according to manufactures instructions. Next the cells were stained with Siglec-10Fc for 1h on ice, prior to staining with anti-CD24 AF647 (BioLegend, 101817), anti-HMGB1 His (ACROBiosystems, HM1-H5220) and a secondary anti-human IgG1 PE (ThermoFisher, MA1-10377). Lastly, the cells were washed, stained with DAPI (1:20000) (D1306, Invitrogen) and analyzed on FACSymphony (BD Bioscience). Data was analysed using FlowJo software (BD Bioscience).

### Desialylation and deglycosylation of T cells

For sialic acid cleavage from T cells, human T cells (1 × 10^6^) were harvested 12 days post activation and incubated with 50µL of α2-3,6,8,9 Neuraminidase A (P0722, NEB) or α2-3 Neuraminidase S (P0743L, NEB) at 37 °C for 1 h. For global deglycosylation, activated T cells (1 × 10^6^) were treated with 50μL of Protein Deglycosylation Mix II (P6044, NEB) and incubated at 37 °C for 30 min according to manufacturer’s instructions. Deglycosylation efficiency was determined by lectin binding as described above.

### Antibody internalization

10µg/mL of S10A IgG4 was labelled with pHrodo deep red TFP Ester Dye (P35358, Invitrogen) according to manufacturer’s instructions. THP-1-CD19 cells (2 x 10^5^) were collected and washed with 1x PBS pH 7.4 for 5mins at 450g. Portion of the cells was incubated for 1h at 37°C with 10 μM cytochlalasin D (PHZ1063, ThermoFisher). Next, the cells were incubated with the labelled antibody at three different timepoints (0, 10, 30 mins and 30mins + Cyt D) at 4°C. The stained cells were then washed once with 1x PBS pH 7.4 for 5mins at 450g. After washing, cells were resuspended in 200 µL staining buffer containing DAPI (1:40,000) (D1306, Invitrogen) before acquisition in FACSymphony (BD Biosciences). Results were analyzed with FlowJo (BD Biosciences).

For confocal microscopy the sample preparation was the same as above. Cells were then cytospined onto slides using EZ Single Cytofunnel™, White (Epredia, Cat. #: A78710003) at 800 rpm for 5 minutes. The slides with adhered cells were fixed with 4% paraformaldehyde (PFA) for 15 minutes at room temperature and washed with 1x PBS pH 7.4. Subsequently, cells were stained with DAPI and mounted using ProLong Gold Antifade Reagent (Thermo Fisher Scientific, Cat. #: P3693). Imaging was performed using a Leica SP8 Lightning confocal microscope.

### CAR-T cell cytotoxicity

Co-culture of the CART cells (effector cells) with THP-1-CD19t-GFP-S10 various constructs (WT, R119A, R127A, R119/127A or ITIM mut) (target cells). The co-culture was set up in triplicate at effector:target (E:T) ratio set to 0.0625:1, with 5×10^4^ target cells/well in a final volume of 200 µL of non-supplemented AIM V media (12055091, ThermoFisher). The S10A IgG4 mAb was used in certain conditions at a constant concentration of 10µg/mL as a Siglec-10 blocking agent. After 16h co-incubation at 37°C, DAPI (62248, ThermoFisher) at the dilution of 1:2000 was added to the wells to allow for exclusion of dead cells. The total number of alive target cells was then measured by flow cytometry on the Attune NxT flow cytometer equipped with an autosampler (ThermoFisher). Percentage of cytotoxicity was calculated as follows: (no. alive GFP+ target cells only – no. alive GFP+ target cells with T cells)/(no. alive GFP+ target cells only) x 100.

The same experimental set up was used for checking the expression levels of Siglec-10 on CART cells at time 0 of the co-culture assay and at time 16h. In this case an anti-Siglec-10 mAb conjugated to APC (BioLegend, 347606) was used to measure Siglec-10 expression levels.

### Proximity labelling

Activated T cell samples from two different donors were analyzed. Siglec-10-Fc WT and Siglec-10Fc R119/127A were firstly preincubated with anti-hFc HRP (A01854 200, GenScript) for 30 min at 4 °C, then mixed with T cells and incubated at 4 °C for 1 h. After washing steps, the labelling solution (TBS + 10 mM H_2_O_2_ + 95 µM Biotin Tyramide (LS-3500, Iris Biotech)) was added to samples and incubated for 7 min with shaking, and the reaction was stopped by adding the quenching buffer (TBS + 100 µM ascorbic acid). Samples were then incubated with 30 μL Streptavidin magnetic beads (ThermoFisher, 88817) for 60 min. The non-bound material was removed by washing the beads on an Eppendorf magnet. The bead bound protein was then analysed by LC-MS to identify protein binders.

For the LC-MS analysis, an ‘on bead’ digestion protocol was followed using trypsinization (12.5µg/mL trypsin in 50mM ammonium bicarbonate) for 20min on ice. This was followed by re-hydration with 50mM ammonium bicarbonate, prior to bead removal. After digestion, peptides were dried out in a RVC2 25 speedvac concentrator (Christ) and resuspended in 0.1% formic acid (FA). Peptides were further desalted, resuspended in 0.1% FA using C18 stage tips (Millipore), and sonicated for 5min prior to analysis. Samples were analyzed in a timsTOF Pro with PASEF (Bruker Daltonics) coupled online to an Evosep ONE liquid chromatograph (Evosep). 200ng were directly loaded onto the Evosep ENDURANCE column (15cm vs 150µm, 1.9µm)andresolvedusingthe30samplesper-day standard protocol defined by the manufacturer (approximately 44min runs). timsTOF mass spectrometer was operated in Data-Dependent Acquisition mode (DDA) using the standard 1.1. second acquisition cycle method (HyStar Version 5.1.8.1). Protein identification and quantification was carried out using Byonic software (v2.16.11, Protein Metrics) through Proteome Discoverer v1.4 (Thermo Fisher). Searches were carried out against a database consisting of Homo sapiens (Uniprot/Swissprot, version 2020_04), with precursor and fragment tolerances of 20 ppm and 0.05Da respectively. Carbamidomethylation of Cysteine was considered as fixed modification whereas oxidation of Methionine was considered as variable modification. A decoy search was carried out to estimate the false discovery rate (FDR) of the searches. Only proteins with at least one peptide identified at FDR<1% were considered for further analysis. Spectral counts (SpC, the number of spectra that identifies peptides for a certain protein72) were used for the comparison of protein presence and abundance between conditions. Proteins with a SpC WT/Mut ratio>2, including those exclusively identified in the WT sample, were considered for further analysis and discussion.

### Western Blot

HEK293T CDD24 overexpressing cells and Ramos cells were used as a positive control, whereas HEK293F cell were used as a negative control. Total T cell lysates were collected at different time points (time 0, 24h, 48h and 72h post activation). All the cell types were lysed using RIPA buffer (ThermoFisher, 89900). Cell lysates were separated by 4-15% Mini-PROTEANTGX precast protein gel (BioRad, 4561083) and transferred to a 0.2 μm PVDF membrane (BioRad, 1704156) using a Trans-Blot Turbo transfer system (BioRad). The membrane was blocked for 1min 5% skim milk and 0.5% Tween-20 diluted in PBS. An overnight incubation with anti-CD24 mAb (1:500) (ThermoFisher, MA5-11828) was performed followed by five washes with PBS (containing 0.5% Tween-20) and incubation with secondary anti-mouse HRP conjugated antibodies (1:5000). After the incubation with the secondary antibody, five additional washes were carried out and Chemiluminescence detection was performed using Clarity Max Western ECL Substrate (BioRad, 170506) on an iBright CL1500 system (Invitrogen).

### Quantification and statistical analysis

Statistical analyses were performed using GraphPad Prism version 10.0. The test applied in each panel is specified in the figure legends.

## AUTHOR INFORMATION

### Author Contributions

K.S., J.J.B., and J.E.O. designed research; K.S., A.A.V., P.V., E.V.D., A.M., M.A.T., Z.J.C., E.N.S., L.H., S.D., I.O., M.E.L., M.J., F.M., S.Y.L., M.A., M.J.M., A.B., A.F., S.Y.L., J.E.D., and L.P.G. performed research; A.A.V., A.P., F.M., E.F., F.C., A.G., L.A.M., R.J., F.E., M.S.M., J.J.B., and J.E.O. contributed reagents/analytic tools; K.S., L.A.M., E.F., F.E., A.G., M.S.M., J.J.B., and J.E.O. analyzed data; and K.S. and J.E.O. wrote the paper.

### Notes

The authors declare no competing financial interests.

## Supporting Information

Detailed methods for all chemical syntheses, complete characterization data of the compounds, supporting figures and tables of calculated binding energies and interaction types, as well as additional references (PDF).

## ACKNOWLEDGMENTS

This work was supported by the European Research Council (788143-RECGLYCANMR to J.J.-B; ERC-2018-StG 804236-NEXTGEN-IO to A.P.) and the Marie-Skłodowska-Curie actions (ITN Glytunes grant agreement No 956758 to K.S.; and ITN DIRNANO grant agreement No 956544 to F.C.). X-ray diffraction experiments described in this paper were performed using beamlines DLSI04 at Diamond synchrotron (UK). The NMR spectrometers are part of the National NMR Network (PTNMR) and are partially supported by Infrastructure Project No 22161 (co-financed by FEDER through COMPETE 2020, POCI and PORL and FCT through PIDDAC). J.J.-B. and J.E.-O. acknowledge EC funding for the GLYCOTwinning project (No. 101079417). A.P.’s research is funded by “La Caixa” Foundation (HR21-00925), AECC (LABAE211744PALA), Fundación FERO, Ikerbasque, and BIOEF EITB MARATOIA BIO19/CP/002. We thank Agencia Estatal de Investigación of Spain for grants PID2019-107956RA-I00 (A.P.), PID2023-150777OB-I0 (J.E.-O.), CONSOLIDACION CNS2023-145146 (funded by MICIU/AEI /10.13039/501100011033 y por la Unión Europea NextGenerationEU/PRTR) (J.E-O), PID2024-161682OB-I00 (F.C.), ID2020-114178GB (R.B. & J.D.S.), RYC2018-024183-I (A.P.), PDC2022-133537-I00 (J.J.B. and J.E.-O.), PID2024-157610OB-I00 (J.J.-B.), PDC2025-166245-I00 (J.J.-B.), and the Severo Ochoa Center of Excellence Accreditation CEX2021-001136-S, all funded by MCIN/AEI/10.13039/501100011033 and by El FSE invierte en tu futuro, as well as CIBERES, and initiative of Instituto de Salud Carlos III (ISCIII, Spain) A.A.-V. receives funding from “La Caixa” Foundation (ID 100010434, LCF/BQ/DR20/11790022). A. B. (AECC Bizkaia Scientific Foundation, PRDVZ19003BOSC). F.C. and J.E-O acknowledges the Mizutani Foundation for Glycoscience (Grants 220115 and 250082, respectively); and J.E-O Caixa Junior Leader, BBVA Fundación for Leonardo Grant and Ramón Areces Fundation. A. G. thanks Agencia Estatal de Investigación of Spain for grants PID2023-150779OA-I00 and RYC2022-037985-I. A. T thanks Europe Marie Skłodowska-Curie Doctoral Networks (MSCA-DN) actions (Grant Agreement 101119499). We thank Prof Rob Field and Iakovia Ttoffi for the 3’SL and 6’SL ligands provided. For the ST3Gal1-/- and ST3Gal4-/- mice, we acknowledge Jamey Marth (Sanford Burnham Prebys Medical Discovery Institute, Infectious and Inflammatory Diseases, USA). IT Solutions at the University of Southampton is gratefully acknowledged for the generous allocation of computational resources on the HPC cluster Iridis. The Science Foundation of Ireland (SFI) Frontiers for the Future Programme is gratefully acknowledged for financial support of SDA postgraduate training (20/FFP-P/8809). The opinions, findings, and conclusions or recommendations expressed in this material are those of the author(s) and do not necessarily reflect the views of the Science Foundation of Ireland.

## DATA AVAILABILITY

The crystallographic data of Siglec-10–S10A Fab complex alone and in complex with 6’SL generated in this study have been deposited in the Protein Data Bank database under accession codes 9R7P and 9R8E. Any remaining information can be obtained from the corresponding author upon request. Source data are provided with this paper.

