## Supplemental Data 1 for "Molecular basis of Siglec-10 ligand recognition and antibody blockade"

**This PDF file includes:**

Supporting text – Supporting materials and methods
Tables S1 to S4
Figures S1 to S19
SI References

### Supporting Information Text

### Supporting Materials and Methods

***6-SO$_3$-6'-SO$_3$-3'SLN* synthesis (Figure S15).**

Reactions were carried out in oven-dried glassware. All reagents were purchased from commercial sources and were used without further purification unless noted. When necessary, the reaction solvents were dried by storing over 4 Å molecular sieves (48 h). Unless stated otherwise, all reactions were carried out at r.t. under a positive pressure of argon and were monitored by TLC on Silica Gel 60 $F_{254}$ (0.25 mm, E. Merck). Spots were detected under UV light or by charring with acidified *p*-anisaldehyde solution in EtOH. Unless otherwise indicated, all column chromatography was performed on Silica Gel (40–60 µM). $^1$H NMR spectra were recorded at 700 MHz or 600 MHz or 500 MHz and chemical shifts were referenced to either TMS (0.0, $CDCl_3$) or $CD_3OD$ (3.30, $CD_3OD$) or HOD (4.78, $D_2O$). $^1$H data were reported as though they were first order. $^{13}$C NMR (APT) spectra were recorded at 150 MHz or 125 MHz, and $^{13}$C chemical shifts were referenced to internal $CDCl_3$ (77.23, $CDCl_3$), or $CD_3OD$ (48.9, $CD_3OD$) or external acetone (31.07, $D_2O$). Organic solutions were concentrated under vacuum at <40 °C. Electrospray mass spectra were recorded on samples suspended in mixtures of THF with $CH_3OH$ and added NaCl.

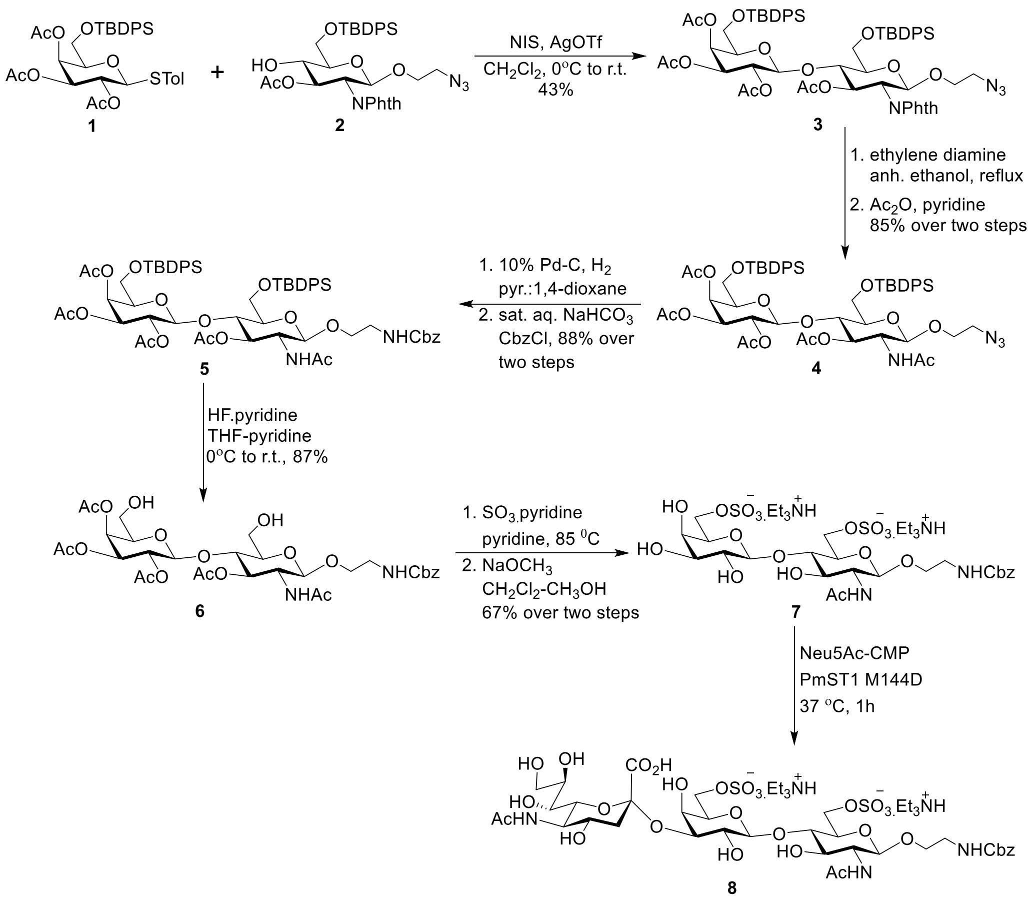


**2-azidoethyl 2-deoxy-2-*N*-phthlimido-4-*O*-[6-*O*-*tert*-butyldiphenylsilyl-2,3,4-tri-*O*-acetyl-*b*-D-galactopyranosyl]-6-*O*-*tert*-butyldiphenylsilyl-*b*-D-glucopyranoside (3)**

Thioglycoside **1** (1.3 g, 2.0 mmol) and alcohol **2** (1.1 g, 1.67 mmol) were dried under vacuum in the presence of $P_2O_5$ for 6 h prior to glycosylation. After drying, $CH_2Cl_2$ (45 mL) was added followed by powdered 4 Å molecular sieves (0.85 g) and stirred for 20 min. The reaction mixture was then cooled to 0°C and *N*-iodosuccinimide (580 mg, 2.6 mmol) and silver triflate (257 mg, 1.0 mmol) were added. The reaction was slowly allowed to warm to r.t. over 3h and was quenched by the addition of triethylamine until the pH of the solution was slightly basic. The reaction mixture was diluted with $CH_2Cl_2$ (40 mL) and filtered through Celite. The filtrate was washed with a saturated aq. solution of sodium thiosulphate (2 X 50 mL), water (30 mL) and brine (30 mL). The organic layer was separated, dried ($Na_2SO_4$), filtered and concentrated to a syrupy residue which was purified by column chromatography (1:1, hexanes/EtOAc) to yield the title compound **3** as a thick syrup. $R_f$ 0.16 (3:1, *n*-hexane–EtOAc, two runs); $^1H$ NMR (700 MHz, $CDCl_3$) δ 7.84 – 7.65

(m, 8H), 7.60 – 7.52 (m, 4H), 7.44 – 7.30 (m, 12H), 5.65 (dd, *J* = 10.7, 9.1 Hz, 1H), 5.53 (d, *J* = 3.3 Hz, 1H), 5.37 (d, *J* = 8.5 Hz, 1H), 5.04 – 4.93 (m, 2H), 4.79 (d, *J* = 7.7 Hz, 1H), 4.21 (dd, *J* = 10.8, 8.5 Hz, 1H), 4.14 – 4.07 (m, 1H), 3.98 – 3.90 (m, 3H), 3.73 – 3.65 (m, 2H), 3.57 (ddd, *J* = 11.0, 7.8, 3.5 Hz, 1H), 3.55 – 3.45 (m, 2H), 3.33 (ddd, *J* = 13.2, 7.7, 3.6 Hz, 1H), 3.17 (ddd, *J* = 13.2, 5.4, 3.5 Hz, 1H), 1.99 (s, 3H), 1.98 (s, 3H), 1.78 (s, 3H), 1.59 (d, *J* = 5.3 Hz, 3H), 1.09 (d, *J* = 2.9 Hz, 9H), 1.02 (s, 9H).; $^{13}$C NMR (126 MHz, $CDCl_3$) δ 170.08, 169.99, 168.86, 136.04, 135.61, 135.50, 133.36, 132.74, 132.69, 132.35, 130.07, 130.07, 129.99, 129.93, 129.93, 128.01, 127.89, 127.79, 127.71, 123.55, 123.55, 100.32, 97.83, 77.29, 77.04, 76.78, 75.34, 74.89, 72.89, 71.49, 70.80, 69.75, 68.18, 66.79, 61.28, 60.82, 54.92, 50.35, 26.90, 26.76, 20.68, 20.66, 20.54, 20.35, 19.41, 19.05.

**2-azidoethyl 2-deoxy-2-*N*-acetyl-4-*O*-[6-*O*-*tert*-butyldiphenylsilyl-2,3,4-tri-*O*-acetyl-*b*-D-galactopyranosyl]-6-*O*-*tert*-butyldiphenylsilyl-*b*-D-glucopyranoside (4) (Figure S16).**

To a solution of compound **3** (570 mg, 0.48 mmol) in anhydrous ethanol (27 mL) was added ethylene diamine (3 mL, 10% in the total volume) and the reaction mixture was heated under reflux for 22 h, then allowed to come to r.t. and concentrated under vacuum on a rotary evaporator. The thick syrupy residue was then dried for an hour under high vacuum before re-dissolving in pyridine (15 mL), cooled to < 4 ºC and acetic anhydride (5 mL) was added dropwise and left stirring at r.t. overnight. The reaction mixture was then diluted with dichloromethane (20 mL), cooled to < 4 °C and methanol (4 mL) was added. The reaction mixture was then concentrated to about 5 mL and was re-dissolved in dichloromethane (60 mL) and washed successively with water (1 × 40 mL), sat. aq. sodium bicarbonate (1 × 40 mL), and water (1 × 40 mL). The separated organic layer was dried ($Na_2SO_4$) and concentrated to a syrupy residue that was purified by column chromatography (3.5:6.5, *n*-hexane–EtOAc) to afford the title compound **4** (445 mg, 85% over two steps) as a fluffy solid; $R_f$ 0.12 (1:1, *n*-hexane–EtOAc); $^{1}$H NMR (500 MHz, $CDCl_3$) δ 7.78 – 7.72 (m, 4H), 7.63 – 7.58 (m, 4H), 7.51 – 7.36 (m, 12H), 5.62 – 5.54 (m, 2H), 5.09 – 4.96 (m, 3H), 4.71 (d, *J* = 7.2 Hz, 1H), 4.48 (d, *J* = 7.7 Hz, 1H), 4.09 (td, *J* = 9.1, 3.4 Hz, 2H), 4.01 – 3.87 (m, 3H), 3.75 (dd, *J* = 9.3, 5.4 Hz, 1H), 3.69 (dd, *J* = 8.3, 6.0 Hz, 1H), 3.63 (ddd, *J* = 11.0, 8.1, 3.3 Hz, 1H), 3.56 (t, *J* = 9.0 Hz, 1H), 3.51 – 3.44 (m, 1H), 3.41 – 3.35 (m, 1H), 3.29 (ddd, *J* = 13.2, 4.9, 3.3 Hz, 1H), 2.05 (d, *J* = 2.8 Hz, 3H),

2.02 (s, 3H), 1.96 (s, 3H), 1.81 (d, *J* = 1.6 Hz, 6H), 1.10 (s, 8H), 1.06 (s, 9H).; $^{13}$C NMR (126 MHz, $CDCl_3$) δ 170.88, 170.23, 170.05, 169.93, 169.14, 135.92, 135.62, 135.50, 135.46, 133.29, 132.66, 132.36, 130.09, 129.97, 128.01, 127.90, 127.81, 127.74, 100.93, 100.13, 77.30, 77.04, 76.79, 75.50, 73.59, 72.90, 72.34, 71.30, 69.69, 67.42, 66.73, 61.43, 60.72, 52.95, 50.66, 26.84, 26.76, 23.37, 20.68, 20.64, 20.60, 20.57, 19.33, 19.05.; HRMS (ESI) calcd. for $(M+Na)^+$ $C_{56}H_{72}NaN_4O_{15}$ 1096.4533, found 1096.4527.

**2-(Benzyloxycarbonylaminoethyl) 2-deoxy-2-*N*-acetyl-4-*O*-[6-*O*-*tert*-butyldiphenylsilyl-2,3,4-tri-*O*-acetyl-*b*-D-galactopyranosyl]-6-*O*-*tert*-butyldiphenylsilyl-*b*-D-glucopyranoside (5) (Figure S17).**

To a solution of compound **4** (444 mg, 0.4 mmol) in a solution of pyridine: 1,4-dioxane (12 mL: 10 mL) was added 10% Pd-C (160 mg) and stirred under an atmosphere of hydrogen (balloon) for 4 h. The catalyst was filtered off through a small pad of Celite and washed with a solution of pyridine: 1,4-dioxane (6 mL: 5 mL). The filtrate was concentrated to about 2 mL and was re-dissolved in 1,4-dioxane (27 mL) followed by the addition of sat. aq. sodium bicarbonate (4.0 mL). Under vigorous stirring CbzCl (0.3 mL) was added dropwise. Ensure that the pH of the solution does not become too acidic and remains slightly basic. When the reaction is complete as indicated by the tlc, it was concentrated on the rotary evaporator to about one third of its original volume and diluted with dichloromethane (100 mL). Water (75 mL) was added, stirred well and the organic layer was separated. The separated organic layer was once again washed with water (1 × 40 mL), separated, dried ($Na_2SO_4$), concentrated and the residue was purified by column chromatography (1:9, *n*-hexane–EtOAc) to afford the title compound **5** as a fluffy solid (429 mg) in 88% yield over two steps. $R_f$ 0.08 (2:3, *n*-hexane–EtOAc).; $^1$H NMR (500 MHz, $CDCl_3$) δ 7.77 – 7.70 (m, 4H), 7.60 (t, *J* = 6.2 Hz, 4H), 7.51 – 7.36 (m, 13H), 7.34 (s, 5H), 5.58 (d, *J* = 2.9 Hz, 1H), 5.23 (s, 1H), 5.17 – 4.97 (m, 4H), 4.90 (t, *J* = 9.3 Hz, 1H), 4.71 (d, *J* = 7.1 Hz, 1H), 4.31 (d, *J* = 7.8 Hz, 1H), 4.05 (dt, *J* = 17.3, 9.1 Hz, 2H), 3.96 – 3.81 (m, 3H), 3.73 (dd, *J* = 9.2, 5.4 Hz, 1H), 3.71 – 3.63 (m, 1H), 3.61 – 3.44 (m, 3H), 3.41 – 3.25 (m, 2H), 2.03 (d, *J* = 9.1 Hz, 6H), 1.90 (s, 3H), 1.80 (d, *J* = 3.9 Hz, 6H), 1.08 (s, 9H), 1.06 (s, 9H).; $^{13}$C NMR (126 MHz, $CDCl_3$) δ 171.11, 170.38, 170.03, 169.91, 169.13, 156.47, 136.57, 135.92, 135.62, 135.49, 135.45, 133.27, 132.66, 132.66, 132.24, 130.09, 130.01, 129.97, 128.50, 128.13, 128.01, 127.90, 127.81, 127.75, 101.62, 100.11, 77.30,

77.04, 76.79, 75.49, 73.52, 72.90, 72.58, 71.30, 69.69, 68.13, 66.75, 66.68, 61.38, 60.75, 53.35, 40.72, 26.83, 26.75, 23.31, 20.68, 20.64, 20.58, 19.33, 19.05.; HRMS (ESI) calcd. for $(M+Na)^+$ $C_{64}H_{80}NaN_2O_{17}Si_2$ 1204.4996, found 1204.4988.

**2-(Benzyloxycarbonylaminoethyl) 2-deoxy-2-*N*-acetyl-4-*O*-[2,3,4-tri-*O*-acetyl-*b*-D-galactopyranosyl]-*b*-D-glucopyranoside (6) (Figure S18).**

To a solution of the compound **5** (429 mg, 0.36 mmol) in THF–pyridine (9 mL: 5 mL) cooled to about < 4 ºC was added HF·pyridine (0.6 mL) dropwise. The reaction mixture was then allowed to come to r.t. and continued stirring for 24 h, before diluting with ethyl acetate (50 mL) and carefully neutralized by the addition of sat. aq. sodium bicarbonate solution. The organic layer was separated, once again washed with water (1 × 30 mL), separated, dried ($Na_2SO_4$), concentrated and the residue was purified by column chromatography (15:1, dichloromethane–methanol) to afford the title compound **6** as a fluffy solid (224 mg) in 87% yield. $R_f$ 0.13 (15:1, dichloromethane–methanol), $^1$H NMR (500 MHz, $CD_3OD$) δ 7.38 (s, 1H), 7.29 (d, *J* = 3.9 Hz, 4H), 7.25 (dd, *J* = 8.8, 4.5 Hz, 1H), 5.12 (dd, *J* = 10.3, 8.0 Hz, 1H), 5.03 (s, 2H), 4.98 – 4.90 (m, 1H), 4.53 (d, *J* = 8.0 Hz, 1H), 4.34 (d, *J* = 8.4 Hz, 1H), 4.25 – 4.14 (m, 2H), 3.94 (d, *J* = 2.8 Hz, 1H), 3.88 – 3.83 (m, 1H), 3.83 – 3.73 (m, 3H), 3.73 – 3.62 (m, 2H), 3.57 (dt, *J* = 10.1, 4.9 Hz, 1H), 3.30 (dt, *J* = 3.3, 1.6 Hz, 1H), 3.29 – 3.24 (m, 3H), 2.04 (s, 3H), 2.02 (s, 3H), 2.00 (s, 3H), 2.00 (s, 3H), 1.84 (s, 3H).; $^{13}$C NMR (126 MHz, $CD_3OD$) δ 173.01, 172.64, 172.18, 171.36, 137.85, 129.81, 129.45, 129.27, 102.62, 102.29, 78.93, 78.68, 78.42, 76.73, 76.65, 75.07, 74.58, 73.65, 71.08, 70.21, 68.03, 67.74, 63.96, 61.76, 61.16, 55.04, 50.65, 50.48, 50.30, 50.13, 49.96, 49.79, 49.62, 42.24, 42.12, 23.86, 22.03, 21.90, 21.85.; HRMS (ESI) calcd. for $(M+Na)^+$ $C_{32}H_{44}NaN_2O_{17}$ 728.2640, found 728.2636.

**2-(Benzyloxycarbonylaminoethyl) 2-deoxy-2-*N*-acetyl-4-*O*-[6-*O*-sulfo-*b*-D-galactopyranosyl]-6-*O*-Sulfo-*b*-D-glucopyranoside (7) (Figure S19).**

To a solution of compound **6** (121 mg, 0.17 mmol) in dry pyridine (16 mL) at < 5 ºC under nitrogen was added $SO_3$·pyridine complex in portions (1.05 g, 6.6 mmol). The reaction mixture was then lowered to an oil bath kept at 85 ºC and stirred for 40 min. The reaction mixture was then allowed to cool to r.t. and triethylamine (2.5 mL) was added dropwise, concentrated to a syrupy residue which was partially purified (some salts will be present after this purification) by column chromatography on silica gel (dichloromethane–methanol 15:1 to 7:1, to 5:1) to yield the sulfated compound as a semi solid. $R_f$ 0.04 (7:1, dichloromethane–methanol); This semi solid was then dried well under vacuum overnight and dissolved in $CH_2Cl_2$–$CH_3OH$ (1:1, 12 mL) followed by the addition of sodium methoxide in $CH_3OH$ to bring the pH of the reaction mixture strictly to 8–9. The solution was stirred for 48h and was then neutralized by the addition of pre-washed Amberlite IR 120 $H^+$ resin, filtered and the filtrate was concentrated to a syrupy residue that was purified first on a silica gel column (chloroform–methanol 7:1 to 3:1 and then $^i$PrOH:$NH_4OH$:$H_2O$7:1:1 to 7:2:1) and then by a C-18 column (water-methanol, gradient elution) to afford the title compound **7** (80 mg, 67% over two steps) as a white fluffy material. $R_f$ 0.42 (7:2:1 $^i$PrOH:$NH_4OH$:$H_2O$); $^1$H NMR (600 MHz, $D_2O$) δ 7.65 – 7.20 (m, 5H), 5.25 – 4.88 (m, 2H), 4.51 (d, *J* = 7.9 Hz, 2H), 4.37 (d, *J* = 10.8 Hz, 1H), 4.25 (dd, *J* = 11.1, 5.0 Hz, 1H), 4.20 – 4.12 (m, 2H), 3.98 – 3.92 (m, 2H), 3.86 (dt, *J* = 10.2, 4.9 Hz, 1H), 3.77 (s, 1H), 3.72 – 3.63 (m, 5H), 3.51 (dd, *J* = 10.0, 7.9 Hz, 1H), 3.34 – 3.21 (m, 2H), 3.16 (q, *J* = 7.3 Hz, 3H), 1.93 (s, 3H), 1.24 (t, *J* = 7.3 Hz, 4H).; $^{13}$C NMR (126 MHz, $D_2O$) δ 175.60, 159.32, 137.48, 129.81, 129.41, 128.81, 103.75, 102.21, 79.66, 73.75, 73.56, 73.30, 73.10, 71.84, 69.78, 69.25, 68.05, 67.54, 56.04, 47.69, 41.35, 23.13, 9.24.; HRMS (ESI) calcd. for $(M-H)^-$ $C_{24}H_{35}NaN_2O_{19}$ 720. 1354, found 720.1361.

**2-(Benzyloxycarbonyl) 5-acetamido-3,5-dideoxy-D-glycero-α-D-galacto-2-nonulopyranosylonate-(2→3)-(6-*O*-sulfate)-β-D-galactopyranosyl-(1→4)-2-N-acetyl-2-deoxy-(6-*O*-sulfate)-β-D-glucopyranoside (8).**

Compound **7** (28 mg, 1 eq) and CMP-Neu5Ac (70 mg, 2.5 eq) were dissolved in 100 mM Tris-HCl buffer containing 20 mM $MgSO_4$ (2.6 mL, pH 8.5). PmST1 M144D (0.4 mg/ml) was added and the reaction was placed in a shaking incubator (37 °C, 1 h) and was monitored by TLC in iPrOH:$NH_4OH$:$H_2O$ (5:2:1 by volume). The reaction was stopped by dilution with a 4-fold of 100% ethanol and put at −20 °C for 1 h to precipitate the enzyme. Precipitated protein was centrifuged (3700 × g, 30 min, 4 °C), the supernatant was decanted into a round bottom flask and evaporated. The residue was resuspended in water and purified on a LH20 column equilibrated in $H_2O$: MeOH (1:1) giving compound **8** in 60% yield (23 mg). $^1$H NMR (500 MHz, $D_2O$) δ = 7.42 (m, 5H), 5.12 (m, 2H), 4.59 (d, J = 7.5 Hz, 1H), 4.52 (d, J = 8.5, 1H), 4.39 (d, J = 11 Hz, 1 H), 4.28 (d, J = 9.5 Hz, 1H)

4.12 (dd, J = 9.1, 9.5 Hz,1H), 3.82-3.98 (m, 6H), 3.59-3.80 (m, 13H), 3.51-3.58 (t, J = 9.5 Hz, 1H), 3.24–3.28 (m, 2H), 2.72 (dd, J = 5, 12.5 Hz, 1H), 2.03 (s, 3H), 1.93 (s, 3H), 1.80 (t, J = 12.5 Hz, 1H).

### NMR experiments

#### *STD-NMR experiments*

Samples for STD-NMR experiments were prepared in 100% $D_2O$ with phosphate-buffered saline 10 mM, 300 mM NaCl, 0.05% NaN3, pH 7.4. Siglec-10 concentrations ranged between 10 and 12 μM for all experiments, and the ligands of interest were respectively added in a 1:100 protein-to-ligand ratio. STD experiments were recorded employing a train of selective 50 ms PC9 pulses focused on the aliphatic region for the on-resonance spectra, at 0.85 ppm. The reference *off-resonance* spectra were acquired irradiating at  100 ppm. Given the large size of the analyzed constructs, a short T2 filter (30 ms) was enough to suppress any residual signal from the protein, and no blank experiment was needed. Ligand blanks were performed to rule out significant direct irradiation of the ligand under the measuring conditions.

For each Siglec-10 /6´SL ligand pair, at least four independent STD experiments were acquired at four different saturation times: 1, 2, 4 and 6 seconds. From these four experiments, the absolute STD percentages (STDabs) were calculated for each ligand proton by comparing the peak intensities of the on ($I_{on}$) and off ($I_{off}$) spectra **(Table S3)**. Considering the 1:100 protein/ligand ratio used, the STD effects are here calculated and reported as amplification factors ($STD_{AF}$) using the following expression:

$$\mathrm{STD_{AF}} = \mathrm{STD_{abs}} \cdot \frac{\mathrm{L_0}}{\mathrm{P_0}} = \frac{\mathrm{I_{off}} - \mathrm{I_{on}}}{\mathrm{I_{off}}} \cdot 100 \qquad (1)$$

Where $L_0$ and $P_0$ are the total amounts of ligand and protein used (mol/L), respectively. The obtained $STD_{AF}$ values for the different saturation times (tsat) were then fitted to derive the STD build-up curve expression for each individual proton (Eq. 2), where $k_{sat}$ stands for the saturation rate in $s^{-1}$ and $STD_{AF}^{MAX}$ is the maximum expected amount of transferred saturation for a given proton in the set experimental conditions (**Table S3**):

$$\mathbf{STD_{AF}} = \mathrm{STD_{AF}^{MAX}} \cdot \left(1 - \mathrm{e}^{-\mathrm{k_{sat}}\mathbf{t_{sat}}}\right) \qquad (2)$$

To analyze the binding epitope in each case, the Initial Slopes Method[47] was used to erase any possible bias arising from the different relaxation rates of the ligand protons. The STD at zero time ("$STD_0$", no $T_1/T_2$ contribution) for a given proton is the slope of the tangent line to its STD build-up curve at tsat = 0, and its value is provided by Eq. 4:

$$\frac{d(STD_{AF})}{dt} = k_{sat} \cdot STD_{AF}^{MAX} \cdot e^{-k_{sat}\mathbf{t_{sat}}} \qquad (3)$$

$$\text{At } t_{sat} = 0 \quad \rightarrow \quad \mathbf{STD_0} = k_{sat} \cdot STD_{AF}^{MAX} \qquad (4)$$

To build the binding epitope maps of the ligands in relative scale, the highest $STD_0$ value found in each case was arbitrarily assigned a value of $\%STD_{0,rel} = 100\%$, while the rest of relative values for the other protons were derived accordingly (**Table S3**):

$$\%STD_{0,rel} = \frac{STD_0}{\text{highest } STD_0} \cdot 100 \qquad (5)$$

**Tables**

**Table S1. Crystallographic data collection and refinement statistics.**

| | **Siglec-10/S10A Fab** | **Siglec-10/S10A Fab/ 6´SLA** |
|---|---|---|
| **PDB ID** | **29TG** | **9R8E** |
| **Data collection Statistics** | | |
| **Wavelength (Å)** | 0.99991 | 0.67017 |
| **Resolution (Å)** | 63.54- 2.69 (2.74- 2.69) | 73.87- 3.67 (3.83- 3.67) |
| **Space group** | C 2 2 21 | C 2 2 21 |
| **Unit cell dimensions** | | |
| **a, b, c (Å)** | 127.11, 148.21 228.08 | 127.36, 146.37, 230.64 |
| **α, β, γ (°)** | 90, 90, 90 | 90, 90, 90 |
| **Multiplicity** | 13.5 (12.9) | 9.1 (7.4) |
| **Completeness (%)** | 95.96 (95.5) | 99.9 (78.1) |
| **Mean I/σI** | 24.5 (1.4) | 19.4 (1.5) |
| **Wilson B-factor (Å2)** | 35.23 | 100.45 |
| **Rmerge** | 0.076 (2.275) | 0.086 (1.220) |
| **Rpim** | 0.022 (0.669) | 0.030 (0.487) |
| **CC1/2** | 99.7 (61.5) | 99.3 (58) |
| **Refinement Statistics** | | |
| **Resolution (Å)** | 63.54- 2.69 | 73.87- 3.67 |
| **No. reflections** | 57,412 (2,804) | 23,512 (2,704) |
| **Rwork/Rfree** | 0.2403 /0.2862 | 0.2208/0.2680 |
| **No. atoms:** | 11,258 | 11,582 |
| **Protein** | 11,243 | 11,523 |
| **Ligand** | 15 | 59 |
| **RMS (bonds)** | 0.016 | 0.003 |
| **RMS (angles)** | 0.65 | 0.63 |
| **Ramachandran statistics:** | | |
| **favoured (%)** | 93.44 | 90.38 |

| | | |
|---|---|---|
| **allowed (%)** | 5.92 | 8.20 |
| **outliers (%)** | 0.63 | 1.42 |
| **Rotamer outliers (%)** | 1.48 | 0.08 |
| **Average B (Å2):** | 57.31 | 107.94 |
| **Protein** | 57.29 | 107.79 |
| **Ligand** | 72.65 | 138.76 |

**Table S2. Residues involved in the Siglec-10-S10A Fab interactions in the crystal.**

| Contact type | Siglec-10 residue | S10A HC residue |
|---|---|---|
| Hydrogen bonds | S37 | T58 |
| | T41 | Y50 |
| | T41 | E59 |
| | Q62 | S31 |
| | S63 | Y106 |
| | V66 | Y103 |
| | R106 | E59 |
| | S37 | T58 |
| | Q62 | Y103 |
| | Q62 | W33 |
| | R64 | A104 |
| | R64 | G105 |
| Salt bridges | R1 | E59 |

| Contact type | Siglec-10 residue | S10A LC residue |
|---|---|---|
| Hydrogen bonds | E65 | Y32 |
| | R64 | D91 |
| | R111 | E92 |
| | E65 | R50 |
| | E65 | R53 |
| Salt bridges | R64 | D91 |
| | R111 | E92 |
| | E65 | R50 |
| | E65 | R53 |

**Table S3. Summary of the $STD_{AF}$ values measured for each ligand proton at the different saturation times established,** using Eq. 1. All values were tracked using non-overlapping regions of the corresponding singlets/multiplets. Zero values mean non-observable STD or too weak to be tracked. For those cases where the overlapping could not be avoided at all, the STDs are jointly reported as unique values for the overlapping protons (as sum). The $STD_0$ column (grey) summarizes the realistic STD factors with no $T_1/T_2$ contribution, calculated from Eq. 4 using the fitting parameters previously estimated via Eq. 2. The relative $STD_0$ values from Eq. 5 are shown in the last column (green), pre-assigning 100% to the highest one in each case. Ligand blanks showed negligible contributions of ca.

| **Siglec-10Fc WT + 100eq 6´SL** | | | | | | | | | |
|---|---|---|---|---|---|---|---|---|---|
| **PROTON** | **$STD_{AF}$ at different $t_{sat}$ (s)** | | | | **Fitting (Initial Slopes)** | | | **$STD_0$** | **$\%STD_{0,REL}$** |
| | **6 s** | **4 s** | **2 s** | **1 s** | **$STD_{AF}^{MAX}$** | **$k_{sat}$ ($s^{-1}$)** | **$R^2$** | | |
| $H3_{ax}Sia$ | 29 | 23 | 17 | 11 | 30.99 | 0.3926 | 0.9767 | 12 | 22 |
| $H3_{eq}Sia$ | 32 | 24 | 18 | 11 | 35.65 | 0.3323 | 0.9678 | 12 | 22 |
| H4Sia + H3Gal | 74 | 67 | 33 | 24 | 98.23 | 0.2483 | 0.9637 | 24 | 45 |
| H5Sia | 66 | 57 | 35 | 16 | 86.35 | 0.2518 | 0.9884 | 22 | 40 |
| Methyl (C5 Sia) | 183 | 163 | 84 | 43 | 268.80 | 0.2030 | 0.9773 | 55 | 100 |
| H6Sia | 101 | 97 | 44 | 18 | 164.60 | 0.1761 | 0.9340 | 29 | 53 |
| H7Sia | 116 | 109 | 62 | 21 | 161.30 | 0.2348 | 0.9466 | 38 | 69 |
| H8Sia + $H9_aSia$ | 49 | 48 | 26 | 12 | 64.16 | 0.2741 | 0.9504 | 18 | 32 |
| $H9_bSia$ | 46 | 47 | 25 | 11 | 60.10 | 0.2856 | 0.9293 | 17 | 31 |
| H1Gal | 21 | 18 | 14 | 8 | 21.98 | 0.4720 | 0.9902 | 10 | 19 |
| H2Gal | 35 | 35 | 25 | 13 | 38.60 | 0.4752 | 0.9859 | 18 | 34 |
| H4Gal | 26 | 30 | 12 | 7 | 41.20 | 0.2152 | 0.9210 | 9 | 16 |
| H5Gal | 26 | 20 | 14 | 7 | 32.70 | 0.2557 | 0.9901 | 8 | 15 |
| H6Gal | 24 | 21 | 16 | 7 | 26.70 | 0.3934 | 0.9698 | 11 | 19 |
| H1Glc | 25 | 17 | 10 | 0 | 72.70 | 0.0694 | 0.9931 | 5 | 9 |
| H2Glc | 26 | 14 | 0 | 0 | N/A | N/A | N/A | N/A | N/A |
| H3/H4/H5Glc | 22 | 18 | 12 | 6 | 27.20 | 0.2741 | 0.9962 | 7 | 14 |
| H6Glc | 22 | 18 | 0 | 0 | N/A | N/A | N/A | N/A | N/A |

| **Siglec-10Fc R127A + 100eq 6´SL** | | | | | | | | | |
|---|---|---|---|---|---|---|---|---|---|
| **PROTON** | **$STD_{AF}$ at different $t_{sat}$ (s)** | | | | **Fitting (Initial Slopes)** | | | **$STD_0$** | **$\%STD_{0,REL}$** |

| | **6 s** | **4 s** | **2 s** | **1 s** | **$STD_{AF}^{M}$ AX** | **$k_{sat}$ ($s^{-1}$)** | **$R^2$** | | |
|---|---|---|---|---|---|---|---|---|---|
| $H3_{ax}$Sia | 0 | 0 | 0 | 0 | N/A | N/A | N/A | N/A | N/A |
| $H3_{eq}$Sia | 23 | 13 | 0 | 0 | N/A | N/A | N/A | N/A | N/A |
| H4Sia + H3Gal | 41 | 35 | 14 | 8 | 97.88 | 0.0951 | 0.9632 | 9 | 24 |
| H5Sia | 33 | 32 | 17 | 8 | 43.99 | 0.2609 | 0.9531 | 11 | 30 |
| Methyl (C5 Sia) | 108 | 94 | 58 | 32 | 132.30 | 0.2927 | 0.9965 | 39 | 100 |
| H6Sia | 53 | 46 | 28 | 12 | 70.86 | 0.2416 | 0.9834 | 17 | 44 |
| H7Sia | 83 | 71 | 39 | 24 | 111.80 | 0.2338 | 0.9914 | 26 | 67 |
| H8Sia + $H9_a$Sia | 29 | 31 | 16 | 8 | 36.41 | 0.3090 | 0.9489 | 11 | 29 |
| $H9_b$Sia | 29 | 27 | 17 | 12 | 31.80 | 0.4285 | 0.9820 | 14 | 35 |
| H1Gal | 14 | 12 | 0 | 0 | N/A | N/A | N/A | N/A | N/A |
| H2Gal | 29 | 23 | 14 | 10 | 35.78 | 0.2678 | 0.9835 | 10 | 25 |
| H4Gal | 9 | 11 | 9 | 5 | 10.38 | 0.7910 | 0.9347 | 8 | 21 |
| H5Gal | 18 | 16 | 15 | 8 | 18.00 | 0.6997 | 0.9279 | 13 | 33 |
| H6Gal | 18 | 14 | 12 | 7 | 18.12 | 0.4853 | 0.9424 | 9 | 23 |
| H1Glc | 0 | 0 | 0 | 0 | N/A | N/A | N/A | N/A | N/A |
| H2Glc | 0 | 0 | 0 | 0 | N/A | N/A | N/A | N/A | N/A |
| H3/H4/H5Glc | 15 | 12 | 7 | 0 | 21.60 | 0.1999 | 0.9989 | 4 | 11 |
| H6Glc | 38 | 34 | 15 | 0 | 61.50 | 0.1708 | 0.9254 | 11 | 27 |

*** Fitting is linear rather than exponential, suggesting unspecific saturation effects..*

**Table S4. List of counter binders for Siglec-10 expressed in human T cells identified by proximity labelling. n=2 biologically independent T cell donors.** The table shows uniprot protein IDs, gene names, protein names and the total number of distinct peptide sequences identified. Siglec-10 R119/127A mutant and anti-Fc IgG HRP were used as controls. The peptides identified in the controls were subtracted from the final peptide count.

| Protein ID | Gene name | Protein name | Total peptide count |
|---|---|---|---|
| P11498 | PC | Pyruvate carboxylase, mitochondrial | 48 |
| Q63HN8 | RNF213 | E3 ubiquitin-protein ligase RNF213 | 48 |
| Q09666 | AHNAK | Neuroblast differentiation-associated protein AHNAK | 46 |
| P05165 | PCCA | Propionyl-CoA carboxylase alpha chain, mitochondrial | 42 |
| Q5D862 | FLG2 | Filaggrin-2 | 42 |
| P38646 | HSPA9 | Stress-70 protein, mitochondrial | 41 |
| P52272 | HNRNPM | Heterogeneous nuclear ribonucleoprotein M | 36 |
| Q96RQ3 | MCCC1 | Methylcrotonoyl-CoA carboxylase subunit alpha, mitochondrial | 34 |
| P14618 | PKM | Pyruvate kinase PKM | 33 |
| B0I1T2 | MYO1G | Unconventional myosin-Ig | 33 |
| P08238 | HSP90AB1 | Heat shock protein HSP 90-beta | 32 |
| Q7L014 | DDX46 | Probable ATP-dependent RNA helicase DDX46 | 32 |
| P11142 | HSPA8 | Heat shock cognate 71 kDa protein | 30 |
| P42704 | LRPPRC | Leucine-rich PPR motif-containing protein, mitochondrial | 30 |
| P10809 | HSPD1 | 60 kDa heat shock protein, mitochondrial | 29 |
| P08575 | PTPRC | Receptor-type tyrosine-protein phosphatase C | 29 |
| P14923 | JUP | Junction plakoglobin | 29 |

| | | | |
|---|---|---|---|
| P11387 | TOP1 | DNA topoisomerase 1 | 29 |
| Q6ZU80 | CEP128 | Centrosomal protein of 128 kDa | 29 |
| P06733 | ENO1 | Alpha-enolase | 27 |
| Q15233 | NONO | Non-POU domain-containing octamer-binding protein | 26 |
| Q02413 | DSG1 | Desmoglein-1 | 26 |
| P60709 | ACTB | Actin, cytoplasmic 1 | 25 |
| P20930 | FLG | Filaggrin | 25 |
| P36578 | RPL4 | Large ribosomal subunit protein uL4 | 24 |
| P49748 | ACADVL | Very long-chain specific acyl-CoA dehydrogenase, mitochondrial | 24 |
| Q9NWB6 | ARGLU1 | Arginine and glutamate-rich protein 1 | 23 |
| P07900 | HSP90AA1 | Heat shock protein HSP 90-alpha | 23 |
| P05166 | PCCB | Propionyl-CoA carboxylase beta chain, mitochondrial | 22 |
| P04406 | GAPDH | Glyceraldehyde-3-phosphate dehydrogenase | 22 |
| P68363 | TUBA1B | Tubulin alpha-1B chain | 21 |
| Q02878 | RPL6 | Large ribosomal subunit protein eL6 | 21 |
| P02545 | LMNA | Prelamin-A/C | 21 |
| P49411 | TUFM | Elongation factor Tu, mitochondrial | 21 |
| P22626 | HNRNPA2B1 | Heterogeneous nuclear ribonucleoproteins A2/B1 | 21 |
| Q08188 | TGM3 | Protein-glutamine gamma-glutamyltransferase E | 21 |
| P48735 | IDH2 | Isocitrate dehydrogenase [NADP], mitochondrial | 21 |
| P68104 | EEF1A1 | Elongation factor 1-alpha 1 | 20 |
| P07437 | TUBB | Tubulin beta chain | 20 |

| | | | |
|---|---|---|---|
| P38159 | RBMX | RNA-binding motif protein, X chromosome | 20 |
| Q00839 | HNRNPU | Heterogeneous nuclear ribonucleoprotein U | 20 |
| P11586 | MTHFD1 | C-1-tetrahydrofolate synthase, cytoplasmic | 20 |
| P68032 | ACTC1 | Actin, alpha cardiac muscle 1 | 20 |
| Q9Y383 | LUC7L2 | Putative RNA-binding protein Luc7-like 2 | 19 |
| Q96PK6 | RBM14 | RNA-binding protein 14 | 19 |
| P07355 | ANXA2 | Annexin A2 | 19 |
| P68371 | TUBB4B | Tubulin beta-4B chain | 19 |
| Q16666 | IFI16 | Gamma-interferon-inducible protein 16 | 19 |
| P60842 | EIF4A1 | Eukaryotic initiation factor 4A-I | 19 |
| P68366 | TUBA4A | Tubulin alpha-4A chain | 19 |
| P40939 | HADHA | Trifunctional enzyme subunit alpha, mitochondrial | 19 |
| P0DMV8 | HSPA1A | Heat shock 70 kDa protein 1A | 19 |
| Q01546 | KRT76 | Keratin, type II cytoskeletal 2 oral | 19 |
| P16403 | H1-2 | Histone H1.2 | 18 |
| P20701 | ITGAL | Integrin alpha-L | 18 |
| P62701 | RPS4X | Small ribosomal subunit protein eS4, X isoform | 18 |
| P39023 | RPL3 | Large ribosomal subunit protein uL3 | 18 |
| P42224 | STAT1 | Signal transducer and activator of transcription 1-alpha/beta | 18 |
| O00763 | ACACB | Acetyl-CoA carboxylase 2 | 18 |
| Q01813 | PFKP | ATP-dependent 6-phosphofructokinase, platelet type | 18 |
| P23246 | SFPQ | Splicing factor, proline- and glutamine-rich | 18 |
| P04439 | HLA-A | HLA class I histocompatibility antigen, A alpha chain | 17 |

| | | | |
|---|---|---|---|
| P2339<br>6 | RPS3 | Small ribosomal subunit protein uS3 | 17 |
| P1041<br>2 | H1-4 | Histone H1.4 | 17 |
| P1640<br>2 | H1-3 | Histone H1.3 | 17 |
| P0657<br>6 | ATP5<br>F1B | ATP synthase subunit beta, mitochondrial | 17 |
| Q1383<br>5 | PKP1 | Plakophilin-1 | 17 |
| P0813<br>3 | ANX<br>A6 | Annexin A6 | 17 |
| Q0821<br>1 | DHX9 | ATP-dependent RNA helicase A | 17 |
| Q9Y3<br>Z3 | SAM<br>HD1 | Deoxynucleoside triphosphate triphosphohydrolase SAMHD1 | 17 |
| P3489<br>7 | SHMT<br>2 | Serine hydroxymethyltransferase, mitochondrial | 17 |
| Q9979<br>8 | ACO2 | Aconitate hydratase, mitochondrial | 17 |
| P0055<br>8 | PGK1 | Phosphoglycerate kinase 1 | 17 |
| P2273<br>5 | TGM1 | Protein-glutamine gamma-glutamyltransferase K | 17 |
| Q0221<br>8 | OGD<br>H | 2-oxoglutarate dehydrogenase complex component E1 | 17 |
| P7852<br>7 | PRKD<br>C | DNA-dependent protein kinase catalytic subunit | 17 |
| P1901<br>2 | KRT1<br>5 | Keratin, type I cytoskeletal 15 | 17 |
| P3677<br>6 | LONP<br>1 | Lon protease homolog, mitochondrial | 17 |
| P0276<br>8 | ALB | Albumin | 16 |
| O9523<br>2 | LUC7<br>L3 | Luc7-like protein 3 | 16 |
| Q14C<br>N4 | KRT7<br>2 | Keratin, type II cytoskeletal 72 | 16 |
| P6124<br>7 | RPS3<br>A | Small ribosomal subunit protein eS1 | 16 |
| P6242<br>4 | RPL7<br>A | Large ribosomal subunit protein eL8 | 16 |
| P0965<br>1 | HNR<br>NPA1 | Heterogeneous nuclear ribonucleoprotein A1 | 16 |
| P0404<br>0 | CAT | Catalase | 16 |

| | | | |
|---|---|---|---|
| P1102<br>1 | HSPA<br>5 | Endoplasmic reticulum chaperone BiP | 16 |
| P1379<br>6 | LCP1 | Plastin-2 | 16 |
| Q9UQ<br>35 | SRRM<br>2 | Serine/arginine repetitive matrix protein 2 | 16 |
| Q9Y6<br>N5 | SQOR | Sulfide:quinone oxidoreductase, mitochondrial | 16 |
| O0057<br>1 | DDX3<br>X | ATP-dependent RNA helicase DDX3X | 16 |
| P4932<br>7 | FASN | Fatty acid synthase | 16 |
| Q9Y4<br>90 | TLN1 | Talin-1 | 16 |
| Q9C0<br>75 | KRT2<br>3 | Keratin, type I cytoskeletal 23 | 16 |
| O9498<br>6 | CEP1<br>52 | Centrosomal protein of 152 kDa | 16 |
| Q8NA<br>72 | POC5 | Centrosomal protein POC5 | 16 |
| P0073<br>8 | HP | Haptoglobin | 15 |
| P1032<br>1 | HLA-<br>C | HLA class I histocompatibility antigen, C alpha chain | 15 |
| Q0795<br>5 | SRSF<br>1 | Serine/arginine-rich splicing factor 1 | 15 |
| O1501<br>4 | ZNF6<br>09 | Zinc finger protein 609 | 15 |
| Q9284<br>1 | DDX1<br>7 | Probable ATP-dependent RNA helicase DDX17 | 15 |
| P0408<br>3 | ANX<br>A1 | Annexin A1 | 15 |
| Q9BU<br>Q8 | DDX2<br>3 | Probable ATP-dependent RNA helicase DDX23 | 15 |
| P3811<br>7 | ETFB | Electron transfer flavoprotein subunit beta | 15 |
| P5508<br>4 | HAD<br>HB | Trifunctional enzyme subunit beta, mitochondrial | 15 |
| O1552<br>3 | DDX3<br>Y | ATP-dependent RNA helicase DDX3Y | 15 |
| Q5BJ<br>F6 | ODF2 | Outer dense fiber protein 2 | 15 |
| Q66G<br>S9 | CEP1<br>35 | Centrosomal protein of 135 kDa | 15 |
| Q8TE<br>P8 | CEP1<br>92 | Centrosomal protein of 192 kDa | 15 |

| | | | |
|---|---|---|---|
| P62995 | TRA2B | Transformer-2 protein homolog beta | 14 |
| P62917 | RPL8 | Large ribosomal subunit protein uL2 | 14 |
| P26368 | U2AF2 | Splicing factor U2AF 65 kDa subunit | 14 |
| P46781 | RPS9 | Small ribosomal subunit protein uS4 | 14 |
| P61978 | HNRNPK | Heterogeneous nuclear ribonucleoprotein K | 14 |
| Q9NVI7 | ATAD3A | ATPase family AAA domain-containing protein 3A | 14 |
| P17858 | PFKL | ATP-dependent 6-phosphofructokinase, liver type | 14 |
| P04350 | TUBB4A | Tubulin beta-4A chain | 14 |
| P16401 | H1-5 | Histone H1.5 | |
| P61313 | RPL15 | Large ribosomal subunit protein eL15 | 13 |
| P05141 | SLC25A5 | ADP/ATP translocase 2 | 13 |
| Q14498 | RBM39 | RNA-binding protein 39 | 13 |
| P26373 | RPL13 | Large ribosomal subunit protein eL13 | 13 |
| Q16630 | CPSF6 | Cleavage and polyadenylation specificity factor subunit 6 | 13 |
| O75494 | SRSF10 | Serine/arginine-rich splicing factor 10 | 13 |
| Q06830 | PRDX1 | Peroxiredoxin-1 | 13 |
| P62249 | RPS16 | Small ribosomal subunit protein uS9 | 13 |
| P05089 | ARG1 | Arginase-1 | 13 |
| O00160 | MYO1F | Unconventional myosin-If | 13 |
| P20700 | LMNB1 | Lamin-B1 | 13 |
| P14866 | HNRNPL | Heterogeneous nuclear ribonucleoprotein L | 13 |
| O75083 | WDR1 | WD repeat-containing protein 1 | 13 |
| P08727 | KRT19 | Keratin, type I cytoskeletal 19 | 13 |

| | | | |
|---|---|---|---|
| Q96SN8 | CDK5RAP2 | CDK5 regulatory subunit-associated protein 2 | 13 |
| O43772 | SLC25A20 | Mitochondrial carnitine/acylcarnitine carrier protein | 12 |
| P01889 | HLA-B | HLA class I histocompatibility antigen, B alpha chain | 12 |
| P84103 | SRSF3 | Serine/arginine-rich splicing factor 3 | 12 |
| P23528 | CFL1 | Cofilin-1 | 12 |
| P31930 | UQCRC1 | Cytochrome b-c1 complex subunit 1, mitochondrial | 12 |
| Q08554 | DSC1 | Desmocollin-1 | 12 |
| P18124 | RPL7 | Large ribosomal subunit protein uL30 | 12 |
| P17987 | TCP1 | T-complex protein 1 subunit alpha | 12 |
| Q9UI08 | EVL | Ena/VASP-like protein | 12 |
| P31944 | CASP14 | Caspase-14 | 12 |
| P50990 | CCT8 | T-complex protein 1 subunit theta | 12 |
| Q14739 | LBR | Delta(14)-sterol reductase LBR | 12 |
| Q86UX7 | FERMT3 | Fermitin family homolog 3 | 12 |
| Q02040 | AKAP17A | A-kinase anchor protein 17A | 12 |
| P14868 | DARS1 | Aspartate--tRNA ligase, cytoplasmic | 12 |
| P67936 | TPM4 | Tropomyosin alpha-4 chain | 12 |
| P54886 | ALDH18A1 | Delta-1-pyrroline-5-carboxylate synthase | 12 |
| P26038 | MSN | Moesin | 11 |
| Q16629 | SRSF7 | Serine/arginine-rich splicing factor 7 | 11 |
| P53999 | SUB1 | Activated RNA polymerase II transcriptional coactivator p15 | 11 |
| P62826 | RAN | GTP-binding nuclear protein Ran | 11 |
| Q02543 | RPL18A | Large ribosomal subunit protein eL20 | 11 |

| | | | |
|---|---|---|---|
| P0675 3 | TPM3 | Tropomyosin alpha-3 chain | 11 |
| P0407 5 | ALDO A | Fructose-bisphosphate aldolase A | 11 |
| O1504 2 | U2SU RP | U2 snRNP-associated SURP motif-containing protein | 11 |
| P0623 9 | LCK | Tyrosine-protein kinase Lck | 11 |
| Q96P 63 | SERPI NB12 | Serpin B12 | 11 |
| P0484 3 | RPN1 | Dolichyl-diphosphooligosaccharide--protein glycosyltransferase subunit 1 | 11 |
| Q5VT L8 | PRPF 38B | Pre-mRNA-splicing factor 38B | 11 |
| O4380 9 | NUDT 21 | Cleavage and polyadenylation specificity factor subunit 5 | 11 |
| P0862 1 | SNRN P70 | U1 small nuclear ribonucleoprotein 70 kDa | 11 |
| Q8W UM4 | PDCD 6IP | Programmed cell death 6-interacting protein | 11 |
| P4694 0 | IQGA P1 | Ras GTPase-activating-like protein IQGAP1 | 11 |
| Q5SSJ 5 | HP1B P3 | Heterochromatin protein 1-binding protein 3 | 11 |
| P2475 2 | ACAT 1 | Acetyl-CoA acetyltransferase, mitochondrial | 11 |
| Q9NU V9 | GIMA P4 | GTPase IMAP family member 4 | 11 |
| P0855 9 | PDHA 1 | Pyruvate dehydrogenase E1 component subunit alpha, somatic form, mitochondrial | 11 |
| Q9N WH9 | SLTM | SAFB-like transcription modulator | 11 |
| P1784 4 | DDX5 | Probable ATP-dependent RNA helicase DDX5 | 11 |
| Q8TC U4 | ALMS 1 | Centrosome-associated protein ALMS1 | 11 |
| Q96L C7 | SIGL EC10 | Sialic acid-binding Ig-like lectin 10 | 11 |
| Q9NS B2 | KRT8 4 | Keratin, type II cuticular Hb4 | 11 |
| P6224 1 | RPS8 | Small ribosomal subunit protein eS8 | 10 |
| P6275 3 | RPS6 | Small ribosomal subunit protein eS6 | 10 |
| P6324 4 | RACK 1 | Small ribosomal subunit protein RACK1 | 10 |

| | | | |
|---|---|---|---|
| P06748 | NPM1 | Nucleophosmin | 10 |
| P40429 | RPL13A | Large ribosomal subunit protein uL13 | 10 |
| Q13151 | HNRNPA0 | Heterogeneous nuclear ribonucleoprotein A0 | 10 |
| P63104 | YWHAZ | 14-3-3 protein zeta/delta | 10 |
| P18077 | RPL35A | Large ribosomal subunit protein eL33 | 10 |
| P18621 | RPL17 | Large ribosomal subunit protein uL22 | 10 |
| Q05519 | SRSF11 | Serine/arginine-rich splicing factor 11 | 10 |
| Q9Y2Q3 | GSTK1 | Glutathione S-transferase kappa 1 | 10 |
| P53396 | ACLY | ATP-citrate synthase | 10 |
| P62910 | RPL32 | Large ribosomal subunit protein eL32 | 10 |
| P43403 | ZAP70 | Tyrosine-protein kinase ZAP-70 | 10 |
| P51991 | HNRNPA3 | Heterogeneous nuclear ribonucleoprotein A3 | 10 |
| Q13867 | BLMH | Bleomycin hydrolase | 10 |
| Q14566 | MCM6 | DNA replication licensing factor MCM6 | 10 |
| O43390 | HNRNPR | Heterogeneous nuclear ribonucleoprotein R | 10 |
| O95466 | FMNL1 | Formin-like protein 1 | 10 |
| O75390 | CS | Citrate synthase, mitochondrial | 10 |
| P12236 | SLC25A6 | ADP/ATP translocase 3 | 10 |
| P40227 | CCT6A | T-complex protein 1 subunit zeta | 10 |
| P78386 | KRT85 | Keratin, type II cuticular Hb5 | 10 |
| Q14204 | DYNC1H1 | Cytoplasmic dynein 1 heavy chain 1 | 10 |
| Q86XR8 | CEP57 | Centrosomal protein of 57 kDa | 10 |
| P62805 | H4C1 | Histone H4 | 9 |

| | | | |
|---|---|---|---|
| P07737 | PFN1 | Profilin-1 | 9 |
| P46779 | RPL28 | Large ribosomal subunit protein eL28 | 9 |
| P02786 | TFRC | Transferrin receptor protein 1 | 9 |
| P22695 | UQCRC2 | Cytochrome b-c1 complex subunit 2, mitochondrial | 9 |
| P45880 | VDAC2 | Voltage-dependent anion-selective channel protein 2 | 9 |
| P62750 | RPL23A | Large ribosomal subunit protein uL23 | 9 |
| P32119 | PRDX2 | Peroxiredoxin-2 | 9 |
| P61254 | RPL26 | Large ribosomal subunit protein uL24 | 9 |
| Q969Q0 | RPL36AL | Ribosomal protein eL42-like | 9 |
| Q9Y277 | VDAC3 | Voltage-dependent anion-selective channel protein 3 | 9 |
| Q9UKV3 | ACIN1 | Apoptotic chromatin condensation inducer in the nucleus | 9 |
| P46777 | RPL5 | Large ribosomal subunit protein uL18 | 9 |
| P31146 | CORO1A | Coronin-1A | 9 |
| Q9NQ29 | LUC7L | Putative RNA-binding protein Luc7-like 1 | 9 |
| P13747 | HLA-E | HLA class I histocompatibility antigen, alpha chain E | 9 |
| Q7Z3Y8 | KRT27 | Keratin, type I cytoskeletal 27 | 9 |
| P11413 | G6PD | Glucose-6-phosphate 1-dehydrogenase | 9 |
| Q03518 | TAP1 | Antigen peptide transporter 1 | 9 |
| Q9BU76 | MMTAG2 | Multiple myeloma tumor-associated protein 2 | 9 |
| Q9P258 | RCC2 | Protein RCC2 | 9 |
| Q7KZF4 | SND1 | Staphylococcal nuclease domain-containing protein 1 | 9 |
| Q8NE71 | ABCF1 | ATP-binding cassette sub-family F member 1 | 9 |
| Q7Z3Z0 | KRT25 | Keratin, type I cytoskeletal 25 | 9 |

| | | | |
|---|---|---|---|
| P19338 | NCL | Nucleolin | 9 |
| Q96I99 | SUCLG2 | Succinate--CoA ligase [GDP-forming] subunit beta, mitochondrial | 9 |
| P11310 | ACADM | Medium-chain specific acyl-CoA dehydrogenase, mitochondrial | 9 |
| P53621 | COPA | Coatomer subunit alpha | 9 |
| O60506 | SYNCRIP | Heterogeneous nuclear ribonucleoprotein Q | 9 |
| P07910 | HNRNPC | Heterogeneous nuclear ribonucleoproteins C1/C2 | 9 |
| P13804 | ETFA | Electron transfer flavoprotein subunit alpha, mitochondrial | 9 |
| P22090 | RPS4Y1 | Small ribosomal subunit protein eS4, Y isoform 1 | 9 |
| P33993 | MCM7 | DNA replication licensing factor MCM7 | 9 |
| P51812 | RPS6KA3 | Ribosomal protein S6 kinase alpha-3 | 9 |
| P83881 | RPL36A | Large ribosomal subunit protein eL42 | 9 |
| Q00610 | CLTC | Clathrin heavy chain 1 | 9 |
| Q5T9A4 | ATAD3B | ATPase family AAA domain-containing protein 3B | 9 |
| P62979 | RPS27A | Ubiquitin-ribosomal protein eS31 fusion protein | 8 |
| Q07020 | RPL18 | Large ribosomal subunit protein eL18 | 8 |
| O60814 | H2BC12 | Histone H2B type 1-K | 8 |
| Q66PJ3 | ARL6IP4 | ADP-ribosylation factor-like protein 6-interacting protein 4 | 8 |
| P04899 | GNAI2 | Guanine nucleotide-binding protein G(i) subunit alpha-2 | 8 |
| P62280 | RPS11 | Small ribosomal subunit protein uS17 | 8 |
| P83731 | RPL24 | Large ribosomal subunit protein eL24 | 8 |
| P19367 | HK1 | Hexokinase-1 | 8 |
| P50914 | RPL14 | Large ribosomal subunit protein eL14 | 8 |
| P62829 | RPL23 | Large ribosomal subunit protein uL14 | 8 |

| | | | |
|---|---|---|---|
| P0670 2 | S100A 9 | Protein S100-A9 | 8 |
| P5259 7 | HNR NPF | Heterogeneous nuclear ribonucleoprotein F | 8 |
| O6044 9 | LY75 | Lymphocyte antigen 75 | 8 |
| P1947 4 | TRIM 21 | E3 ubiquitin-protein ligase TRIM21 | 8 |
| Q9H0 U4 | RAB1 B | Ras-related protein Rab-1B | 8 |
| Q1293 1 | TRAP 1 | Heat shock protein 75 kDa, mitochondrial | 8 |
| P5099 1 | CCT4 | T-complex protein 1 subunit delta | 8 |
| O4329 0 | SART 1 | U4/U6.U5 tri-snRNP-associated protein 1 | 8 |
| P0781 4 | EPRS 1 | Bifunctional glutamate/proline--tRNA ligase | 8 |
| Q1410 3 | HNR NPD | Heterogeneous nuclear ribonucleoprotein D0 | 8 |
| Q9NY F8 | BCLA F1 | Bcl-2-associated transcription factor 1 | 8 |
| P1226 8 | IMPD H2 | Inosine-5'-monophosphate dehydrogenase 2 | 8 |
| P3780 2 | TAGL N2 | Transgelin-2 | 8 |
| Q1669 8 | DECR 1 | 2,4-dienoyl-CoA reductase [(3E)-enoyl-CoA-producing], mitochondrial | 8 |
| P3565 9 | DEK | Protein DEK | 8 |
| Q7Z3 Y7 | KRT2 8 | Keratin, type I cytoskeletal 28 | 8 |
| P6282 0 | RAB1 A | Ras-related protein Rab-1A | 8 |
| O1514 3 | ARPC 1B | Actin-related protein 2/3 complex subunit 1B | 8 |
| Q0253 9 | H1-1 | Histone H1.1 | 8 |
| P5021 3 | IDH3 A | Isocitrate dehydrogenase [NAD] subunit alpha, mitochondrial | 8 |
| Q9BS J8 | ESYT 1 | Extended synaptotagmin-1 | 8 |
| P1117 7 | PDHB | Pyruvate dehydrogenase E1 component subunit beta, mitochondrial | 8 |
| P1194 0 | PABP C1 | Polyadenylate-binding protein 1 | 8 |

| | | | |
|---|---|---|---|
| P42357 | HAL | Histidine ammonia-lyase | 8 |
| Q92608 | DOCK2 | Dedicator of cytokinesis protein 2 | 8 |
| Q9H845 | ACAD9 | Complex I assembly factor ACAD9, mitochondrial | 8 |
| Q9UBC9 | SPRR3 | Small proline-rich protein 3 | 8 |
| P62263 | RPS14 | Small ribosomal subunit protein uS11 | 7 |
| P16070 | CD44 | CD44 antigen | 7 |
| P06127 | CD5 | T-cell surface glycoprotein CD5 | 7 |
| P69905 | HBA1 | Hemoglobin subunit alpha | 7 |
| P81605 | DCD | Dermcidin | 7 |
| P62266 | RPS23 | Small ribosomal subunit protein uS12 | 7 |
| Q01130 | SRSF2 | Serine/arginine-rich splicing factor 2 | 7 |
| Q13595 | TRA2A | Transformer-2 protein homolog alpha | 7 |
| P31943 | HNRNPH1 | Heterogeneous nuclear ribonucleoprotein H | 7 |
| P05204 | HMGN2 | Non-histone chromosomal protein HMG-17 | 7 |
| P26447 | S100A4 | Protein S100-A4 | 7 |
| Q99714 | HSD17B10 | 3-hydroxyacyl-CoA dehydrogenase type-2 | 7 |
| P05107 | ITGB2 | Integrin beta-2 | 7 |
| Q15287 | RNPS1 | RNA-binding protein with serine-rich domain 1 | 7 |
| P61204 | ARF3 | ADP-ribosylation factor 3 | 7 |
| P62328 | TMSB4X | Thymosin beta-4 | 7 |
| Q08170 | SRSF4 | Serine/arginine-rich splicing factor 4 | 7 |
| P61006 | RAB8A | Ras-related protein Rab-8A | 7 |
| P36542 | ATP5F1C | ATP synthase subunit gamma, mitochondrial | 7 |

| | | | |
|---|---|---|---|
| P13010 | XRCC5 | X-ray repair cross-complementing protein 5 | 7 |
| P62244 | RPS15A | Small ribosomal subunit protein uS8 | 7 |
| P14625 | HSP90B1 | Endoplasmin | 7 |
| Q9Y6C9 | MTCH2 | Mitochondrial carrier homolog 2 | 7 |
| P38919 | EIF4A3 | Eukaryotic initiation factor 4A-III | 7 |
| O00148 | DDX39A | ATP-dependent RNA helicase DDX39A | 7 |
| P49756 | RBM25 | RNA-binding protein 25 | 7 |
| P52566 | ARHGDIB | Rho GDP-dissociation inhibitor 2 | 7 |
| P22087 | FBL | rRNA 2'-O-methyltransferase fibrillarin | 7 |
| P49368 | CCT3 | T-complex protein 1 subunit gamma | 7 |
| O00299 | CLIC1 | Chloride intracellular channel protein 1 | 7 |
| P07384 | CAPN1 | Calpain-1 catalytic subunit | 7 |
| O43143 | DHX15 | ATP-dependent RNA helicase DHX15 | 7 |
| O94925 | GLS | Glutaminase kidney isoform, mitochondrial | 7 |
| P18085 | ARF4 | ADP-ribosylation factor 4 | 7 |
| P22314 | UBA1 | Ubiquitin-like modifier-activating enzyme 1 | 7 |
| P27144 | AK4 | Adenylate kinase 4, mitochondrial | 7 |
| P29508 | SERPINB3 | Serpin B3 | 7 |
| P49736 | MCM2 | DNA replication licensing factor MCM2 | 7 |
| Q13838 | DDX39B | Spliceosome RNA helicase DDX39B | 7 |
| Q15323 | KRT31 | Keratin, type I cuticular Ha1 | 7 |
| Q9UKM9 | RALY | RNA-binding protein Raly | 7 |
| P35658 | NUP214 | Nuclear pore complex protein Nup214 | 6 |

| | | | |
|---|---|---|---|
| P15153 | RAC2 | Ras-related C3 botulinum toxin substrate 2 | 6 |
| P62913 | RPL11 | Large ribosomal subunit protein uL5 | 6 |
| Q15365 | PCBP1 | Poly(rC)-binding protein 1 | 6 |
| P84098 | RPL19 | Large ribosomal subunit protein eL19 | 6 |
| P27635 | RPL10 | Large ribosomal subunit protein uL16 | 6 |
| Q8IYB3 | SRRM1 | Serine/arginine repetitive matrix protein 1 | 6 |
| P31942 | HNRNPH3 | Heterogeneous nuclear ribonucleoprotein H3 | 6 |
| P46776 | RPL27A | Large ribosomal subunit protein uL15 | 6 |
| P32969 | RPL9 | Large ribosomal subunit protein uL6 | 6 |
| Q13761 | RUNX3 | Runt-related transcription factor 3 | 6 |
| P68871 | HBB | Hemoglobin subunit beta | 6 |
| Q99623 | PHB2 | Prohibitin-2 | 6 |
| P27348 | YWHAQ | 14-3-3 protein theta | 6 |
| P05556 | ITGB1 | Integrin beta-1 | 6 |
| P15880 | RPS2 | Small ribosomal subunit protein uS5 | 6 |
| P46778 | RPL21 | Large ribosomal subunit protein eL21 | 6 |
| Q8WVK2 | SNRNP27 | U4/U6.U5 small nuclear ribonucleoprotein 27 kDa protein | 6 |
| P84095 | RHOG | Rho-related GTP-binding protein RhoG | 6 |
| Q96QA5 | GSDMA | Gasdermin-A | 6 |
| Q8WVV4 | POF1B | Protein POF1B | 6 |
| P62937 | PPIA | Peptidyl-prolyl cis-trans isomerase A | 6 |
| Q14019 | COTL1 | Coactosin-like protein | 6 |
| P78371 | CCT2 | T-complex protein 1 subunit beta | 6 |

| | | | |
|---|---|---|---|
| P84085 | ARF5 | ADP-ribosylation factor 5 | 6 |
| P27824 | CANX | Calnexin | 6 |
| O75400 | PRPF40A | Pre-mRNA-processing factor 40 homolog A | 6 |
| P05109 | S100A8 | Protein S100-A8 | 6 |
| O00154 | ACOT7 | Cytosolic acyl coenzyme A thioester hydrolase | 6 |
| Q9BVP2 | GNL3 | Guanine nucleotide-binding protein-like 3 | 6 |
| P53597 | SUCLG1 | Succinate--CoA ligase [ADP/GDP-forming] subunit alpha, mitochondrial | 6 |
| P27708 | CAD | Multifunctional protein CAD | 6 |
| Q9HCC0 | MCCC2 | Methylcrotonoyl-CoA carboxylase beta chain, mitochondrial | 6 |
| Q8IX12 | CCAR1 | Cell division cycle and apoptosis regulator protein 1 | 6 |
| P15311 | EZR | Ezrin | 6 |
| P31946 | YWHAB | 14-3-3 protein beta/alpha | 6 |
| O43837 | IDH3B | Isocitrate dehydrogenase [NAD] subunit beta, mitochondrial | 6 |
| O95299 | NDUFA10 | NADH dehydrogenase [ubiquinone] 1 alpha subcomplex subunit 10, mitochondrial | 6 |
| P30048 | PRDX3 | Thioredoxin-dependent peroxide reductase, mitochondrial | 6 |
| P35900 | KRT20 | Keratin, type I cytoskeletal 20 | 6 |
| P42765 | ACAA2 | 3-ketoacyl-CoA thiolase, mitochondrial | 6 |
| P51159 | RAB27A | Ras-related protein Rab-27A | 6 |
| Q13523 | PRP4K | Serine/threonine-protein kinase PRP4 homolog | 6 |
| Q13576 | IQGAP2 | Ras GTPase-activating-like protein IQGAP2 | 6 |
| Q14525 | KRT33B | Keratin, type I cuticular Ha3-II | 6 |
| Q14574 | DSC3 | Desmocollin-3 | 6 |
| Q15424 | SAFB | Scaffold attachment factor B1 | 6 |

| | | | |
|---|---|---|---|
| Q16822 | PCK2 | Phosphoenolpyruvate carboxykinase [GTP], mitochondrial | 6 |
| Q6UB35 | MTHFD1L | Monofunctional C1-tetrahydrofolate synthase, mitochondrial | 6 |
| Q9NSE4 | IARS2 | Isoleucine--tRNA ligase, mitochondrial | 6 |
| O95571 | ETHE1 | Persulfide dioxygenase ETHE1, mitochondrial | 5 |
| Q01081 | U2AF1 | Splicing factor U2AF 35 kDa subunit | 5 |
| P62861 | FAU | Ubiquitin-like FUBI-ribosomal protein eS30 fusion protein | 5 |
| Q15517 | CDSN | Corneodesmosin | 5 |
| P09382 | LGALS1 | Galectin-1 | 5 |
| P0C0S8 | H2AC11 | Histone H2A type 1 | 5 |
| P10599 | TXN | Thioredoxin | 5 |
| P27487 | DPP4 | Dipeptidyl peptidase 4 | 5 |
| P02790 | HPX | Hemopexin | 5 |
| Q969P0 | IGSF8 | Immunoglobulin superfamily member 8 | 5 |
| P13612 | ITGA4 | Integrin alpha-4 | 5 |
| Q8TA86 | RP9 | Retinitis pigmentosa 9 protein | 5 |
| P50416 | CPT1A | Carnitine O-palmitoyltransferase 1, liver isoform | 5 |
| P79483 | HLA-DRB3 | HLA class II histocompatibility antigen, DR beta 3 chain | 5 |
| P20073 | ANXA7 | Annexin A7 | 5 |
| P46782 | RPS5 | Small ribosomal subunit protein uS7 | 5 |
| P48643 | CCT5 | T-complex protein 1 subunit epsilon | 5 |
| P00338 | LDHA | L-lactate dehydrogenase A chain | 5 |
| P07195 | LDHB | L-lactate dehydrogenase B chain | 5 |
| Q8WXA9 | SREK1 | Splicing regulatory glutamine/lysine-rich protein 1 | 5 |

| | | | |
|---|---|---|---|
| P51148 | RAB5C | Ras-related protein Rab-5C | 5 |
| Q15366 | PCBP2 | Poly(rC)-binding protein 2 | 5 |
| P17096 | HMGA1 | High mobility group protein HMG-I/HMG-Y | 5 |
| P68431 | H3C1 | Histone H3.1 | 5 |
| Q15717 | ELAVL1 | ELAV-like protein 1 | 5 |
| Q13242 | SRSF9 | Serine/arginine-rich splicing factor 9 | 5 |
| P26583 | HMGB2 | High mobility group protein B2 | 5 |
| P08754 | GNAI3 | Guanine nucleotide-binding protein G(i) subunit alpha-3 | 5 |
| P63241 | EIF5A | Eukaryotic translation initiation factor 5A-1 | 5 |
| Q6ZVX7 | NCCRP1 | F-box only protein 50 | 5 |
| P09429 | HMGB1 | High mobility group protein B1 | 5 |
| P42229 | STAT5A | Signal transducer and activator of transcription 5A | 5 |
| P35998 | PSMC2 | 26S proteasome regulatory subunit 7 | 5 |
| Q92945 | KHSRP | Far upstream element-binding protein 2 | 5 |
| Q14152 | EIF3A | Eukaryotic translation initiation factor 3 subunit A | 5 |
| P55809 | OXCT1 | Succinyl-CoA:3-ketoacid coenzyme A transferase 1, mitochondrial | 5 |
| P50570 | DNM2 | Dynamin-2 | 5 |
| P26641 | EEF1G | Elongation factor 1-gamma | 5 |
| O60234 | GMFG | Glia maturation factor gamma | 5 |
| Q13427 | PPIG | Peptidyl-prolyl cis-trans isomerase G | 5 |
| O00567 | NOP56 | Nucleolar protein 56 | 5 |
| Q13263 | TRIM28 | Transcription intermediary factor 1-beta | 5 |
| O43242 | PSMD3 | 26S proteasome non-ATPase regulatory subunit 3 | 5 |

| | | | |
|---|---|---|---|
| O7534<br>2 | ALOX<br>12B | Arachidonate 12-lipoxygenase, 12R-type | 5 |
| O9480<br>4 | STK1<br>0 | Serine/threonine-protein kinase 10 | 5 |
| P3004<br>1 | PRDX<br>6 | Peroxiredoxin-6 | 5 |
| P3193<br>9 | ATIC | Bifunctional purine biosynthesis protein ATIC | 5 |
| P3194<br>7 | SFN | 14-3-3 protein sigma | 5 |
| P3194<br>8 | STIP1 | Stress-induced-phosphoprotein 1 | 5 |
| P4606<br>3 | RECQ<br>L | ATP-dependent DNA helicase Q1 | 5 |
| P5114<br>9 | RAB7<br>A | Ras-related protein Rab-7a | 5 |
| P5361<br>8 | COPB<br>1 | Coatomer subunit beta | 5 |
| P5999<br>8 | ARPC<br>4 | Actin-related protein 2/3 complex subunit 4 | 5 |
| P6110<br>6 | RAB1<br>4 | Ras-related protein Rab-14 | 5 |
| P6198<br>1 | YWH<br>AG | 14-3-3 protein gamma | 5 |
| Q1324<br>7 | SRSF<br>6 | Serine/arginine-rich splicing factor 6 | 5 |
| Q1354<br>7 | HDA<br>C1 | Histone deacetylase 1 | 5 |
| Q1453<br>2 | KRT3<br>2 | Keratin, type I cuticular Ha2 | 5 |
| Q9259<br>8 | HSPH<br>1 | Heat shock protein 105 kDa | 5 |
| Q9276<br>4 | KRT3<br>5 | Keratin, type I cuticular Ha5 | 5 |
| Q9292<br>8 | RAB1<br>C | Putative Ras-related protein Rab-1C | 5 |
| Q9293<br>0 | RAB8<br>B | Ras-related protein Rab-8B | 5 |
| Q96D<br>N5 | TBC1<br>D31 | TBC1 domain family member 31 | 5 |
| Q9BR<br>D0 | BUD1<br>3 | BUD13 homolog | 5 |
| Q9H2<br>U2 | PPA2 | Inorganic pyrophosphatase 2, mitochondrial | 5 |
| Q9NS<br>B4 | KRT8<br>2 | Keratin, type II cuticular Hb2 | 5 |

| | | | |
|---|---|---|---|
| Q9Y2X3 | NOP58 | Nucleolar protein 58 | 5 |
| P47914 | RPL29 | Large ribosomal subunit protein eL29 | 4 |
| P02652 | APOA2 | Apolipoprotein A-II | 4 |
| Q92522 | H1-10 | Histone H1.10 | 4 |
| Q8N9Q2 | SREK1IP1 | Protein SREK1IP1 | 4 |
| P61927 | RPL37 | Large ribosomal subunit protein eL37 | 4 |
| P06454 | PTMA | Prothymosin alpha | 4 |
| P08195 | SLC3A2 | Amino acid transporter heavy chain SLC3A2 | 4 |
| Q9Y388 | RBMX2 | RNA-binding motif protein, X-linked 2 | 4 |
| P04217 | A1BG | Alpha-1B-glycoprotein | 4 |
| Q9NYJ8 | TAB2 | TGF-beta-activated kinase 1 and MAP3K7-binding protein 2 | 4 |
| Q96CV9 | OPTN | Optineurin | 4 |
| Q9H9B4 | SFXN1 | Sideroflexin-1 | 4 |
| P22234 | PAICS | Bifunctional phosphoribosylaminoimidazole carboxylase/phosphoribosylaminoimidazole succinocarboxamide synthetase | 4 |
| P61224 | RAP1B | Ras-related protein Rap-1b | 4 |
| Q9Y5A9 | YTHDF2 | YTH domain-containing family protein 2 | 4 |
| O95864 | FADS2 | Acyl-CoA 6-desaturase | 4 |
| Q00325 | SLC25A3 | Solute carrier family 25 member 3 | 4 |
| P62851 | RPS25 | Small ribosomal subunit protein eS25 | 4 |
| O60427 | FADS1 | Acyl-CoA (8-3)-desaturase | 4 |
| Q9NZB2 | FAM120A | Constitutive coactivator of PPAR-gamma-like protein 1 | 4 |
| Q9H307 | PNN | Pinin | 4 |

| | | | |
|---|---|---|---|
| P01903 | HLA-DRA | HLA class II histocompatibility antigen, DR alpha chain | 4 |
| O00303 | EIF3F | Eukaryotic translation initiation factor 3 subunit F | 4 |
| Q86V81 | ALYREF | THO complex subunit 4 | 4 |
| Q01469 | FABP5 | Fatty acid-binding protein 5 | 4 |
| P27797 | CALR | Calreticulin | 4 |
| Q96AE4 | FUBP1 | Far upstream element-binding protein 1 | 4 |
| Q03519 | TAP2 | Antigen peptide transporter 2 | 4 |
| Q5TEC6 | H3-7 | Histone H3-7 | 4 |
| P61513 | RPL37A | Large ribosomal subunit protein eL43 | 4 |
| P63313 | TMSB10 | Thymosin beta-10 | 4 |
| B5ME19 | EIF3CL | Eukaryotic translation initiation factor 3 subunit C-like protein | 4 |
| Q13557 | CAMK2D | Calcium/calmodulin-dependent protein kinase type II subunit delta | 4 |
| P62269 | RPS18 | Small ribosomal subunit protein uS13 | 4 |
| Q8WUA2 | PPIL4 | Peptidyl-prolyl cis-trans isomerase-like 4 | 4 |
| P41240 | CSK | Tyrosine-protein kinase CSK | 4 |
| Q92804 | TAF15 | TATA-binding protein-associated factor 2N | 4 |
| Q9Y520 | PRRC2C | Protein PRRC2C | 4 |
| P61026 | RAB10 | Ras-related protein Rab-10 | 4 |
| Q9UQ80 | PA2G4 | Proliferation-associated protein 2G4 | 4 |
| Q9P2E9 | RRBP1 | Ribosome-binding protein 1 | 4 |
| O95747 | OXSR1 | Serine/threonine-protein kinase OSR1 | 4 |
| Q92918 | MAP4K1 | Mitogen-activated protein kinase kinase kinase kinase 1 | 4 |
| P12956 | XRCC6 | X-ray repair cross-complementing protein 6 | 4 |

| | | | |
|---|---|---|---|
| P3974 8 | FEN1 | Flap endonuclease 1 | 4 |
| P4349 0 | NAM PT | Nicotinamide phosphoribosyltransferase | 4 |
| Q9Y3 U8 | RPL3 6 | Large ribosomal subunit protein eL36 | 4 |
| Q9NR 30 | DDX2 1 | Nucleolar RNA helicase 2 | 4 |
| O0023 2 | PSMD 12 | 26S proteasome non-ATPase regulatory subunit 12 | 4 |
| O1538 2 | BCAT 2 | Branched-chain-amino-acid aminotransferase, mitochondrial | 4 |
| O4370 7 | ACTN 4 | Alpha-actinin-4 | 4 |
| O7522 3 | GGCT | Gamma-glutamylcyclotransferase | 4 |
| O7601 3 | KRT3 6 | Keratin, type I cuticular Ha6 | 4 |
| P0408 0 | CSTB | Cystatin-B | 4 |
| P0418 1 | OAT | Ornithine aminotransferase, mitochondrial | 4 |
| P0479 2 | HSPB 1 | Heat shock protein beta-1 | 4 |
| P0733 9 | CTSD | Cathepsin D | 4 |
| P0774 1 | APRT | Adenine phosphoribosyltransferase | 4 |
| P1051 5 | DLAT | Dihydrolipoyllysine-residue acetyltransferase component of pyruvate dehydrogenase complex, mitochondrial | 4 |
| P2328 4 | PPIB | Peptidyl-prolyl cis-trans isomerase B | 4 |
| P2352 6 | AHC Y | Adenosylhomocysteinase | 4 |
| P2520 5 | MCM 3 | DNA replication licensing factor MCM3 | 4 |
| P2531 1 | AZGP 1 | Zinc-alpha-2-glycoprotein | 4 |
| P3004 4 | PRDX 5 | Peroxiredoxin-5, mitochondrial | 4 |
| P3008 4 | ECHS 1 | Enoyl-CoA hydratase, mitochondrial | 4 |
| P3083 7 | ALDH 1B1 | Aldehyde dehydrogenase X, mitochondrial | 4 |

| | | | |
|---|---|---|---|
| P33992 | MCM5 | DNA replication licensing factor MCM5 | 4 |
| P42167 | TMPO | Lamina-associated polypeptide 2, isoforms beta/gamma | 4 |
| P42766 | RPL35 | Large ribosomal subunit protein uL29 | 4 |
| P43897 | TSFM | Elongation factor Ts, mitochondrial | 4 |
| P45973 | CBX5 | Chromobox protein homolog 5 | 4 |
| P47756 | CAPZB | F-actin-capping protein subunit beta | 4 |
| P50552 | VASP | Vasodilator-stimulated phosphoprotein | 4 |
| P50995 | ANXA11 | Annexin A11 | 4 |
| P52565 | ARHGDIA | Rho GDP-dissociation inhibitor 1 | 4 |
| P60891 | PRPS1 | Ribose-phosphate pyrophosphokinase 1 | 4 |
| Q13185 | CBX3 | Chromobox protein homolog 3 | 4 |
| Q16836 | HADH | Hydroxyacyl-coenzyme A dehydrogenase, mitochondrial | 4 |
| Q1KMD3 | HNRNPUL2 | Heterogeneous nuclear ribonucleoprotein U-like protein 2 | 4 |
| Q6PI48 | DARS2 | Aspartate--tRNA ligase, mitochondrial | 4 |
| Q8TC44 | POC1B | POC1 centriolar protein homolog B | 4 |
| Q8WYJ6 | SEPTIN1 | Septin-1 | 4 |
| Q92619 | ARHGAP45 | Rho GTPase-activating protein 45 | 4 |
| Q92947 | GCDH | Glutaryl-CoA dehydrogenase, mitochondrial | 4 |
| Q96RP9 | GFM1 | Elongation factor G, mitochondrial | 4 |
| Q99497 | PARK7 | Parkinson disease protein 7 | 4 |
| Q9BR77 | CCDC77 | Coiled-coil domain-containing protein 77 | 4 |
| Q9P2R7 | SUCLA2 | Succinate--CoA ligase [ADP-forming] subunit beta, mitochondrial | 4 |

| | | | |
|---|---|---|---|
| Q9UH99 | SUN2 | SUN domain-containing protein 2 | 4 |
| Q9UK58 | CCNL1 | Cyclin-L1 | 4 |
| Q9Y305 | ACOT9 | Acyl-coenzyme A thioesterase 9, mitochondrial | 4 |
| Q9Y3I0 | RTCB | RNA-splicing ligase RtcB homolog | 4 |
| Q9Y4W6 | AFG3L2 | Mitochondrial inner membrane m-AAA protease component AFG3L2 | 4 |
| Q5T749 | KPRP | Keratinocyte proline-rich protein | 3 |
| P07477 | PRSS1 | Serine protease 1 | 3 |
| P62854 | RPS26 | Small ribosomal subunit protein eS26 | 3 |
| Q01650 | SLC7A5 | Large neutral amino acids transporter small subunit 1 | 3 |
| Q15388 | TOMM20 | Mitochondrial import receptor subunit TOM20 homolog | 3 |
| P01732 | CD8A | T-cell surface glycoprotein CD8 alpha chain | 3 |
| Q9NZT1 | CALML5 | Calmodulin-like protein 5 | 3 |
| P31949 | S100A11 | Protein S100-A11 | 3 |
| P11717 | IGF2R | Cation-independent mannose-6-phosphate receptor | 3 |
| Q6UWP8 | SBSN | Suprabasin | 3 |
| P08708 | RPS17 | Small ribosomal subunit protein eS17 | 3 |
| O95758 | PTBP3 | Polypyrimidine tract-binding protein 3 | 3 |
| Q71UM5 | RPS27L | Ribosomal protein eS27-like | 3 |
| P08648 | ITGA5 | Integrin alpha-5 | 3 |
| P60660 | MYL6 | Myosin light polypeptide 6 | 3 |
| P60866 | RPS20 | Small ribosomal subunit protein uS10 | 3 |
| P01911 | HLA-DRB1 | HLA class II histocompatibility antigen, DRB1 beta chain | 3 |
| Q13148 | TARDBP | TAR DNA-binding protein 43 | 3 |

| | | | |
|---|---|---|---|
| Q9NVE7 | PANK4 | 4'-phosphopantetheine phosphatase | 3 |
| Q7Z2W4 | ZC3HAV1 | Zinc finger CCCH-type antiviral protein 1 | 3 |
| P49207 | RPL34 | Large ribosomal subunit protein eL34 | 3 |
| P05198 | EIF2S1 | Eukaryotic translation initiation factor 2 subunit 1 | 3 |
| P67809 | YBX1 | Y-box-binding protein 1 | 3 |
| Q6UN15 | FIP1L1 | Pre-mRNA 3'-end-processing factor FIP1 | 3 |
| Q9Y262 | EIF3L | Eukaryotic translation initiation factor 3 subunit L | 3 |
| Q9UJU6 | DBNL | Drebrin-like protein | 3 |
| P61626 | LYZ | Lysozyme C | 3 |
| Q9H299 | SH3BGRL3 | SH3 domain-binding glutamic acid-rich-like protein 3 | 3 |
| Q8NC51 | SERBP1 | SERPINE1 mRNA-binding protein 1 | 3 |
| P47929 | LGALS7 | Galectin-7 | 3 |
| P46977 | STT3A | Dolichyl-diphosphooligosaccharide--protein glycosyltransferase subunit STT3A | 3 |
| P04844 | RPN2 | Dolichyl-diphosphooligosaccharide--protein glycosyltransferase subunit 2 | 3 |
| O75828 | CBR3 | Carbonyl reductase [NADPH] 3 | 3 |
| P05388 | RPLP0 | Large ribosomal subunit protein uL10 | 3 |
| Q9Y2W1 | THRAP3 | Thyroid hormone receptor-associated protein 3 | 3 |
| O76094 | SRP72 | Signal recognition particle subunit SRP72 | 3 |
| P61586 | RHOA | Transforming protein RhoA | 3 |
| Q99729 | HNRNPAB | Heterogeneous nuclear ribonucleoprotein A/B | 3 |
| P29350 | PTPN6 | Tyrosine-protein phosphatase non-receptor type 6 | 3 |
| O43175 | PHGDH | D-3-phosphoglycerate dehydrogenase | 3 |
| O15427 | SLC16A3 | Monocarboxylate transporter 4 | 3 |

| | | | |
|---|---|---|---|
| P26599 | PTBP1 | Polypyrimidine tract-binding protein 1 | 3 |
| Q14974 | KPNB1 | Importin subunit beta-1 | 3 |
| P60903 | S100A10 | Protein S100-A10 | 3 |
| P62277 | RPS13 | Small ribosomal subunit protein uS15 | 3 |
| O75874 | IDH1 | Isocitrate dehydrogenase [NADP] cytoplasmic | 3 |
| Q06210 | GFPT1 | Glutamine--fructose-6-phosphate aminotransferase [isomerizing] 1 | 3 |
| P30153 | PPP2R1A | Serine/threonine-protein phosphatase 2A 65 kDa regulatory subunit A alpha isoform | 3 |
| P01040 | CSTA | Cystatin-A | 3 |
| Q96FW1 | OTUB1 | Ubiquitin thioesterase OTUB1 | 3 |
| Q9NTJ3 | SMC4 | Structural maintenance of chromosomes protein 4 | 3 |
| P30414 | NKTR | NK-tumor recognition protein | 3 |
| P49790 | NUP153 | Nuclear pore complex protein Nup153 | 3 |
| P60174 | TPI1 | Triosephosphate isomerase | 3 |
| A0A8I5KQE6 | RPSA2 | Small ribosomal subunit protein uS2B | 3 |
| P06744 | GPI | Glucose-6-phosphate isomerase | 3 |
| P62847 | RPS24 | Small ribosomal subunit protein eS24 | 3 |
| Q15286 | RAB35 | Ras-related protein Rab-35 | 3 |
| Q13200 | PSMD2 | 26S proteasome non-ATPase regulatory subunit 2 | 3 |
| Q9HB71 | CACYBP | Calcyclin-binding protein | 3 |
| Q15027 | ACAP1 | Arf-GAP with coiled-coil, ANK repeat and PH domain-containing protein 1 | 3 |
| O14818 | PSMA7 | Proteasome subunit alpha type-7 | 3 |
| O14950 | MYL12B | Myosin regulatory light chain 12B | 3 |

| | | | |
|---|---|---|---|
| O15533 | TAPBP | Tapasin | 3 |
| O60264 | SMARCA5 | SWI/SNF-related matrix-associated actin-dependent regulator of chromatin subfamily A member 5 | 3 |
| O75396 | SEC22B | Vesicle-trafficking protein SEC22b | 3 |
| O75439 | PMPCB | Mitochondrial-processing peptidase subunit beta | 3 |
| O94776 | MTA2 | Metastasis-associated protein MTA2 | 3 |
| P00491 | PNP | Purine nucleoside phosphorylase | 3 |
| P01876 | IGHA1 | Immunoglobulin heavy constant alpha 1 | 3 |
| P02452 | COL1A1 | Collagen alpha-1(I) chain | 3 |
| P04279 | SEMG1 | Semenogelin-1 | 3 |
| P04632 | CAPNS1 | Calpain small subunit 1 | 3 |
| P10144 | GZMB | Granzyme B | 3 |
| P15259 | PGAM2 | Phosphoglycerate mutase 2 | 3 |
| P15498 | VAV1 | Proto-oncogene vav | 3 |
| P16219 | ACADS | Short-chain specific acyl-CoA dehydrogenase, mitochondrial | 3 |
| P18583 | SON | Protein SON | 3 |
| P22102 | GART | Trifunctional purine biosynthetic protein adenosine-3 | 3 |
| P25786 | PSMA1 | Proteasome subunit alpha type-1 | 3 |
| P26440 | IVD | Isovaleryl-CoA dehydrogenase, mitochondrial | 3 |
| P30085 | CMPK1 | UMP-CMP kinase | 3 |
| P33991 | MCM4 | DNA replication licensing factor MCM4 | 3 |
| P35637 | FUS | RNA-binding protein FUS | 3 |
| P48047 | ATP5PO | ATP synthase subunit O, mitochondrial | 3 |
| P51114 | FXR1 | RNA-binding protein FXR1 | 3 |

| | | | |
|---|---|---|---|
| P51553 | IDH3G | Isocitrate dehydrogenase [NAD] subunit gamma, mitochondrial | 3 |
| P61158 | ACTR3 | Actin-related protein 3 | 3 |
| P61160 | ACTR2 | Actin-related protein 2 | 3 |
| P61221 | ABCE1 | ATP-binding cassette sub-family E member 1 | 3 |
| P62136 | PPP1CA | Serine/threonine-protein phosphatase PP1-alpha catalytic subunit | 3 |
| P62258 | YWHAE | 14-3-3 protein epsilon | 3 |
| P62879 | GNB2 | Guanine nucleotide-binding protein G(I)/G(S)/G(T) subunit beta-2 | 3 |
| Q01518 | CAP1 | Adenylyl cyclase-associated protein 1 | 3 |
| Q08945 | SSRP1 | FACT complex subunit SSRP1 | 3 |
| Q13428 | TCOF1 | Treacle protein | 3 |
| Q14847 | LASP1 | LIM and SH3 domain protein 1 | 3 |
| Q15126 | PMVK | Phosphomevalonate kinase | 3 |
| Q16610 | ECM1 | Extracellular matrix protein 1 | 3 |
| Q16740 | CLPP | ATP-dependent Clp protease proteolytic subunit, mitochondrial | 3 |
| Q6ZT62 | BARGIN | Bargin | 3 |
| Q86X95 | CIR1 | Corepressor interacting with RBPJ 1 | 3 |
| Q8N4C6 | NIN | Ninein | 3 |
| Q92556 | ELMO1 | Engulfment and cell motility protein 1 | 3 |
| Q92882 | OSTF1 | Osteoclast-stimulating factor 1 | 3 |
| Q96F07 | CYFIP2 | Cytoplasmic FMR1-interacting protein 2 | 3 |
| Q96F15 | GIMAP5 | GTPase IMAP family member 5 | 3 |
| Q96PP8 | GBP5 | Guanylate-binding protein 5 | 3 |
| Q9BRL6 | SRSF8 | Serine/arginine-rich splicing factor 8 | 3 |

| | | | |
|---|---|---|---|
| Q9BXW7 | HDHD5 | Haloacid dehalogenase-like hydrolase domain-containing 5 | 3 |
| Q9H0H5 | RACGAP1 | Rac GTPase-activating protein 1 | 3 |
| Q9H2W6 | MRPL46 | Large ribosomal subunit protein mL46 | 3 |
| Q9NP81 | SARS2 | Serine--tRNA ligase, mitochondrial | 3 |
| Q9NR31 | SAR1A | Small COPII coat GTPase SAR1A | 3 |
| Q9UBG3 | CRNN | Cornulin | 3 |
| Q9UHD8 | SEPTIN9 | Septin-9 | 3 |
| Q9UI42 | CPA4 | Carboxypeptidase A4 | 3 |
| Q9UL46 | PSME2 | Proteasome activator complex subunit 2 | 3 |
| Q9Y5B9 | SUPT16H | FACT complex subunit SPT16 | 3 |
| P06703 | S100A6 | Protein S100-A6 | 2 |
| P61769 | B2M | Beta-2-microglobulin | 2 |
| Q02978 | SLC25A11 | Mitochondrial 2-oxoglutarate/malate carrier protein | 2 |
| P35268 | RPL22 | Large ribosomal subunit protein eL22 | 2 |
| P16150 | SPN | Leukosialin | 2 |
| P02765 | AHSG | Alpha-2-HS-glycoprotein | 2 |
| O60669 | SLC16A7 | Monocarboxylate transporter 2 | 2 |
| Q6UWD8 | C16orf54 | Transmembrane protein C16orf54 | 2 |
| Q6YHK3 | CD109 | CD109 antigen | 2 |
| O95487 | SEC24B | Protein transport protein Sec24B | 2 |
| O00560 | SDCBP | Syntenin-1 | 2 |
| P42677 | RPS27 | Small ribosomal subunit protein eS27 | 2 |
| P30203 | CD6 | T-cell differentiation antigen CD6 | 2 |

| | | | |
|---|---|---|---|
| P3294 2 | ICAM 3 | Intercellular adhesion molecule 3 | 2 |
| Q9Y3 P8 | SIT1 | Signaling threshold-regulating transmembrane adapter 1 | 2 |
| Q1542 8 | SF3A 2 | Splicing factor 3A subunit 2 | 2 |
| P2664 0 | VARS 1 | Valine--tRNA ligase | 2 |
| P0502 3 | ATP1 A1 | Sodium/potassium-transporting ATPase subunit alpha-1 | 2 |
| O6049 6 | DOK2 | Docking protein 2 | 2 |
| P5045 2 | SERPI NB8 | Serpin B8 | 2 |
| Q9Y2 S6 | TMA7 | Translation machinery-associated protein 7 | 2 |
| P4092 6 | MDH 2 | Malate dehydrogenase, mitochondrial | 2 |
| P6289 9 | RPL3 1 | Large ribosomal subunit protein eL31 | 2 |
| O6076 3 | USO1 | General vesicular transport factor p115 | 2 |
| P6309 2 | GNAS | Guanine nucleotide-binding protein G(s) subunit alpha isoforms short | 2 |
| Q8W WP7 | GIMA P1 | GTPase IMAP family member 1 | 2 |
| A6NE C2 | NPEP PSL1 | Puromycin-sensitive aminopeptidase-like protein | 2 |
| P1116 9 | SLC2 A3 | Solute carrier family 2, facilitated glucose transporter member 3 | 2 |
| Q9UH X1 | PUF6 0 | Poly(U)-binding-splicing factor PUF60 | 2 |
| Q1545 9 | SF3A 1 | Splicing factor 3A subunit 1 | 2 |
| Q9288 8 | ARH GEF1 | Rho guanine nucleotide exchange factor 1 | 2 |
| P6195 6 | SUM O2 | Small ubiquitin-related modifier 2 | 2 |
| Q96E P5 | DAZA P1 | DAZ-associated protein 1 | 2 |
| P8410 1 | SERF 2 | Small EDRK-rich factor 2 | 2 |
| P5300 4 | BLVR A | Biliverdin reductase A | 2 |
| Q9NR 56 | MBN L1 | Muscleblind-like protein 1 | 2 |

| | | | |
|---|---|---|---|
| P62195 | PSMC5 | 26S proteasome regulatory subunit 8 | 2 |
| Q8N684 | CPSF7 | Cleavage and polyadenylation specificity factor subunit 7 | 2 |
| P51571 | SSR4 | Translocon-associated protein subunit delta | 2 |
| Q15125 | EBP | 3-beta-hydroxysteroid-Delta(8),Delta(7)-isomerase | 2 |
| Q9UL25 | RAB21 | Ras-related protein Rab-21 | 2 |
| P62857 | RPS28 | Small ribosomal subunit protein eS28 | 2 |
| P27816 | MAP4 | Microtubule-associated protein 4 | 2 |
| O95433 | AHSA1 | Activator of 90 kDa heat shock protein ATPase homolog 1 | 2 |
| Q9BUJ2 | HNRNPUL1 | Heterogeneous nuclear ribonucleoprotein U-like protein 1 | 2 |
| O75791 | GRAP2 | GRB2-related adapter protein 2 | 2 |
| Q04759 | PRKCQ | Protein kinase C theta type | 2 |
| P24666 | ACP1 | Low molecular weight phosphotyrosine protein phosphatase | 2 |
| Q9Y285 | FARSA | Phenylalanine--tRNA ligase alpha subunit | 2 |
| P10747 | CD28 | T-cell-specific surface glycoprotein CD28 | 2 |
| P50749 | RASSF2 | Ras association domain-containing protein 2 | 2 |
| P60900 | PSMA6 | Proteasome subunit alpha type-6 | 2 |
| Q9H0A0 | NAT10 | RNA cytidine acetyltransferase | 2 |
| Q9Y228 | TRAF3IP3 | TRAF3-interacting JNK-activating modulator | 2 |
| P61077 | UBE2D3 | Ubiquitin-conjugating enzyme E2 D3 | 2 |
| Q16891 | IMMT | MICOS complex subunit MIC60 | 2 |
| P50402 | EMD | Emerin | 2 |
| P46013 | MKI67 | Proliferation marker protein Ki-67 | 2 |

| | | | |
|---|---|---|---|
| O94906 | PRPF6 | Pre-mRNA-processing factor 6 | 2 |
| P25788 | PSMA3 | Proteasome subunit alpha type-3 | 2 |
| A0A5B9 | TRBC2 | T cell receptor beta constant 2 | 2 |
| A0FGR8 | ESYT2 | Extended synaptotagmin-2 | 2 |
| C9JLJ4 | USP17L13 | Ubiquitin carboxyl-terminal hydrolase 17-like protein 13 | 2 |
| O00483 | NDUFA4 | Cytochrome c oxidase subunit NDUFA4 | 2 |
| O14617 | AP3D1 | AP-3 complex subunit delta-1 | 2 |
| O14773 | TPP1 | Tripeptidyl-peptidase 1 | 2 |
| O15145 | ARPC3 | Actin-related protein 2/3 complex subunit 3 | 2 |
| O15371 | EIF3D | Eukaryotic translation initiation factor 3 subunit D | 2 |
| O15511 | ARPC5 | Actin-related protein 2/3 complex subunit 5 | 2 |
| O43684 | BUB3 | Mitotic checkpoint protein BUB3 | 2 |
| O60488 | ACSL4 | Long-chain-fatty-acid--CoA ligase 4 | 2 |
| O60610 | DIAPH1 | Protein diaphanous homolog 1 | 2 |
| O60880 | SH2D1A | SH2 domain-containing protein 1A | 2 |
| O75131 | CPNE3 | Copine-3 | 2 |
| O75534 | CSDE1 | Cold shock domain-containing protein E1 | 2 |
| O75663 | TIPRL | TIP41-like protein | 2 |
| P02788 | LTF | Lactotransferrin | 2 |
| P06493 | CDK1 | Cyclin-dependent kinase 1 | 2 |
| P07476 | IVL | Involucrin | 2 |
| P09211 | GSTP1 | Glutathione S-transferase P | 2 |
| P12273 | PIP | Prolactin-inducible protein | 2 |

| | | | |
|---|---|---|---|
| P12694 | BCKDHA | 2-oxoisovalerate dehydrogenase subunit alpha, mitochondrial | 2 |
| P13489 | RNH1 | Ribonuclease inhibitor | 2 |
| P13693 | TPT1 | Translationally-controlled tumor protein | 2 |
| P14174 | MIF | Macrophage migration inhibitory factor | 2 |
| P14222 | PRF1 | Perforin-1 | 2 |
| P14678 | SNRPB | Small nuclear ribonucleoprotein-associated proteins B and B' | 2 |
| P16615 | ATP2A2 | Sarcoplasmic/endoplasmic reticulum calcium ATPase 2 | 2 |
| P17026 | ZNF22 | Zinc finger protein 22 | 2 |
| P17252 | PRKCA | Protein kinase C alpha type | 2 |
| P17931 | LGALS3 | Galectin-3 | 2 |
| P18031 | PTPN1 | Tyrosine-protein phosphatase non-receptor type 1 | 2 |
| P21912 | SDHB | Succinate dehydrogenase [ubiquinone] iron-sulfur subunit, mitochondrial | 2 |
| P23786 | CPT2 | Carnitine O-palmitoyltransferase 2, mitochondrial | 2 |
| P23919 | DTYMK | Thymidylate kinase | 2 |
| P25325 | MPST | 3-mercaptopyruvate sulfurtransferase | 2 |
| P28838 | LAP3 | Cytosol aminopeptidase | 2 |
| P29728 | OAS2 | 2'-5'-oligoadenylate synthase 2 | 2 |
| P31040 | SDHA | Succinate dehydrogenase [ubiquinone] flavoprotein subunit, mitochondrial | 2 |
| P31151 | S100A7 | Protein S100-A7 | 2 |
| P32456 | GBP2 | Guanylate-binding protein 2 | 2 |
| P35606 | COPB2 | Coatomer subunit beta' | 2 |
| P41250 | GARS1 | Glycine--tRNA ligase | 2 |
| P42694 | HELZ | Probable helicase with zinc finger domain | 2 |

| | | | |
|---|---|---|---|
| P43243 | MATR3 | Matrin-3 | 2 |
| P43487 | RANBP1 | Ran-specific GTPase-activating protein | 2 |
| P43652 | AFM | Afamin | 2 |
| P45954 | ACADSB | Short/branched chain specific acyl-CoA dehydrogenase, mitochondrial | 2 |
| P47897 | QARS1 | Glutamine--tRNA ligase | 2 |
| P49406 | MRPL19 | Large ribosomal subunit protein bL19m | 2 |
| P49720 | PSMB3 | Proteasome subunit beta type-3 | 2 |
| P51659 | HSD17B4 | Peroxisomal multifunctional enzyme type 2 | 2 |
| P54136 | RARS1 | Arginine--tRNA ligase, cytoplasmic | 2 |
| P54577 | YARS1 | Tyrosine--tRNA ligase, cytoplasmic | 2 |
| P55072 | VCP | Transitional endoplasmic reticulum ATPase | 2 |
| P57088 | TMEM33 | Transmembrane protein 33 | 2 |
| P60981 | DSTN | Destrin | 2 |
| P62273 | RPS29 | Small ribosomal subunit protein uS14 | 2 |
| P62330 | ARF6 | ADP-ribosylation factor 6 | 2 |
| P62491 | RAB11A | Ras-related protein Rab-11A | 2 |
| P62891 | RPL39 | Large ribosomal subunit protein eL39 | 2 |
| P63167 | DYNLL1 | Dynein light chain 1, cytoplasmic | 2 |
| Q02252 | ALDH6A1 | Methylmalonate-semialdehyde/malonate-semialdehyde dehydrogenase [acylating], mitochondrial | 2 |
| Q02750 | MAP2K1 | Dual specificity mitogen-activated protein kinase kinase 1 | 2 |
| Q04637 | EIF4G1 | Eukaryotic translation initiation factor 4 gamma 1 | 2 |
| Q07666 | KHDRBS1 | KH domain-containing, RNA-binding, signal transduction-associated protein 1 | 2 |
| Q10713 | PMPCA | Mitochondrial-processing peptidase subunit alpha | 2 |

| | | | |
|---|---|---|---|
| Q13045 | FLII | Protein flightless-1 homolog | 2 |
| Q13422 | IKZF1 | DNA-binding protein Ikaros | 2 |
| Q13423 | NNT | NAD(P) transhydrogenase, mitochondrial | 2 |
| Q13951 | CBFB | Core-binding factor subunit beta | 2 |
| Q14761 | PTPRCAP | Protein tyrosine phosphatase receptor type C-associated protein | 2 |
| Q14839 | CHD4 | Chromodomain-helicase-DNA-binding protein 4 | 2 |
| Q15056 | EIF4H | Eukaryotic translation initiation factor 4H | 2 |
| Q15181 | PPA1 | Inorganic pyrophosphatase | 2 |
| Q16555 | DPYSL2 | Dihydropyrimidinase-related protein 2 | 2 |
| Q29RF7 | PDS5A | Sister chromatid cohesion protein PDS5 homolog A | 2 |
| Q53QZ3 | ARHGAP15 | Rho GTPase-activating protein 15 | 2 |
| Q5JXC2 | MIIP | Migration and invasion-inhibitory protein | 2 |
| Q5T750 | KPLCE | Protein KPLCE | 2 |
| Q6IBS0 | TWF2 | Twinfilin-2 | 2 |
| Q6P1X5 | TAF2 | Transcription initiation factor TFIID subunit 2 | 2 |
| Q70J99 | UNC13D | Protein unc-13 homolog D | 2 |
| Q7L0Y3 | TRMT10C | tRNA methyltransferase 10 homolog C | 2 |
| Q7L4I2 | RSRC2 | Arginine/serine-rich coiled-coil protein 2 | 2 |
| Q86YV0 | RASAL3 | RAS protein activator like-3 | 2 |
| Q8N8A2 | ANKRD44 | Serine/threonine-protein phosphatase 6 regulatory ankyrin repeat subunit B | 2 |
| Q8NF50 | DOCK8 | Dedicator of cytokinesis protein 8 | 2 |
| Q8TAD8 | SNIP1 | Smad nuclear-interacting protein 1 | 2 |

| | | | |
|---|---|---|---|
| Q92820 | GGH | Gamma-glutamyl hydrolase | 2 |
| Q96KR1 | ZFR | Zinc finger RNA-binding protein | 2 |
| Q96MT8 | CEP63 | Centrosomal protein of 63 kDa | 2 |
| Q96T37 | RBM15 | RNA-binding protein 15 | 2 |
| Q9BPW8 | NIPSNAP1 | Protein NipSnap homolog 1 | 2 |
| Q9BRT6 | LLPH | Protein LLP homolog | 2 |
| Q9BTT0 | ANP32E | Acidic leucine-rich nuclear phosphoprotein 32 family member E | 2 |
| Q9BZE4 | GTPBP4 | GTP-binding protein 4 | 2 |
| Q9H4M9 | EHD1 | EH domain-containing protein 1 | 2 |
| Q9H7N4 | SCAF1 | Splicing factor, arginine/serine-rich 19 | 2 |
| Q9NQ31 | AKIP1 | A-kinase-interacting protein 1 | 2 |
| Q9NRR4 | DROSHA | Ribonuclease 3 | 2 |
| Q9NTI5 | PDS5B | Sister chromatid cohesion protein PDS5 homolog B | 2 |
| Q9NY33 | DPP3 | Dipeptidyl peptidase 3 | 2 |
| Q9NYK5 | MRPL39 | Large ribosomal subunit protein mL39 | 2 |
| Q9NZ01 | TECR | Very-long-chain enoyl-CoA reductase | 2 |
| Q9P0L0 | VAPA | Vesicle-associated membrane protein-associated protein A | 2 |
| Q9UG63 | ABCF2 | ATP-binding cassette sub-family F member 2 | 2 |
| Q9UHD2 | TBK1 | Serine/threonine-protein kinase TBK1 | 2 |
| Q9UIQ6 | LNPEP | Leucyl-cystinyl aminopeptidase | 2 |
| Q9Y2Z4 | YARS2 | Tyrosine--tRNA ligase, mitochondrial | 2 |
| Q9Y3F4 | STRAP | Serine-threonine kinase receptor-associated protein | 2 |
| Q9Y697 | NFS1 | Cysteine desulfurase | 2 |

| | | | |
|---|---|---|---|
| Q8TF72 | SHROOM3 | Protein Shroom3 | 1 |
| P02656 | APOC3 | Apolipoprotein C-III | 1 |
| O94762 | RECQL5 | ATP-dependent DNA helicase Q5 | 1 |
| Q8TCT9 | HM13 | Minor histocompatibility antigen H13 | 1 |
| Q96S97 | MYADM | Myeloid-associated differentiation marker | 1 |
| P33176 | KIF5B | Kinesin-1 heavy chain | 1 |
| Q10589 | BST2 | Bone marrow stromal antigen 2 | 1 |
| Q32MZ4 | LRRFIP1 | Leucine-rich repeat flightless-interacting protein 1 | 1 |
| P35030 | PRSS3 | Trypsin-3 | 1 |
| P61964 | WDR5 | WD repeat-containing protein 5 | 1 |
| P04156 | PRNP | Major prion protein | 1 |
| O15269 | SPTLC1 | Serine palmitoyltransferase 1 | 1 |
| P13164 | IFITM1 | Interferon-induced transmembrane protein 1 | 1 |
| P05387 | RPLP2 | Large ribosomal subunit protein P2 | 1 |
| P30536 | TSPO | Translocator protein | 1 |
| Q9UBM7 | DHCR7 | 7-dehydrocholesterol reductase | 1 |
| P04818 | TYMS | Thymidylate synthase | 1 |
| Q99808 | SLC29A1 | Equilibrative nucleoside transporter 1 | 1 |
| P61088 | UBE2N | Ubiquitin-conjugating enzyme E2 N | 1 |
| P61966 | AP1S1 | AP-1 complex subunit sigma-1A | 1 |
| Q9HCY8 | S100A14 | Protein S100-A14 | 1 |
| Q16186 | ADRM1 | Proteasomal ubiquitin receptor ADRM1 | 1 |
| Q14242 | SELPLG | P-selectin glycoprotein ligand 1 | 1 |

| | | | |
|---|---|---|---|
| Q01844 | EWSR1 | RNA-binding protein EWS | 1 |
| Q08380 | LGALS3BP | Galectin-3-binding protein | 1 |
| O94826 | TOMM70 | Mitochondrial import receptor subunit TOM70 | 1 |
| O95202 | LETM1 | Mitochondrial proton/calcium exchanger protein | 1 |
| P30040 | ERP29 | Endoplasmic reticulum resident protein 29 | 1 |
| Q9BRF8 | CPPED1 | Serine/threonine-protein phosphatase CPPED1 | 1 |
| P28072 | PSMB6 | Proteasome subunit beta type-6 | 1 |
| A6NHR9 | SMCHD1 | Structural maintenance of chromosomes flexible hinge domain-containing protein 1 | 1 |
| P49458 | SRP9 | Signal recognition particle 9 kDa protein | 1 |
| A6NJW9 | CD8B2 | T-cell surface glycoprotein CD8 beta-2 chain | 1 |
| P06729 | CD2 | T-cell surface antigen CD2 | 1 |
| P19256 | CD58 | Lymphocyte function-associated antigen 3 | 1 |
| Q92973 | TNPO1 | Transportin-1 | 1 |
| P05114 | HMGN1 | Non-histone chromosomal protein HMG-14 | 1 |
| P06396 | GSN | Gelsolin | 1 |
| A1KXE4 | FAM168B | Myelin-associated neurite-outgrowth inhibitor | 1 |
| O75153 | CLUH | Clustered mitochondria protein homolog | 1 |
| P48444 | ARCN1 | Coatomer subunit delta | 1 |
| Q9Y3Y2 | CHTOP | Chromatin target of PRMT1 protein | 1 |
| P00414 | MT-CO3 | Cytochrome c oxidase subunit 3 | 1 |
| Q12913 | PTPRJ | Receptor-type tyrosine-protein phosphatase eta | 1 |
| O15294 | OGT | UDP-N-acetylglucosamine--peptide N-acetylglucosaminyltransferase 110 kDa subunit | 1 |
| P60953 | CDC42 | Cell division control protein 42 homolog | 1 |

| | | | |
|---|---|---|---|
| Q9H7M9 | VSIR | V-type immunoglobulin domain-containing suppressor of T-cell activation | 1 |
| P51610 | HCFC1 | Host cell factor 1 | 1 |
| P63173 | RPL38 | Large ribosomal subunit protein eL38 | 1 |
| P48729 | CSNK1A1 | Casein kinase I isoform alpha | 1 |
| Q86UE4 | MTDH | Protein LYRIC | 1 |
| P22061 | PCMT1 | Protein-L-isoaspartate(D-aspartate) O-methyltransferase | 1 |
| Q9Y266 | NUDC | Nuclear migration protein nudC | 1 |
| Q92621 | NUP205 | Nuclear pore complex protein Nup205 | 1 |
| E9PAV3 | NACA | Nascent polypeptide-associated complex subunit alpha, muscle-specific form | 1 |
| P33241 | LSP1 | Lymphocyte-specific protein 1 | 1 |
| Q92854 | SEMA4D | Semaphorin-4D | 1 |
| Q86VP6 | CAND1 | Cullin-associated NEDD8-dissociated protein 1 | 1 |
| Q7Z4W1 | DCXR | L-xylulose reductase | 1 |
| Q9UPN7 | PPP6R1 | Serine/threonine-protein phosphatase 6 regulatory subunit 1 | 1 |
| Q99426 | TBCB | Tubulin-folding cofactor B | 1 |
| P02750 | LRG1 | Leucine-rich alpha-2-glycoprotein | 1 |
| P23368 | ME2 | NAD-dependent malic enzyme, mitochondrial | 1 |
| Q9H0D6 | XRN2 | 5'-3' exoribonuclease 2 | 1 |
| Q14061 | COX17 | Cytochrome c oxidase copper chaperone | 1 |
| Q8NDC0 | MAPK1IP1L | MAPK-interacting and spindle-stabilizing protein-like | 1 |
| Q9Y2L1 | DIS3 | Exosome complex exonuclease RRP44 | 1 |
| Q86UU0 | BCL9L | B-cell CLL/lymphoma 9-like protein | 1 |

| | | | |
|---|---|---|---|
| P22033 | MMUT | Methylmalonyl-CoA mutase, mitochondrial | 1 |
| A0A0B4J2D5 | GATD3B | Putative glutamine amidotransferase-like class 1 domain-containing protein 3B, mitochondrial | 1 |
| A6NFQ2 | TCAF2 | TRPM8 channel-associated factor 2 | 1 |
| A6NKT7 | RGPD3 | RanBP2-like and GRIP domain-containing protein 3 | 1 |
| O00116 | AGPS | Alkyldihydroxyacetonephosphate synthase, peroxisomal | 1 |
| O00139 | KIF2A | Kinesin-like protein KIF2A | 1 |
| O00231 | PSMD11 | 26S proteasome non-ATPase regulatory subunit 11 | 1 |
| O14602 | EIF1AY | Eukaryotic translation initiation factor 1A, Y-chromosomal | 1 |
| O14745 | NHERF1 | Na(+)/H(+) exchange regulatory cofactor NHE-RF1 | 1 |
| O14880 | MGST3 | Glutathione S-transferase 3, mitochondrial | 1 |
| O14980 | XPO1 | Exportin-1 | 1 |
| O15173 | PGRMC2 | Membrane-associated progesterone receptor component 2 | 1 |
| O15260 | SURF4 | Surfeit locus protein 4 | 1 |
| O15355 | PPM1G | Protein phosphatase 1G | 1 |
| O15372 | EIF3H | Eukaryotic translation initiation factor 3 subunit H | 1 |
| O43548 | TGM5 | Protein-glutamine gamma-glutamyltransferase 5 | 1 |
| O43615 | TIMM44 | Mitochondrial import inner membrane translocase subunit TIM44 | 1 |
| O75164 | KDM4A | Lysine-specific demethylase 4A | 1 |
| O75306 | NDUFS2 | NADH dehydrogenase [ubiquinone] iron-sulfur protein 2, mitochondrial | 1 |
| O75323 | NIPSNAP2 | Protein NipSnap homolog 2 | 1 |
| O75582 | RPS6KA5 | Ribosomal protein S6 kinase alpha-5 | 1 |
| O75635 | SERPINB7 | Serpin B7 | 1 |

| | | | |
|---|---|---|---|
| O75643 | SNRNP200 | U5 small nuclear ribonucleoprotein 200 kDa helicase | 1 |
| O75821 | EIF3G | Eukaryotic translation initiation factor 3 subunit G | 1 |
| O76031 | CLPX | ATP-dependent Clp protease ATP-binding subunit clpX-like, mitochondrial | 1 |
| O94903 | PLPBP | Pyridoxal phosphate homeostasis protein | 1 |
| O95319 | CELF2 | CUGBP Elav-like family member 2 | 1 |
| O95363 | FARS2 | Phenylalanine--tRNA ligase, mitochondrial | 1 |
| P00387 | CYB5R3 | NADH-cytochrome b5 reductase 3 | 1 |
| P00403 | MT-CO2 | Cytochrome c oxidase subunit 2 | 1 |
| P01591 | JCHAIN | Immunoglobulin J chain | 1 |
| P01833 | PIGR | Polymeric immunoglobulin receptor | 1 |
| P02100 | HBE1 | Hemoglobin subunit epsilon | 1 |
| P04746 | AMY2A | Pancreatic alpha-amylase | 1 |
| P07237 | P4HB | Protein disulfide-isomerase | 1 |
| P07686 | HEXB | Beta-hexosaminidase subunit beta | 1 |
| P07954 | FH | Fumarate hydratase, mitochondrial | 1 |
| P09525 | ANXA4 | Annexin A4 | 1 |
| P09874 | PARP1 | Poly [ADP-ribose] polymerase 1 | 1 |
| P0DOY2 | IGLC2 | Immunoglobulin lambda constant 2 | 1 |
| P10155 | RO60 | RNA-binding protein RO60 | 1 |
| P10619 | CTSA | Lysosomal protective protein | 1 |
| P10644 | PRKAR1A | cAMP-dependent protein kinase type I-alpha regulatory subunit | 1 |
| P11216 | PYGB | Glycogen phosphorylase, brain form | 1 |
| P13995 | MTHFD2 | Bifunctional methylenetetrahydrofolate dehydrogenase/cyclohydrolase, mitochondrial | 1 |

| | | | |
|---|---|---|---|
| P14317 | HCLS1 | Hematopoietic lineage cell-specific protein | 1 |
| P15170 | GSPT1 | Eukaryotic peptide chain release factor GTP-binding subunit ERF3A | 1 |
| P15954 | COX7C | Cytochrome c oxidase subunit 7C, mitochondrial | 1 |
| P17655 | CAPN2 | Calpain-2 catalytic subunit | 1 |
| P22830 | FECH | Ferrochelatase, mitochondrial | 1 |
| P23381 | WARS1 | Tryptophan--tRNA ligase, cytoplasmic | 1 |
| P25398 | RPS12 | Small ribosomal subunit protein eS12 | 1 |
| P25789 | PSMA4 | Proteasome subunit alpha type-4 | 1 |
| P25815 | S100P | Protein S100-P | 1 |
| P26196 | DDX6 | Probable ATP-dependent RNA helicase DDX6 | 1 |
| P27707 | DCK | Deoxycytidine kinase | 1 |
| P28074 | PSMB5 | Proteasome subunit beta type-5 | 1 |
| P29466 | CASP1 | Caspase-1 | 1 |
| P30050 | RPL12 | Large ribosomal subunit protein uL11 | 1 |
| P30086 | PEBP1 | Phosphatidylethanolamine-binding protein 1 | 1 |
| P30101 | PDIA3 | Protein disulfide-isomerase A3 | 1 |
| P30405 | PPIF | Peptidyl-prolyl cis-trans isomerase F, mitochondrial | 1 |
| P31689 | DNAJA1 | DnaJ homolog subfamily A member 1 | 1 |
| P31937 | HIBADH | 3-hydroxyisobutyrate dehydrogenase, mitochondrial | 1 |
| P32320 | CDA | Cytidine deaminase | 1 |
| P32970 | CD70 | CD70 antigen | 1 |
| P40200 | CD96 | T-cell surface protein tactile | 1 |
| P40616 | ARL1 | ADP-ribosylation factor-like protein 1 | 1 |

| | | | |
|---|---|---|---|
| P40938 | RFC3 | Replication factor C subunit 3 | 1 |
| P41091 | EIF2S3 | Eukaryotic translation initiation factor 2 subunit 3 | 1 |
| P41143 | OPRD1 | Delta-type opioid receptor | 1 |
| P41252 | IARS1 | Isoleucine--tRNA ligase, cytoplasmic | 1 |
| P42331 | ARHGAP25 | Rho GTPase-activating protein 25 | 1 |
| P47755 | CAPZA2 | F-actin-capping protein subunit alpha-2 | 1 |
| P48651 | PTDSS1 | Phosphatidylserine synthase 1 | 1 |
| P49189 | ALDH9A1 | 4-trimethylaminobutyraldehyde dehydrogenase | 1 |
| P49257 | LMAN1 | Protein ERGIC-53 | 1 |
| P49588 | AARS1 | Alanine--tRNA ligase, cytoplasmic | 1 |
| P49591 | SARS1 | Serine--tRNA ligase, cytoplasmic | 1 |
| P49773 | HINT1 | Adenosine 5'-monophosphoramidase HINT1 | 1 |
| P49821 | NDUFV1 | NADH dehydrogenase [ubiquinone] flavoprotein 1, mitochondrial | 1 |
| P49841 | GSK3B | Glycogen synthase kinase-3 beta | 1 |
| P50238 | CRIP1 | Cysteine-rich protein 1 | 1 |
| P52292 | KPNA2 | Importin subunit alpha-1 | 1 |
| P53602 | MVD | Diphosphomevalonate decarboxylase | 1 |
| P53634 | CTSC | Dipeptidyl peptidase 1 | 1 |
| P55160 | NCKAP1L | Nck-associated protein 1-like | 1 |
| P55265 | ADAR | Double-stranded RNA-specific adenosine deaminase | 1 |
| P56192 | MARS1 | Methionine--tRNA ligase, cytoplasmic | 1 |
| P56537 | EIF6 | Eukaryotic translation initiation factor 6 | 1 |

| | | | |
|---|---|---|---|
| P58546 | MTPN | Myotrophin | 1 |
| P59768 | GNG2 | Guanine nucleotide-binding protein G(I)/G(S)/G(O) subunit gamma-2 | 1 |
| P61353 | RPL27 | Large ribosomal subunit protein eL27 | 1 |
| P61619 | SEC61A1 | Protein transport protein Sec61 subunit alpha isoform 1 | 1 |
| P62070 | RRAS2 | Ras-related protein R-Ras2 | 1 |
| P62081 | RPS7 | Small ribosomal subunit protein eS7 | 1 |
| P62495 | ETF1 | Eukaryotic peptide chain release factor subunit 1 | 1 |
| P62841 | RPS15 | Small ribosomal subunit protein uS19 | 1 |
| P63151 | PPP2R2A | Serine/threonine-protein phosphatase 2A 55 kDa regulatory subunit B alpha isoform | 1 |
| P63172 | DYNLT1 | Dynein light chain Tctex-type 1 | 1 |
| P68402 | PAFAH1B2 | Platelet-activating factor acetylhydrolase IB subunit alpha2 | 1 |
| P82650 | MRPS22 | Small ribosomal subunit protein mS22 | 1 |
| P82912 | MRPS11 | Small ribosomal subunit protein uS11m | 1 |
| P82921 | MRPS21 | Small ribosomal subunit protein bS21m | 1 |
| P82979 | SARNP | SAP domain-containing ribonucleoprotein | 1 |
| Q00653 | NFKB2 | Nuclear factor NF-kappa-B p100 subunit | 1 |
| Q02338 | BDH1 | D-beta-hydroxybutyrate dehydrogenase, mitochondrial | 1 |
| Q02790 | FKBP4 | Peptidyl-prolyl cis-trans isomerase FKBP4 | 1 |
| Q0P6D6 | CCDC15 | Coiled-coil domain-containing protein 15 | 1 |
| Q12849 | GRSF1 | G-rich sequence factor 1 | 1 |
| Q13043 | STK4 | Serine/threonine-protein kinase 4 | 1 |
| Q13363 | CTBP1 | C-terminal-binding protein 1 | 1 |
| Q13596 | SNX1 | Sorting nexin-1 | 1 |

| | | | |
|---|---|---|---|
| Q14203 | DCTN1 | Dynactin subunit 1 | 1 |
| Q14318 | FKBP8 | Peptidyl-prolyl cis-trans isomerase FKBP8 | 1 |
| Q14562 | DHX8 | ATP-dependent RNA helicase DHX8 | 1 |
| Q14697 | GANAB | Neutral alpha-glucosidase AB | 1 |
| Q15046 | KARS1 | Lysine--tRNA ligase | 1 |
| Q15185 | PTGES3 | Prostaglandin E synthase 3 | 1 |
| Q15628 | TRADD | Tumor necrosis factor receptor type 1-associated DEATH domain protein | 1 |
| Q15646 | OASL | 2'-5'-oligoadenylate synthase-like protein | 1 |
| Q15828 | CST6 | Cystatin-M | 1 |
| Q15833 | STXBP2 | Syntaxin-binding protein 2 | 1 |
| Q16378 | PRR4 | Proline-rich protein 4 | 1 |
| Q16637 | SMN1 | Survival motor neuron protein | 1 |
| Q16762 | TST | Thiosulfate sulfurtransferase | 1 |
| Q16774 | GUK1 | Guanylate kinase | 1 |
| Q16795 | NDUFA9 | NADH dehydrogenase [ubiquinone] 1 alpha subcomplex subunit 9, mitochondrial | 1 |
| Q2M2I8 | AAK1 | AP2-associated protein kinase 1 | 1 |
| Q2TAL8 | QRICH1 | Transcriptional regulator QRICH1 | 1 |
| Q52LJ0 | FAM98B | Protein FAM98B | 1 |
| Q5HYI8 | RABL3 | Rab-like protein 3 | 1 |
| Q5JRX3 | PITRM1 | Presequence protease, mitochondrial | 1 |
| Q5JTV8 | TOR1AIP1 | Torsin-1A-interacting protein 1 | 1 |
| Q6DD88 | ATL3 | Atlastin-3 | 1 |
| Q6P1L8 | MRPL14 | Large ribosomal subunit protein uL14m | 1 |

| | | | |
|---|---|---|---|
| Q6P2E9 | EDC4 | Enhancer of mRNA-decapping protein 4 | 1 |
| Q7L1Q6 | BZW1 | eIF5-mimic protein 2 | 1 |
| Q7L2H7 | EIF3M | Eukaryotic translation initiation factor 3 subunit M | 1 |
| Q86SX6 | GLRX5 | Glutaredoxin-related protein 5, mitochondrial | 1 |
| Q86X76 | NIT1 | Deaminated glutathione amidase | 1 |
| Q8IUI8 | CRLF3 | Cytokine receptor-like factor 3 | 1 |
| Q8IZL8 | PELP1 | Proline-, glutamic acid- and leucine-rich protein 1 | 1 |
| Q8N163 | CCAR2 | Cell cycle and apoptosis regulator protein 2 | 1 |
| Q8N1K5 | THEMIS | Protein THEMIS | 1 |
| Q8N4Q0 | PTGR3 | Prostaglandin reductase 3 | 1 |
| Q8NAV1 | PRPF38A | Pre-mRNA-splicing factor 38A | 1 |
| Q8NI60 | COQ8A | Atypical kinase COQ8A, mitochondrial | 1 |
| Q8TAE8 | GADD45GIP1 | Large ribosomal subunit protein mL64 | 1 |
| Q8TAQ2 | SMARCC2 | SWI/SNF complex subunit SMARCC2 | 1 |
| Q8TC12 | RDH11 | Retinol dehydrogenase 11 | 1 |
| Q8TDD1 | DDX54 | ATP-dependent RNA helicase DDX54 | 1 |
| Q8WU79 | SMAP2 | Stromal membrane-associated protein 2 | 1 |
| Q8WUH6 | TMEM263 | Transmembrane protein 263 | 1 |
| Q8WWC4 | MAIP1 | m-AAA protease-interacting protein 1, mitochondrial | 1 |
| Q92616 | GCN1 | Stalled ribosome sensor GCN1 | 1 |
| Q92835 | INPP5D | Phosphatidylinositol 3,4,5-trisphosphate 5-phosphatase 1 | 1 |
| Q92974 | ARHGEF2 | Rho guanine nucleotide exchange factor 2 | 1 |

| | | | |
|---|---|---|---|
| Q93084 | ATP2A3 | Sarcoplasmic/endoplasmic reticulum calcium ATPase 3 | 1 |
| Q969L2 | MAL2 | Protein MAL2 | 1 |
| Q969Z0 | TBRG4 | FAST kinase domain-containing protein 4 | 1 |
| Q96AG4 | LRRC59 | Leucine-rich repeat-containing protein 59 | 1 |
| Q96B97 | SH3KBP1 | SH3 domain-containing kinase-binding protein 1 | 1 |
| Q96BM9 | ARL8A | ADP-ribosylation factor-like protein 8A | 1 |
| Q96C19 | EFHD2 | EF-hand domain-containing protein D2 | 1 |
| Q96DA0 | ZG16B | Pancreatic adenocarcinoma up-regulated factor | 1 |
| Q96GC5 | MRPL48 | Large ribosomal subunit protein mL48 | 1 |
| Q96I25 | RBM17 | Splicing factor 45 | 1 |
| Q96IZ7 | RSRC1 | Serine/Arginine-related protein 53 | 1 |
| Q96N66 | MBOAT7 | Lysophospholipid acyltransferase 7 | 1 |
| Q99829 | CPNE1 | Copine-1 | 1 |
| Q99873 | PRMT1 | Protein arginine N-methyltransferase 1 | 1 |
| Q9BRU9 | UTP23 | rRNA-processing protein UTP23 homolog | 1 |
| Q9BSH4 | TACO1 | Translational activator of cytochrome c oxidase 1 | 1 |
| Q9BT23 | LIMD2 | LIM domain-containing protein 2 | 1 |
| Q9BU61 | NDUFAF3 | NADH dehydrogenase [ubiquinone] 1 alpha subcomplex assembly factor 3 | 1 |
| Q9BV79 | MECR | Enoyl-[acyl-carrier-protein] reductase, mitochondrial | 1 |
| Q9H0E2 | TOLLIP | Toll-interacting protein | 1 |
| Q9H1E1 | RNASE7 | Ribonuclease 7 | 1 |
| Q9H3K6 | BOLA2 | BolA-like protein 2 | 1 |
| Q9H4L5 | OSBPL3 | Oxysterol-binding protein-related protein 3 | 1 |

| | | | |
|---|---|---|---|
| Q9HAH7 | FBRS | Probable fibrosin-1 | 1 |
| Q9HB58 | SP110 | Sp110 nuclear body protein | 1 |
| Q9NP64 | ZCCHC17 | Zinc finger CCHC domain-containing protein 17 | 1 |
| Q9NQC3 | RTN4 | Reticulon-4 | 1 |
| Q9NRN7 | AASDHPPT | L-aminoadipate-semialdehyde dehydrogenase-phosphopantetheinyl transferase | 1 |
| Q9NUL3 | STAU2 | Double-stranded RNA-binding protein Staufen homolog 2 | 1 |
| Q9NVS2 | MRPS18A | Large ribosomal subunit protein mL66 | 1 |
| Q9NWU5 | MRPL22 | Large ribosomal subunit protein uL22m | 1 |
| Q9NX58 | LYAR | Cell growth-regulating nucleolar protein | 1 |
| Q9NY93 | DDX56 | Probable ATP-dependent RNA helicase DDX56 | 1 |
| Q9NZ08 | ERAP1 | Endoplasmic reticulum aminopeptidase 1 | 1 |
| Q9P2J5 | LARS1 | Leucine--tRNA ligase, cytoplasmic | 1 |
| Q9UBQ0 | VPS29 | Vacuolar protein sorting-associated protein 29 | 1 |
| Q9UBW5 | BIN2 | Bridging integrator 2 | 1 |
| Q9UGI8 | TES | Testin | 1 |
| Q9UGL9 | CRCT1 | Cysteine-rich C-terminal protein 1 | 1 |
| Q9UI10 | EIF2B4 | Translation initiation factor eIF2B subunit delta | 1 |
| Q9UNM6 | PSMD13 | 26S proteasome non-ATPase regulatory subunit 13 | 1 |
| Q9Y241 | HIGD1A | HIG1 domain family member 1A, mitochondrial | 1 |
| Q9Y2Q9 | MRPS28 | Small ribosomal subunit protein bS1m | 1 |
| Q9Y2S7 | POLDIP2 | Polymerase delta-interacting protein 2 | 1 |
| Q9Y2V2 | CARHSP1 | Calcium-regulated heat-stable protein 1 | 1 |
| Q9Y2Z0 | SUGT1 | Protein SGT1 homolog | 1 |

| | | | |
|---|---|---|---|
| Q9Y316 | MEMO1 | Protein MEMO1 | 1 |
| Q9Y3B7 | MRPL11 | Large ribosomal subunit protein uL11m | 1 |
| Q9Y4P3 | TBL2 | Transducin beta-like protein 2 | 1 |
| Q9Y5K6 | CD2AP | CD2-associated protein | 1 |
| Q9Y606 | PUS1 | Pseudouridylate synthase 1 homolog | 1 |

### Figures

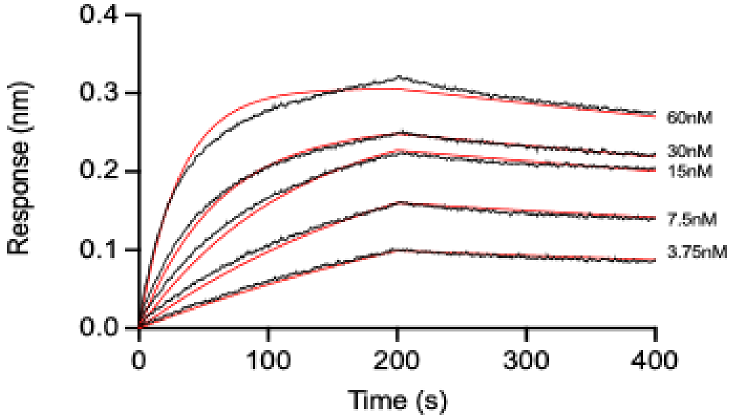


| $K_D$ (nM) | $K_{ON}$ ($10^5$ 1/Ms) | $K_{OFF}$ ($10^{-4}$ 1/Ms) |
|---|---|---|
| 1.1 ± 0.1 | 5.8 ± 0.6 | 6.1 ± 0.3 |

**Figure S1. S10A Fab binding to recombinant Siglec-10Fc.** Representative sensorgrams of S10A Fab binding to Siglec-10Fc (black) and the 1:1 complexes that provide the best fit for each concentration (red), calculated by BLI. The Table gathers the mean values of the kinetic parameters with the standard errors (SEM, representative of three independent measurements).

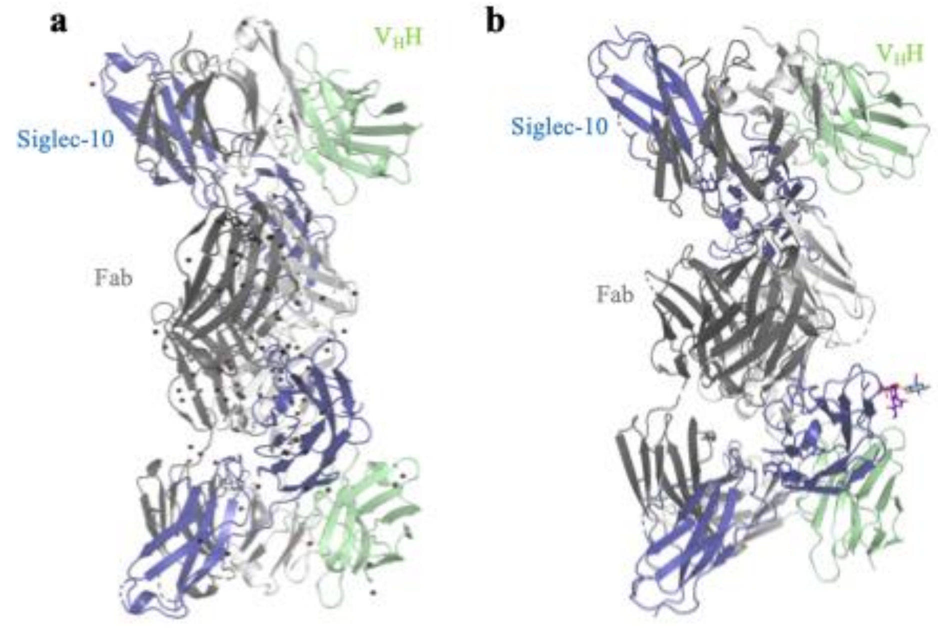


**Figure S2. Cartoon representation of the molecules present at the asymmetric unit of the crystals.** a) Cartoon representation of the two molecules of Siglec-10$d_1d_2$-S10A Fab-$V_HH$ complex found at the asymmetric unit of C2221 crystal. b) Cartoon representation of the two molecules of Siglec-10$d_1d_2$-S10A Fab-$V_HH$-6'SL complex at the asymmetric unit of the crystal with space group C2221. The 6'SL ligand is represented as sticks. Siglec-10 in blue, S10A HC in dark grey, S10A light chain in light grey and anti-kappa $V_HH$ in green.

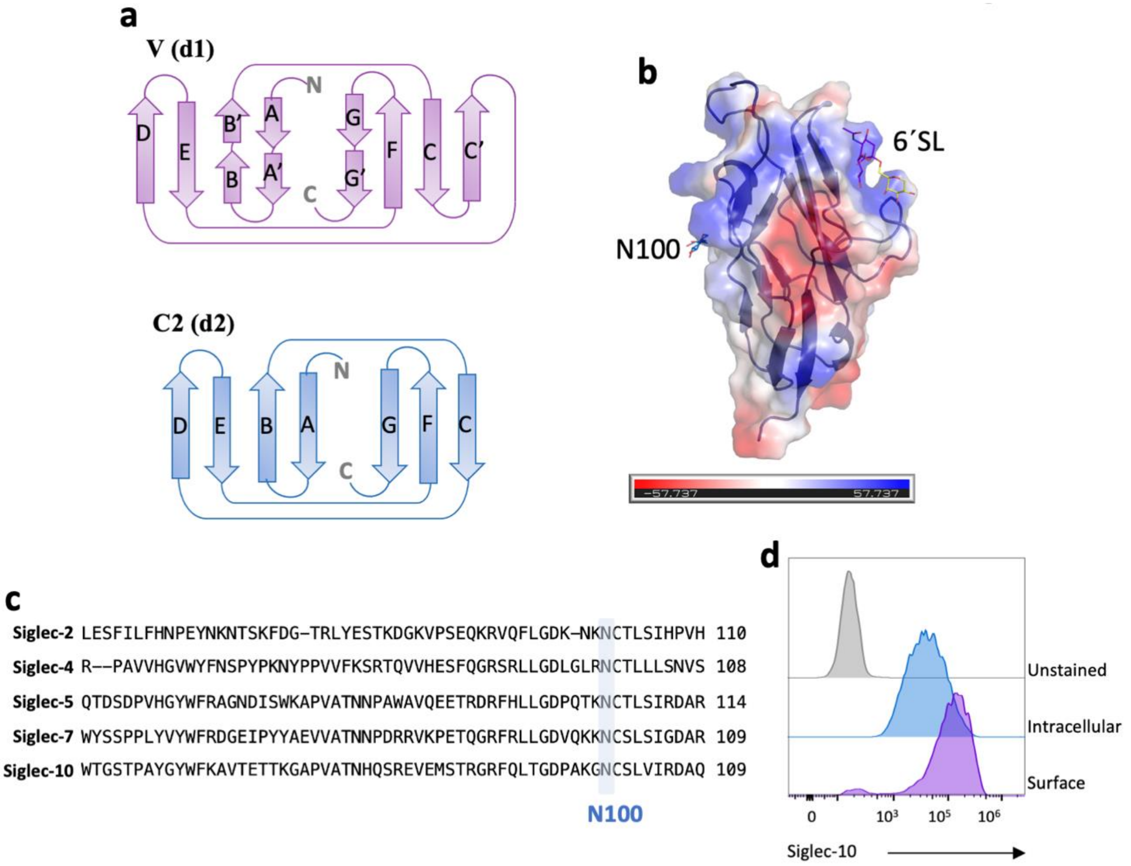


**Figure S3. Three-dimensional structure of human Siglec-10 and N100-linked glycans.** a) Schematic diagram of the V and C2 type Ig domains found in the crystal structure of Siglec-10. b) Electrostatic surface representation of the V-Ig domain of Siglec-10. The calculation of the surface electrostatics was made with the APBS software,[1] prepared using PyMOL[2] and is displayed on a scale of −5 kT/e (*red*) to 5 kT/e (*blue*). c) Sequence alignment of the V-Ig domain of human Siglec-10 and other members of the Siglec family (Siglec-2, -4, -5 and -7). The conserved N glycosylation site at N100 is highlighted in blue. d) Representative flow cytometry staining histograms indicating that intracellular and surface expression levels of Siglec-10 are not affected by the point mutation at N100 amino acid residue.

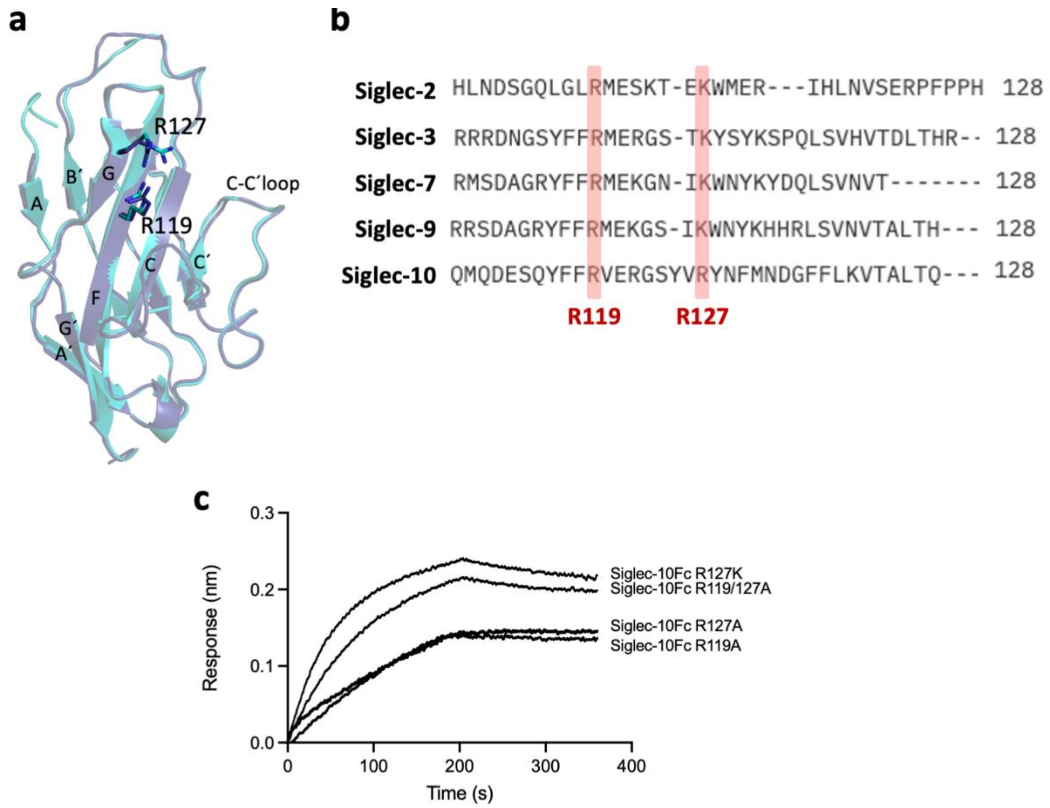


**Figure S4. Sequence comparison of V-Ig domains of distinct Siglecs.** a) Superposition of the crystal structures of the V-Ig domain of Siglec-10 in the apo form (dark blue) and in complex with 6´SL (cyan). b) Sequence alignment of residues 109–128 of Siglec-10 within the V-Ig domain and the corresponding regions of Siglecs-2, -3, -7, and -9. The canonical Arg residue is shown in red (R119 in Siglec-10), whereas R127 (in red) in Siglec-10 is substituted by a K residue in Siglecs-2, -3, -7, and -9. c) Representative sensorgrams showing S10A Fab binding to all mutant versions of recombinant Siglec-10Fc, as measured by BLI.

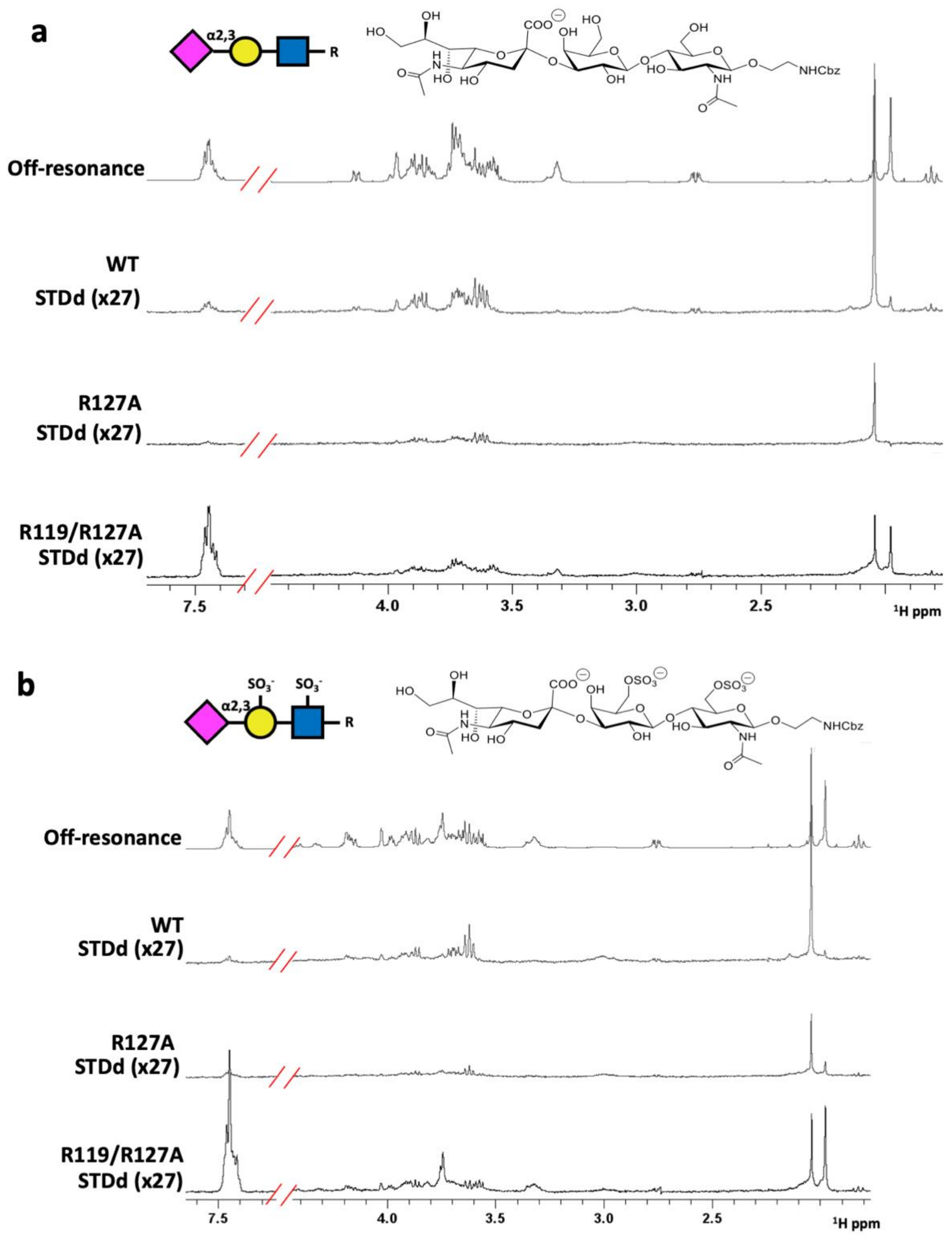


**Figure S5. STD-NMR profiles for Siglec-10 binding to 3'SLN and 6-*S*-6'-*S*-3'SLN.** a) Siglec-10Fc STD-NMR profile with 3'SLN. Mutation at the R127 position globally reduces the STD response, whereas R119A/R127A mutation completely abolishes the binding. b) Siglec-10Fc STD-NMR profile with 6-*S*-6'-*S*-3'SLN. Same observation as with 3'SLN.

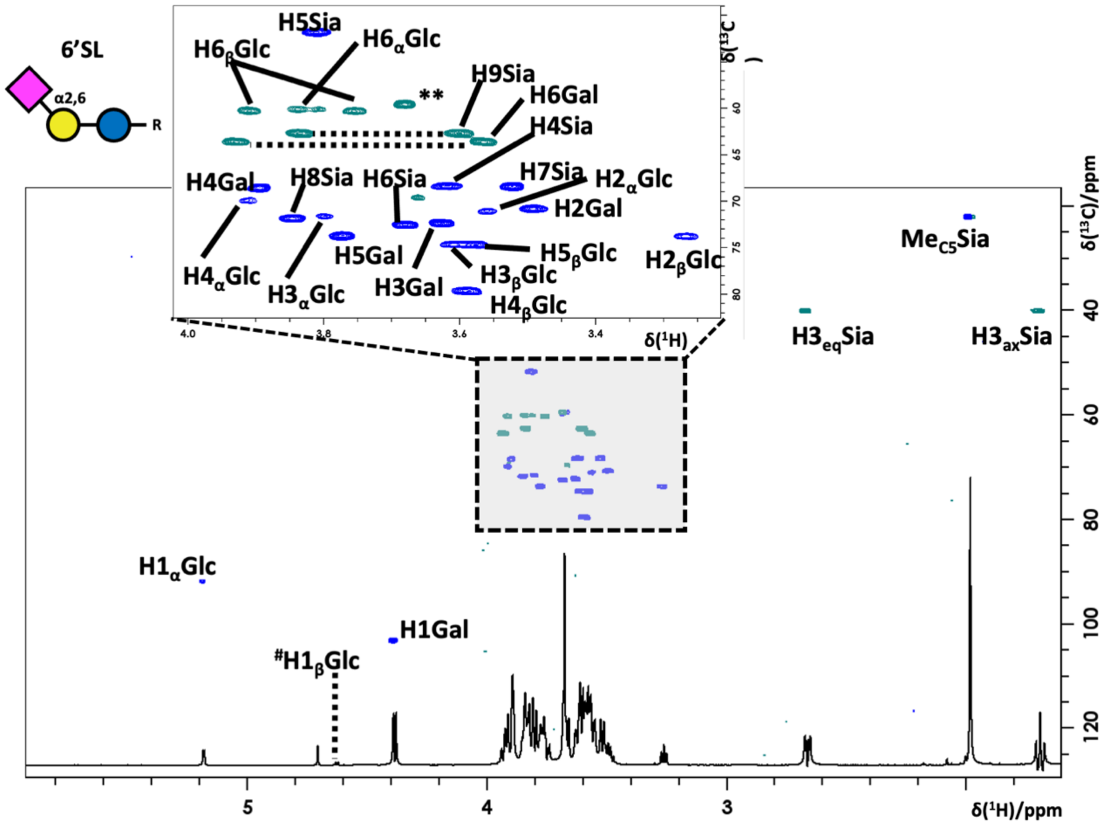


**Figure S6. $^{1}$H,$^{13}$C-HSQC spectra of 6'SL for ligand characterization.** $^{1}$H,$^{13}$C-HSQC spectra were recorded at 288K (500 MHz) in phosphate-buffered saline containing 10 mM, 300 mM NaCl, 0.1% NaN3, pH 7.4 in (100% $D_2O$), with their corresponding assignment.

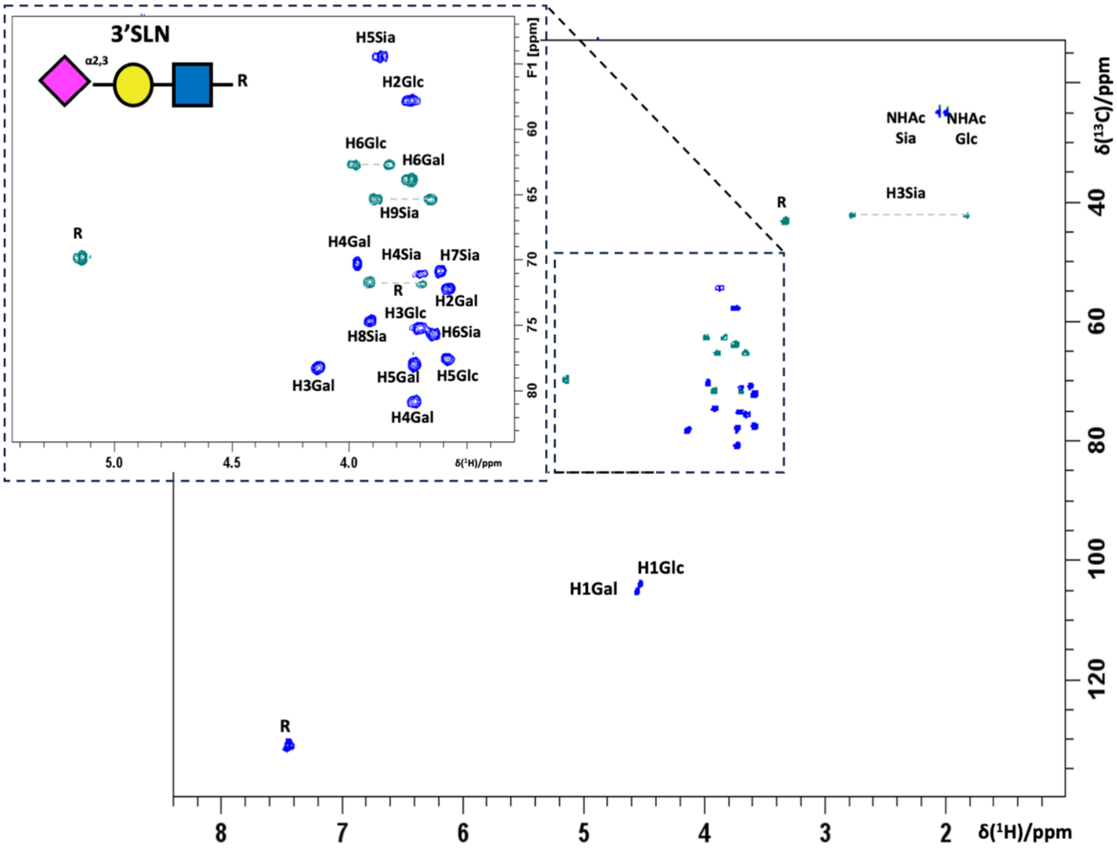


**Figure S7. ¹H,¹³C-HSQC spectra of 3’SLN.** ¹H,¹³C-HSQC spectra of 3’SLN at 288 K (500 MHz) in phosphate-buffered saline containing 10 mM, 300 mM NaCl, 0.1% NaN3, pH 7.4 in (100% $D_2O$), with their corresponding assignment.

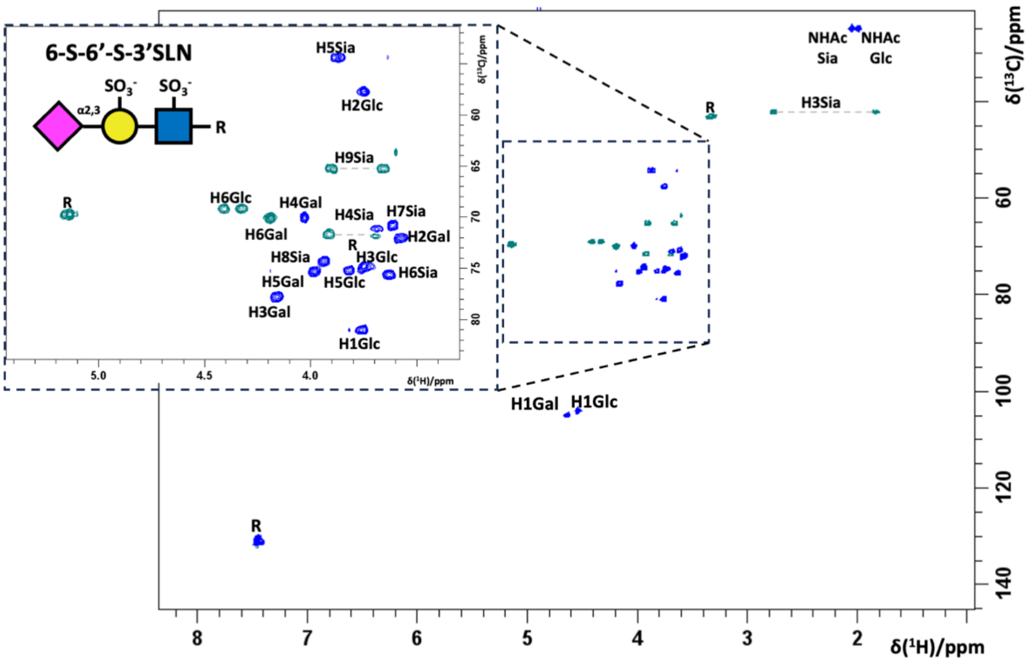


**Figure S8. ¹H,¹³C-HSQC spectra of the sulfated molecule 6-S-6'-S-3'SLN.** ¹H,¹³C-HSQC spectra of 6-S-6'-S-3'SLN at 288K (500 MHz) in phosphate-buffered saline containing 10 mM, 300 mM NaCl, 0.1% NaN3, pH 7.4 in (100% D2O), with their corresponding assignment.

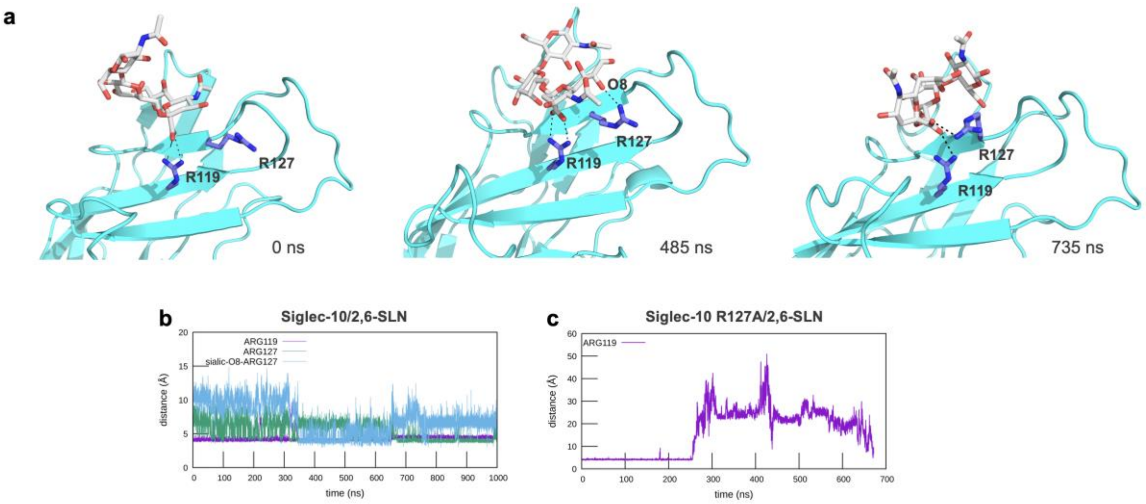


**Figure S9. Molecular dynamics (MD) simulations**. a) Snapshots from three different time points (220 and 232 ns) of a 1 μs MD simulation of Siglec-10 in complex with the 2,6-SL ligand, highlighting key interactions between the side chains of R119 and R127 (blue) and the sialic acid (shown as sticks, colored by element). Time evolution of the distances between the sialic acid and residues R119 and R127 over three independent simulations: b) WT Siglec-10 with 2,6-SL. c) Siglec-10 R127A mutant with 2,6-SL. In each plot, distances are shown between the selected atoms of R119 (CZ) and R127 (CZ) and the sialic acid (C1) as a function of simulation time. In the case of the WT Siglec-10 complex, time evolution of the distances between 2,6-SL and residues N129 (ND2) and T67 (OG1) are also shown.

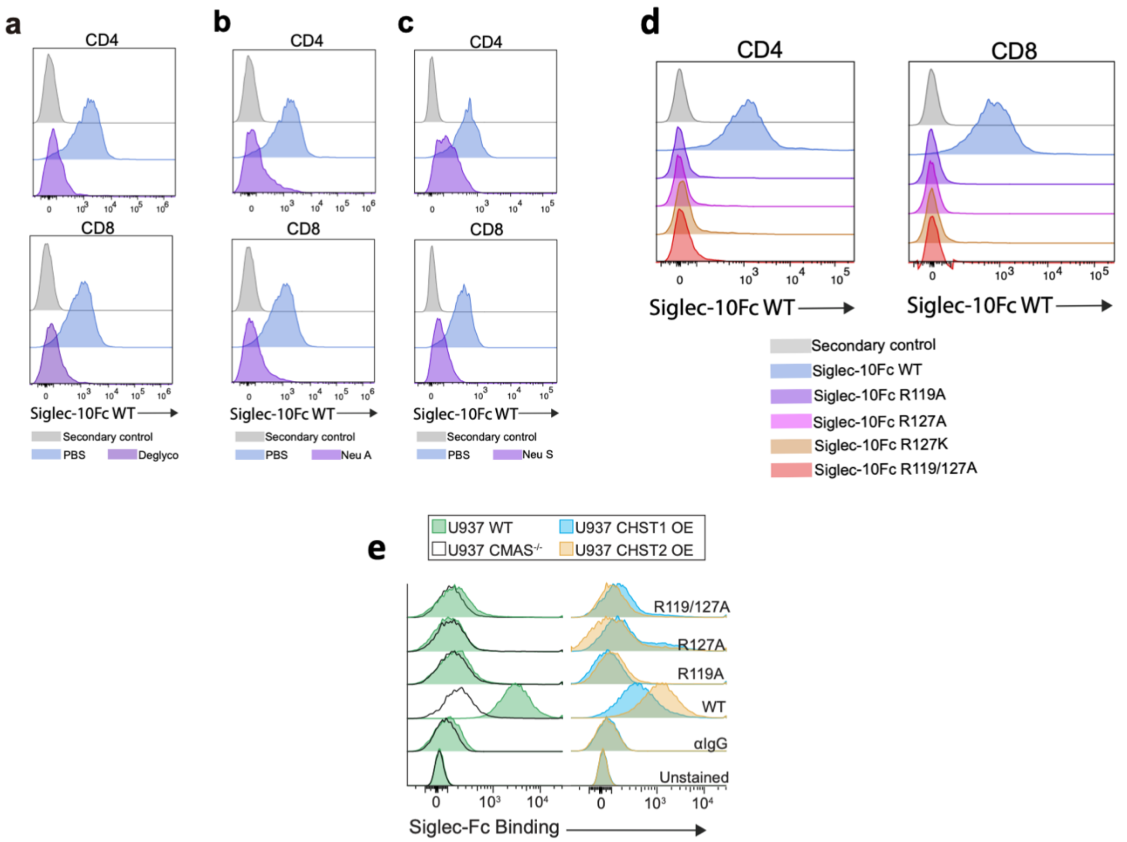


**Figure S10. Representative data of Siglec-10 binding to human cells.** a) Siglec-10Fc WT binding to deglycosylated CD4+ and CD8+ T cells. b) Siglec-10Fc WT binding to neuraminidase A treated CD4+ and CD8+ T cells. c) Siglec-10Fc WT binding to neuraminidase S treated CD4+ and CD8+ T cells. d) Binding of each of Siglec-10Fc mutants (R119A, R127A, R127K, and R119/127K) to T cells in comparison to Siglec-10Fc WT. e) Siglec-10Fc (WT, R119A, R127A and R119/127A) binding to U937 cell line. Left: Siglec-10Fc binding to parental cell line in comparison to U937 CMAS KO, representing that the sialic acid moiety is the driving force for the interaction. Right: Siglec-10Fc binding to parental U937 in comparison to CHST1 and CHST2 OE cells, suggesting that sulfation has no impact on Siglec-10 binding.

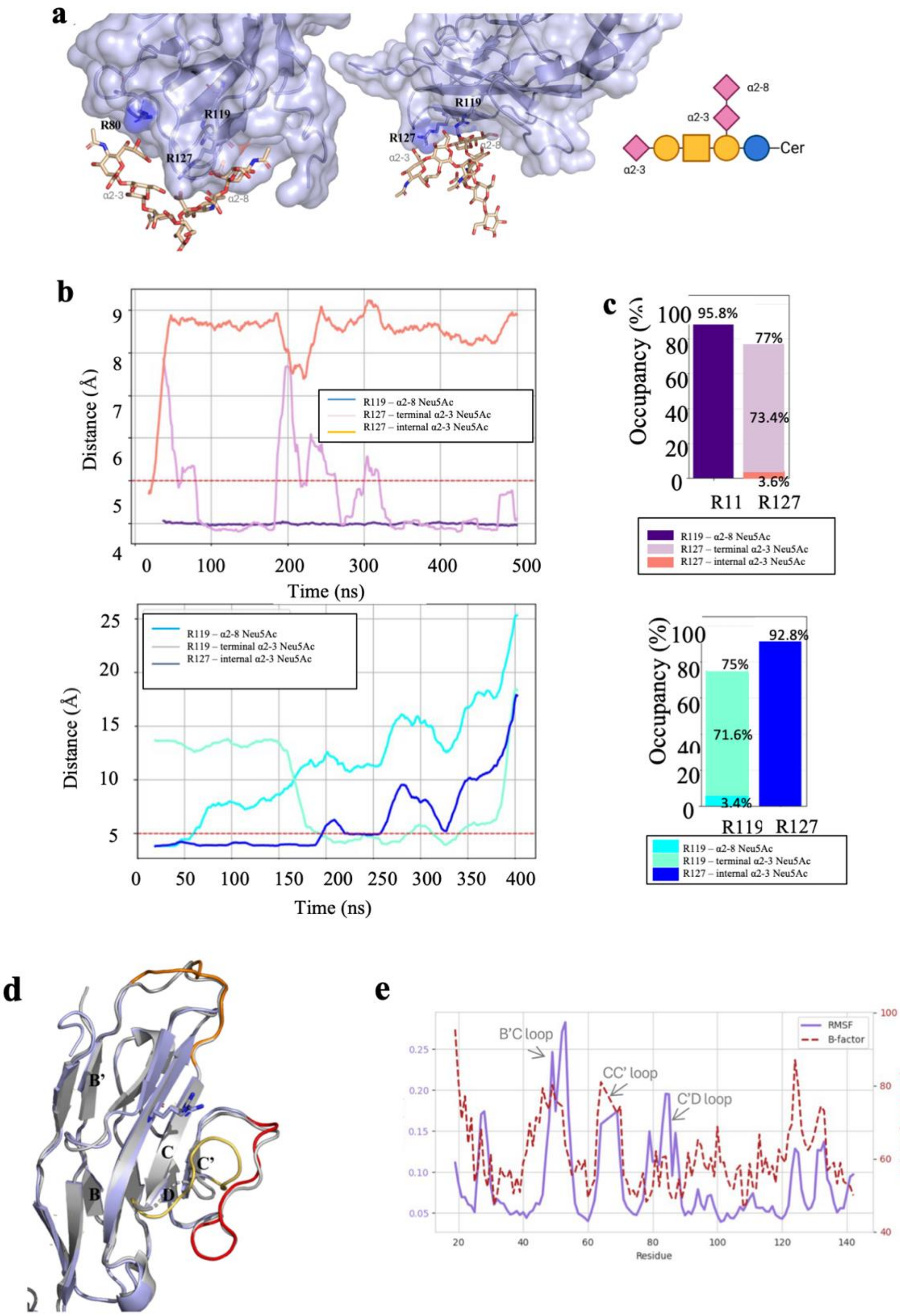


**Figure S11. GT1b ganglioside in complex with Siglec-10.** a) Left panel: 3D model of the Siglec-10/GT1b complex used for the MD simulations. The protein was rendered with transparent surface and cartoon in light blue to highlight secondary structure motifs. The GT1b ganglioside rendered with sticks with C atoms in tan, O in red, and N in blue. The three Neu5Ac are labelled according to their linkages (α2-3 terminal, α2-3 internal, and α2-8). Right panel: Representative frame from the MD simulation showing a rearranged binding conformation. The R127 shift its interaction to the terminal α2-3 Neu5Ac, while R119 maintains its interaction with the α2-8 Neu5Ac. b) Distance plots showing the interactions between Siglec-10 arginine residues and the Neu5Ac moieties of GT1b over 500 ns (MD1) (top) and 400 ns (MD2) (bottom). The red dashed line at 5 Å indicates the threshold distance for interaction. c) Occupancy analysis for the interactions between Arg and Neu5Ac residues in MD1 (top) and MD2 (bottom), respectively. Occupancy is defined as the percentage of simulation frames where the interatomic distance remains below 5 Å. MD1 shows dominant occupancy for R119 α2-8 (95.8%) and R127 α2-3 terminal (77.0%). In MD2, R119 switches preference, with 71.6% occupancy toward α2-3 terminal, and R127 shows a strong preference for α2-3 internal (92.8%). d) Structural alignment and RMSF analysis of the Siglec-10 V-set domain. *Left:* Cartoon representation of the V-set domain of Siglec-10 (AF-Q96LC7) is shown in light blue, aligned with the X-ray structure (this work) shown in grey. The B′C, CC′, and C′D loops are highlighted in orange, yellow and red, respectively. *Center*, SNFG representation of the GT1b ganglioside epitope, showing the three sialic acid arrangement: two α2-3-linked Neu5Ac (one terminal, one internal) and one α2-8-linked Neu5Ac bridging the internal sialic acid to galactose. e) Root mean square fluctuation (RMSF) from the MD simulation (purple line) compared with crystallographic B-factors (red dashed line) plotted as a function of residue number.

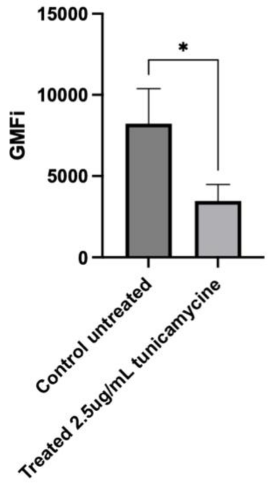


**Figure S12. Flow cytometry data of THP-1 cells stained with concanavalin A.** using streptavidin-APC (1: 2500) after treatment with tunycamycin. Mean of the untreated and treated with DMSO VS tunicamycine treatment.

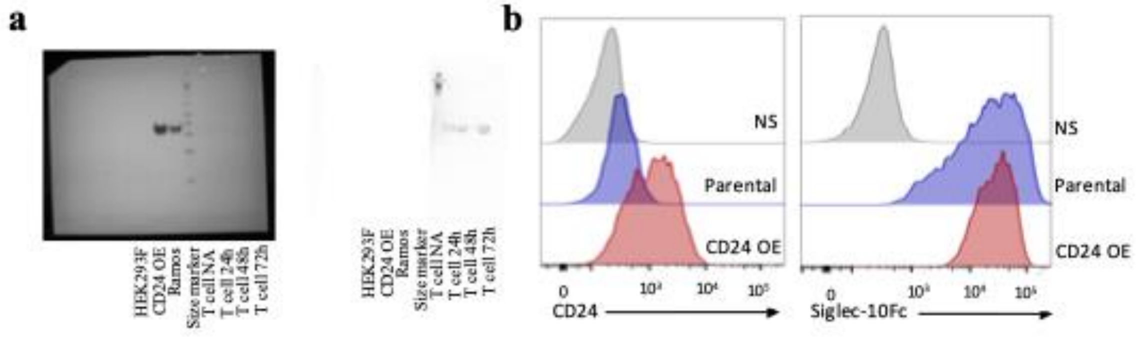


**Figure S13. Siglec-10 and CD24 interactions.** a) Whole Western Blot membranes which are corresponding to main text Figure 4a. The right membrane is partially covered to visualize the expression on T cells without overexposure of the CD24 expressed in control cell lines. b) Left – Representative histogram showing CD24 expression in the HEK293T CD24 overexpressing (OE) cell line generated in comparison to the parental cells. Right – Representation of Siglec-10 binding to the CD24 OE cells in comparison to the parental cell line.

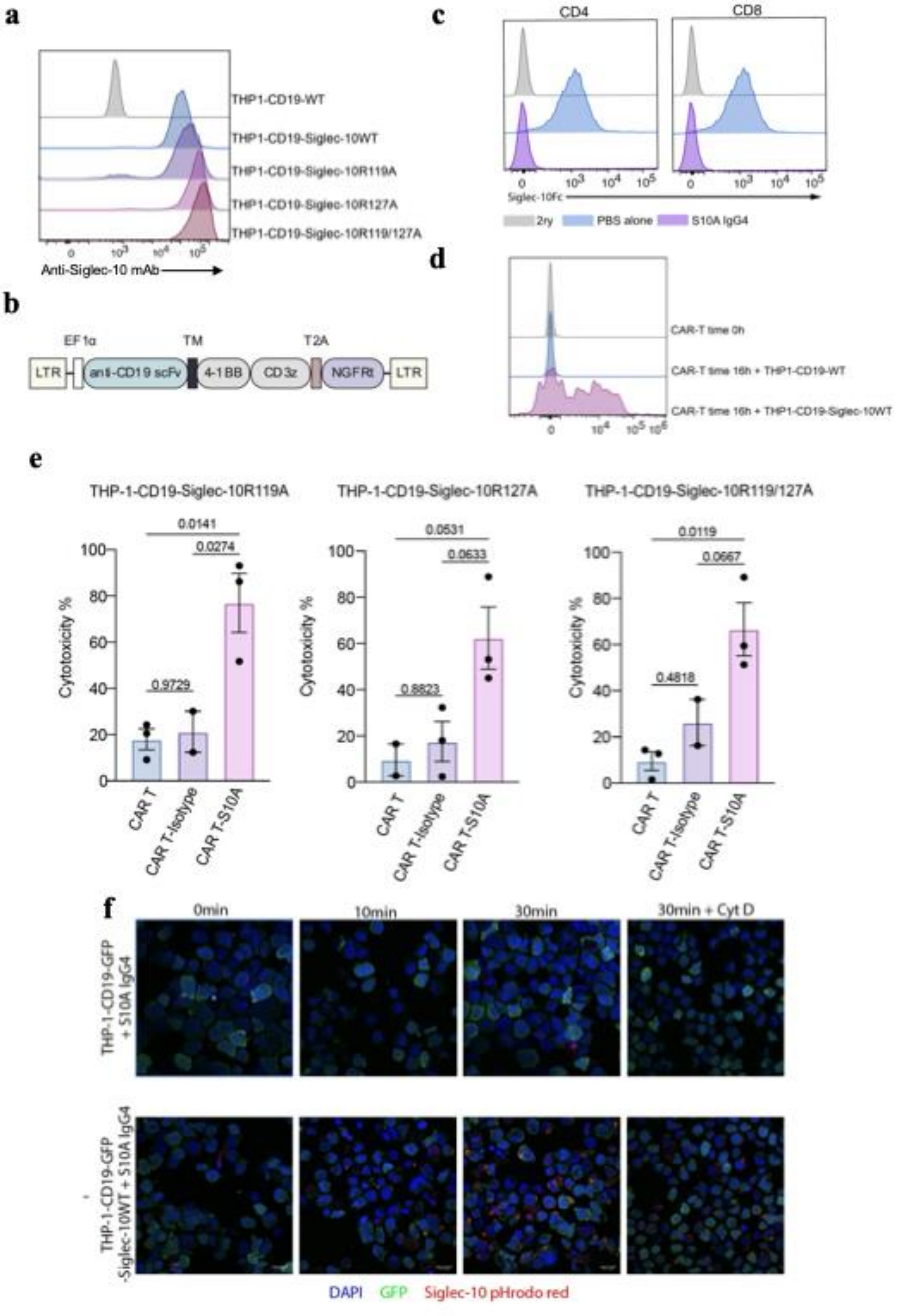


**Figure S14. The effect of Siglec-10 mutants on CAR-T cell function.** a) Representative histogram of Siglec-10 (WT, R119A, R127A, R119/127A and ITIM mut) overexpression on THP-1-CD19 cell line in comparison to the parental cell line. b) Schematic diagram representing the anti-CD19 CAR construct. Abbreviations: Long terminal repeat (LTR), Elongation factor 1 alpha (EF1α), FMC63 single chain variable fragment (anti-CD19 scFv), transmembrane domain (TM), tumor necrosis factor receptor superfamily member 9 (4-1BB), CD3 zeta chain (CD3z), self-cleaving 2A peptide sequence (T2A), truncated nerve growth factor receptor (NGFRt) c) Representative histograms illustrating S10A IgG4 blockade of Siglec-10 interactions on activated human CD4+ and CD8+ T cells. d) Representative histogram of Siglec-10 expression on CAR-T cells after 16h coculture with THP-1-CD19-Siglec-10WT overexpressing cells. e) Pooled data representing the impact of R119A, R127A and R119/127A mutations on the CAR-T cell function (n=3 donors, data represent mean ± SEM; one-way ANOVA). f) Confocal microscopy images depicting S10A IgG4 internalization in THP-1 cells.

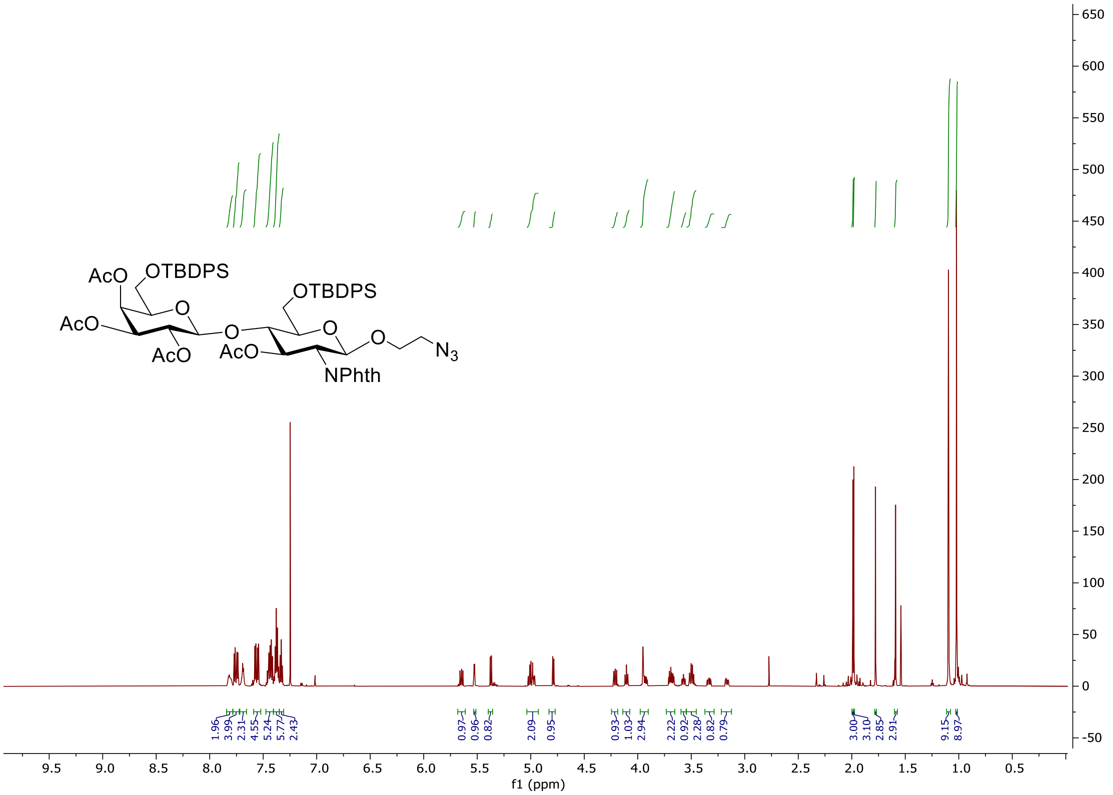

**Figure S15. [1]H NMR spectra of compound 3 of the 6-$SO_3$-6'-$SO_3$-3'SLN synthesis pathway.**

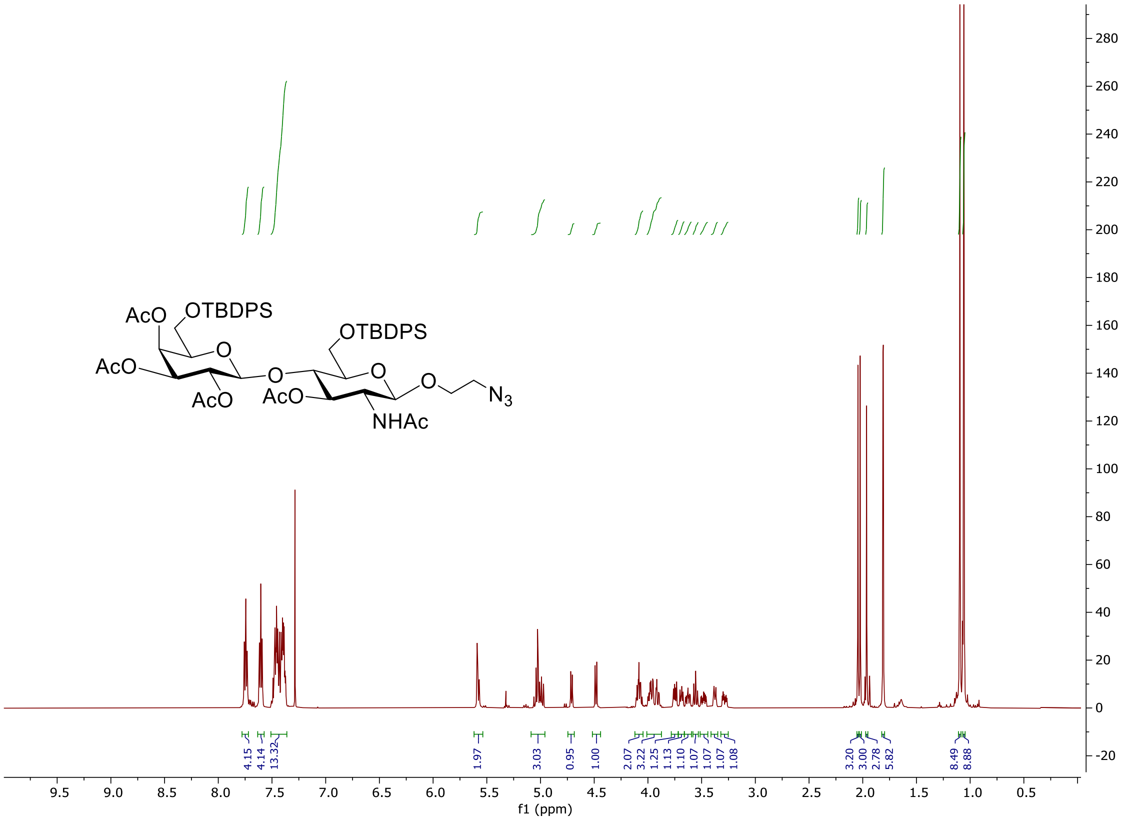


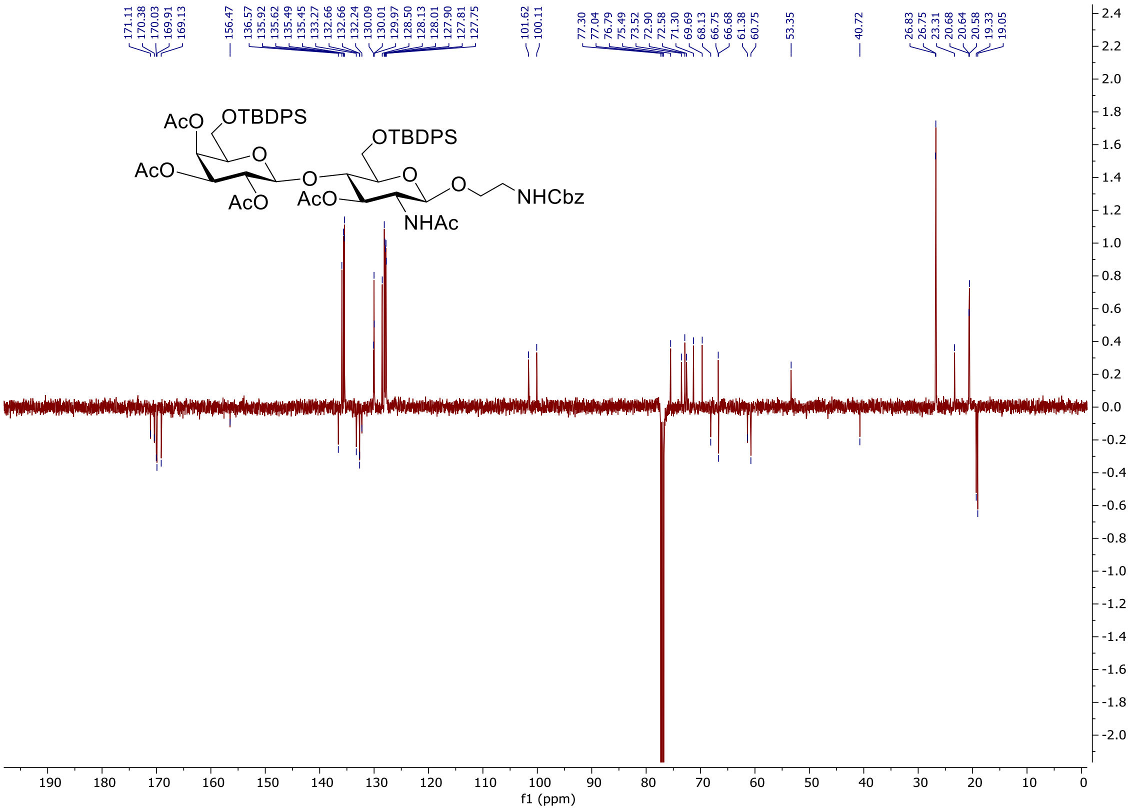


**Figure S16. NMR spectra of compound 4 of the 6-$SO_3$-6'-$SO_3$-3'SLN synthesis pathway.**

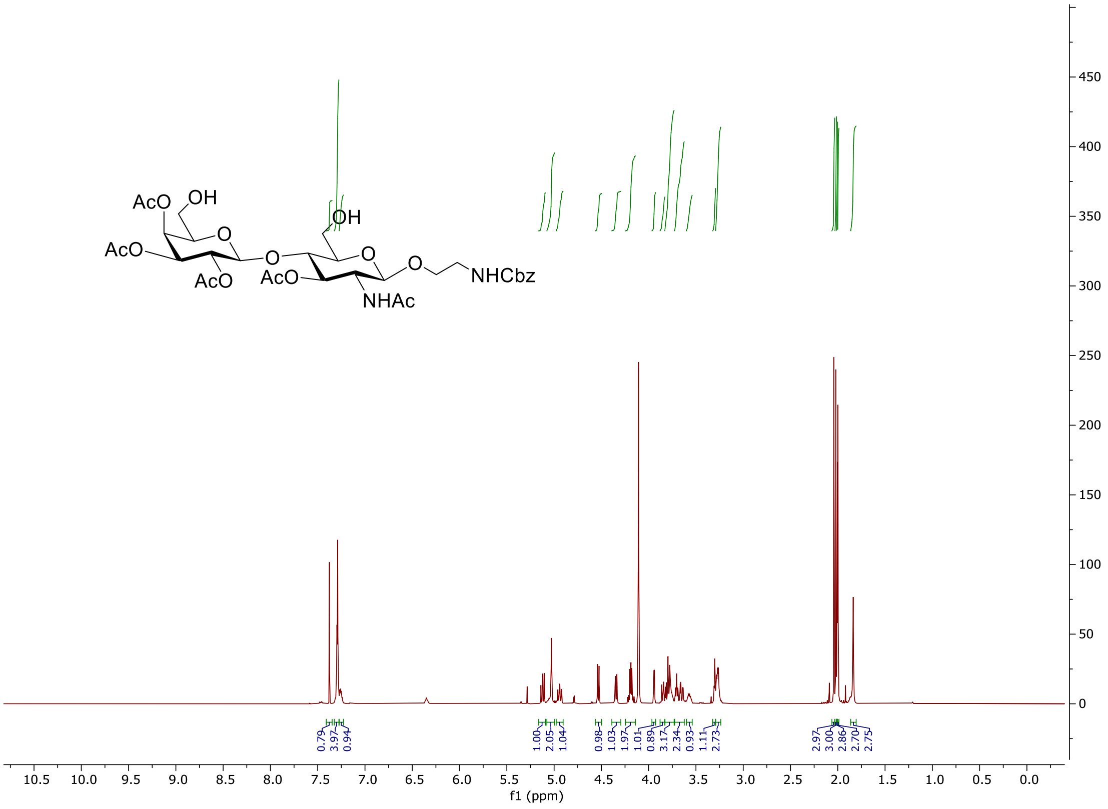

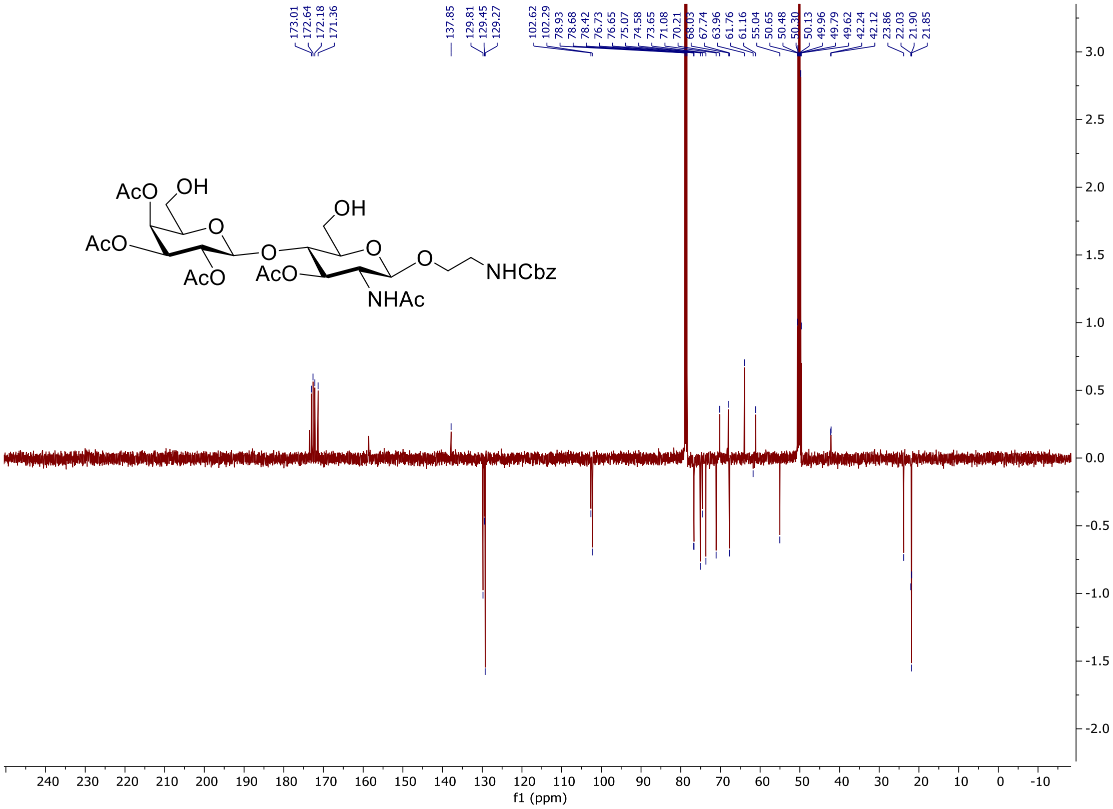

**Figure S17. NMR spectra of compound 5 of the 6-$SO_3$-6'-$SO_3$-3'SLN synthesis pathway.**

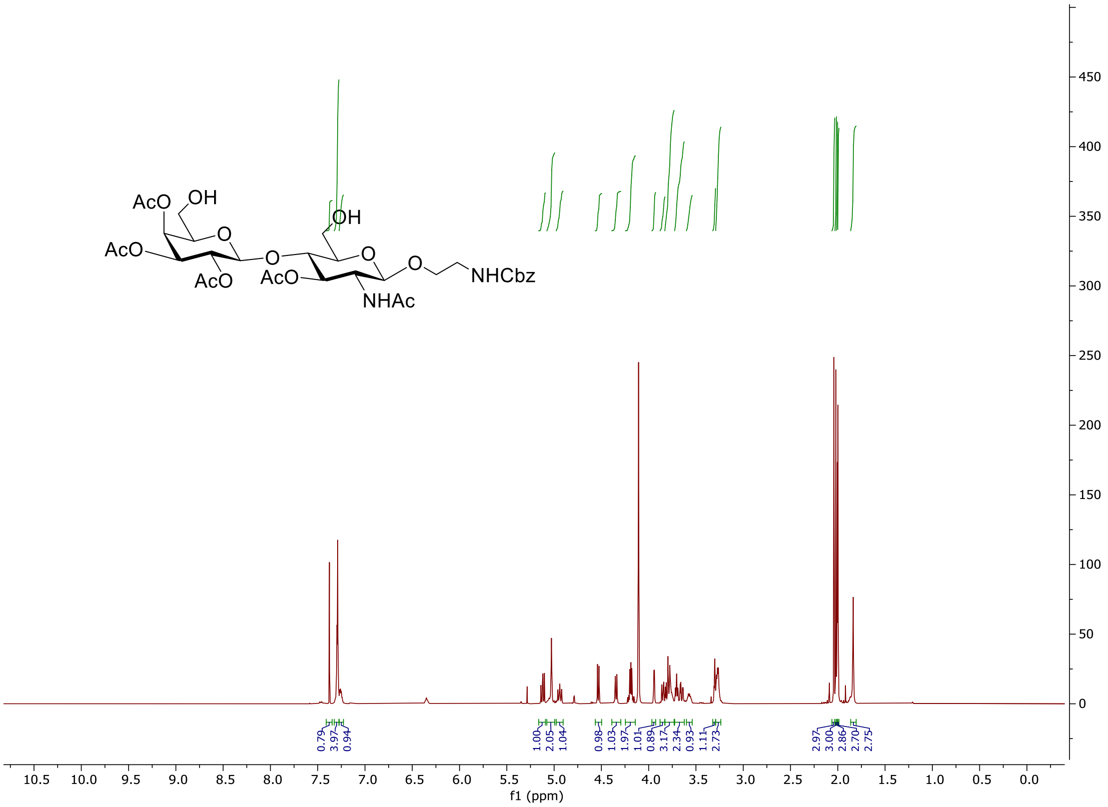

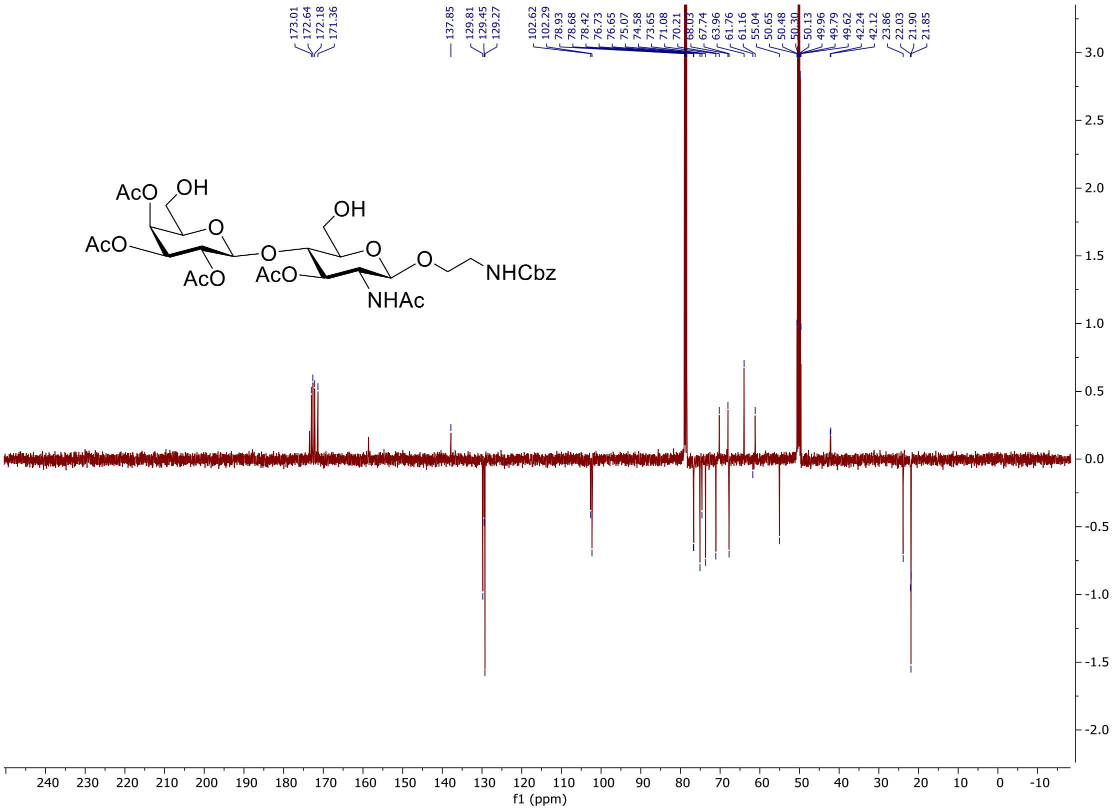

**Figure S18. NMR spectra of compound 6 of the 6-SO3-6’-SO3-3’SLN synthesis pathway.**

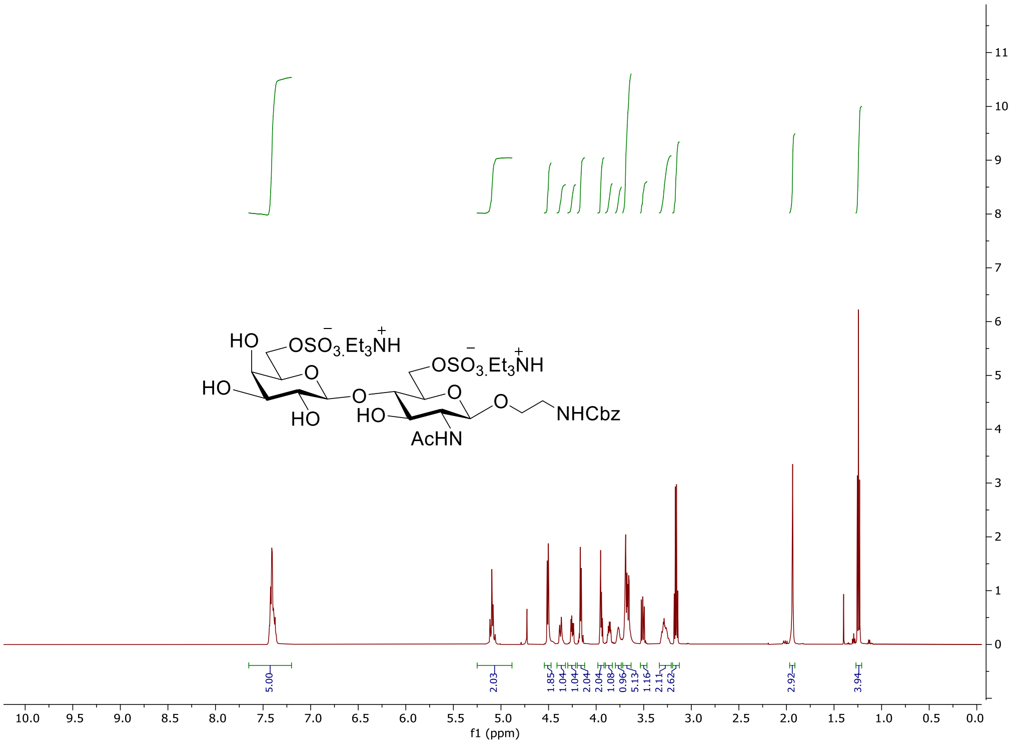


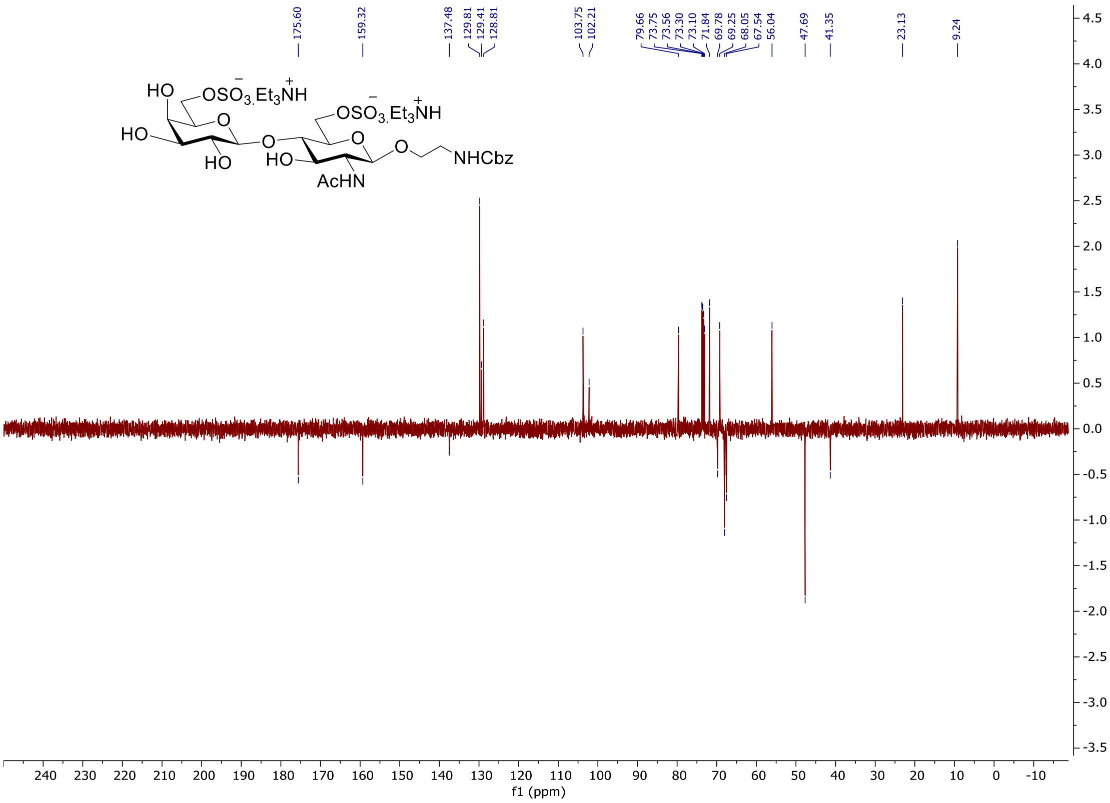


**Figure S19. NMR spectra of compound 7 of the 6-SO3-6’-SO3-3’SLN synthesis pathway.**

2. PyMOL | pymol.org. https://pymol.org/#page-top.
